# Age-Dependent Chromatin Remodeling Drives Inflammatory Dysregulation in Tendon Cells

**DOI:** 10.64898/2026.01.05.697725

**Authors:** Tyler E. Blanch, Ellen Y. Zhang, Seongwoo Kim, Bat-Ider Tumenbayar, Xi Jiang, Il Keun Kwon, Nathaniel A. Dyment, Inkyung Jung, Su Chin Heo

## Abstract

Aging impairs tissue function and tolerance to cellular stress by reprogramming the behavior of resident cells. With global increases in lifespan, the prevalence of chronic and degenerative musculoskeletal disorders, including tendon degeneration, continues to rise; however, effective interventions to counteract age-related decline remain limited. Here, we investigate how a central age-associated stressor, inflammation, differentially modulates tendon cell behavior derived from young and mature-aged donors. Using super-resolution microscopy to resolve nanoscale chromatin organization in conjunction with epigenomic and transcriptomic profiling, we identify age-dependent regulatory mechanisms that govern inflammatory responsiveness. Mature-aged tendon cells exhibit exaggerated pro-inflammatory and catabolic responses across chromatin, gene expression, and protein signaling levels, characterized by enhanced TNFα receptor organization, elevated accessibility at pro-inflammatory regulatory elements, and robust induction of matrix-degrading enzymes. Notably, the AP-1 transcription factor family emerges as a central age-dependent regulator, displaying distinct motif accessibility patterns that bias mature tenocytes toward inflammatory and degenerative transcriptional programs. Taken together, our findings demonstrate that age-dependent epigenetic priming amplifies inflammatory sensitivity and constrains reparative gene regulation in mature tendon cells. This work provides a mechanistic framework linking chromatin remodeling to tendon degeneration and holds potential to identify epigenetic and transcriptional pathways as potential targets for rejuvenation strategies in aging musculoskeletal tissues.

## 1. Introduction

Tendons play a crucial role in musculoskeletal function by transmitting forces from muscle to bone and stabilizing joint motion ^1,2^. However, their intrinsically low cellularity and limited vascularity make them prone to injury and slow to heal ^3^, often leading to tendinopathy, partial tears, or chronic degeneration ^4^. The tendon healing process involves a sequence of overlapping phases: inflammation, proliferation, and remodeling ^5^. The inflammatory phase is initiated immediately after injury, characterized by the release of pro-inflammatory cytokines such as tumor necrosis factor-α (TNFα), which promote clot formation, recruit immune cells, and initiate the early repair process ^5–7^. While transient inflammation is essential for effective tissue repair ^8,9^, chronic or dysregulated inflammation disrupts extracellular matrix (ECM) homeostasis, promotes prolonged matrix degradation, and drives progressive cell- and tissue-level dysfunction ^10–12^. This maladaptive inflammatory state is particularly prevalent in aging tissues, where impaired ECM homeostasis often contributes to structural failure and acute loss of mechanical integrity ^13,14^.

With maturation and aging, tendons undergo profound structural, biomechanical, and cellular alterations, including increased stiffness, diminished elasticity, and changes in tenocyte phenotype ^1,15^. These age-associated alterations compromise load-bearing capacity, delay healing, and predispose tendons to degeneration ^12,15,16^. A hallmark of aged tendon tissue is chronic low-grade inflammation, which accelerates local ECM breakdown and destabilizes tissue homeostasis, potentially reinforcing a cycle of inflammation and degeneration mediated by altered cell-matrix signaling ^4,10,12^. Recent research further demonstrates that tendon development and aging not only delay healing but fundamentally alter the injury response at the cellular level ^17,18^. Despite growing insight into tendon biology, the cellular and molecular mechanisms underlying age-related differences in tendon healing remain inadequately understood. In particular, how tendon cells (i.e., tenocytes) of different maturational states respond to inflammatory stimuli remains largely unexplored ^9,12,19^. Bridging this knowledge gap is crucial for identifying new therapeutic targets capable of restoring tissue homeostasis and mitigating chronic tendon degeneration in aging populations.

Chromatin organization is a fundamental regulator of gene expression, governing transcriptional output by modulating DNA accessibility to the transcriptional machinery ^20–23^. Epigenetic modifications, such as histone acetylation (e.g., acetylation of histone H3 at lysine 9, H3K9ac) and methylation (e.g., trimethylation of histone H3 at lysine 27, H3K27me3), dynamically remodel chromatin architecture and gene expression in response to environmental and biochemical cues, including inflammatory signals ^24–28^. These modifications regulate the balance between transcriptionally permissive euchromatin and repressive heterochromatin, thereby orchestrating gene activation and silencing programs ^29–31^. Despite growing recognition of the role of chromatin dynamics in cellular adaptation, the regulation of chromatin states during tendon development, aging, and inflammatory injury remains poorly defined. Importantly, studying chromatin accessibility provides a direct readout of how injury-associated stimuli, such as TNFα-induced inflammation, affect the transcriptional potential of key genes in tenocytes required for tissue homeostasis and repair. Therefore, age-dependent differences in chromatin accessibility may represent a key molecular determinant of the divergent healing capacities observed between young and mature tendons ^18,32^. Understanding these epigenetic mechanisms could provide a foundation for targeted interventions to restore tissue homeostasis and mitigate tendon degeneration.

Thus, to address these gaps, we investigate age-dependent gene expression and chromatin remodeling in tenocytes under inflammatory conditions, examining how these changes influence cellular function. Using advanced approaches such as super-resolution imaging and epigenetic profiling ^22^, we explore how age-dependent chromatin remodeling governs tenocyte inflammatory responsiveness and functional output. We demonstrate that maturation and aging fundamentally reprogram the chromatin landscape and inflammatory signaling capacity of tenocytes. Mature tenocyte gene expression patterns exhibit disrupted ECM homeostasis following TNFα stimulation ^9^, accompanied by enhanced TNFα receptor clustering and amplified inflammatory signal propagation. Super-resolution imaging and ATAC-seq pathway analysis further reveal pronounced baseline differences in genome organization and chromatin accessibility between young and mature tenocytes that are further exacerbated under inflammatory challenge, resulting in a marked expansion of heterochromatin domains. Motifs accessibility analysis further identifies age-dependent enrichment of transcription factor families including AP-1, RFX, and IRF, implicating chromatin-mediated amplification of transcriptional networks with inflammation and aging ^27^.

By focusing on age-dependent changes in chromatin accessibility and their functional consequences in tenocytes, this study reveals epigenetic mechanisms that couple aging to degenerative tendon pathology. These findings establish a new mechanistic framework for targeting chromatin-based pathways and developing therapeutic interventions to restore tenocyte function, rebalance inflammatory responses, and improve regenerative outcomes in aging tendons and chronic tendinopathy.

## 2. Results

### 2.1 Age-Dependent Chromatin Remodeling and Epigenetic Plasticity in Tenocytes Under Inflammatory Stress

Given the central role of chromatin accessibility and organization in regulating transcription factor activity and gene expression ^23,33,34^, and previous evidence linking inflammatory signaling to epigenetic remodeling ^35–37^, we first sought to identify inflammation-dependent mechanisms underlying tenocyte chromatin conformation and epigenetic dysfunction. To do this, primary tenocytes were isolated from tail tendons from young (4 weeks old) or mature (45-50 weeks old) mice, and to mimic the initial acute inflammatory response observed in tendon injuries ^9^, cells were treated with inflammatory cues (i.e., TNFα) for up to 6 hours **(Fig. 1a)**.

**Figure 1:**
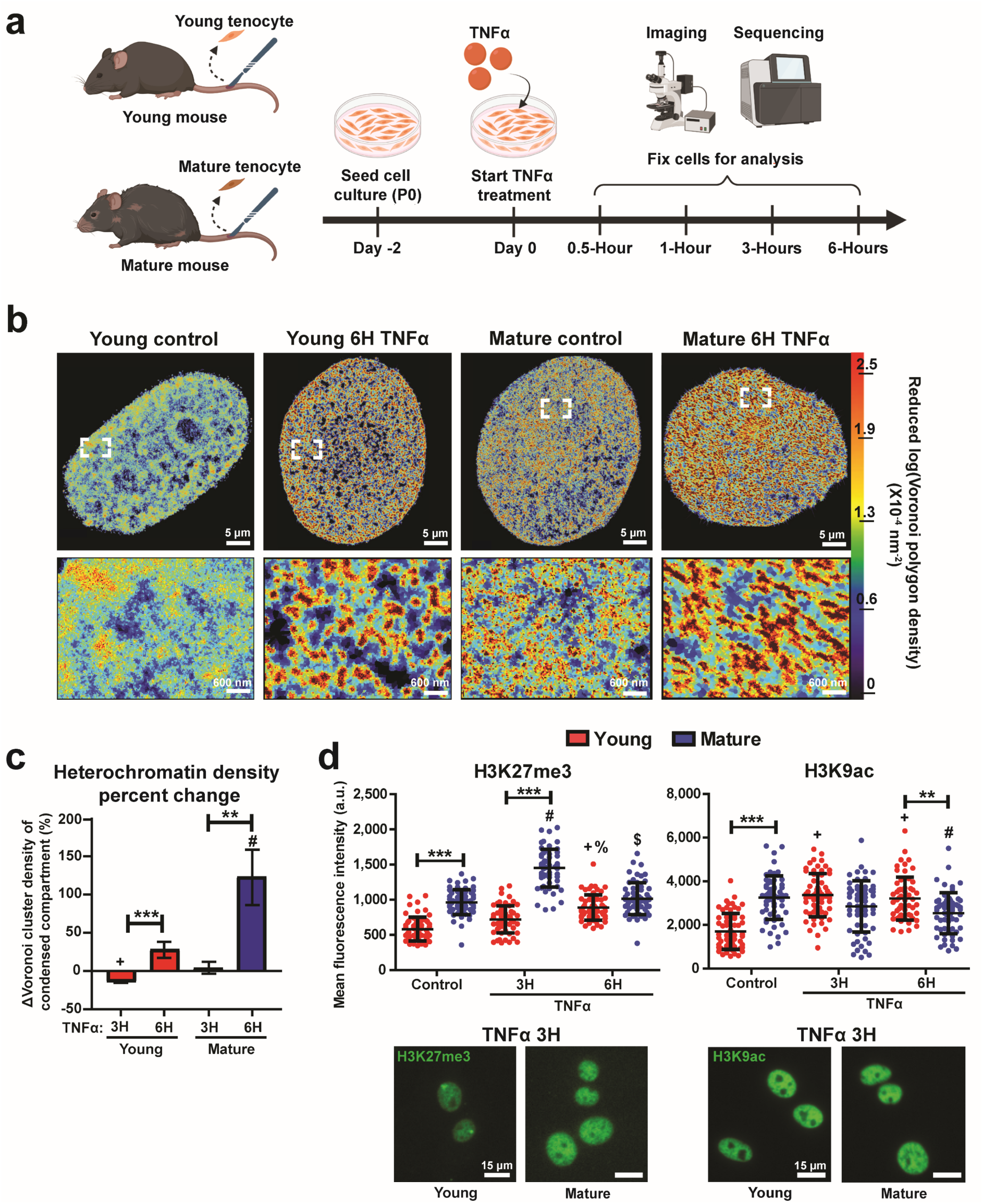
TNFα treatment promotes chromatin rearrangements and epigenetic alterations primarily in mature tenocytes. **(a)** Schematic showing isolation of young and mature mouse tail tenocytes and inflammatory stimulation with TNFα. **(b)** Representative H2B-STORM heat maps showing nanoscale H2B localization density within the nucleus of young and mature tenocytes under control and 6-hour TNFα treatment (6H). The lower panels show 8-fold magnified regions (white boxes), and color indicates nanoscale chromatin density determined by Voronoi tessellation. **(c)** Percentage change in the heterochromatin domain density following TNFα treatment in young (red) and mature (blue) tenocytes, normalized to respective control groups. (n=16 cells/group). **(d)** Immunofluorescence mean intensity quantification (top) and representative images (bottom) of histone modifications H3K27me3 (left) and H3K9ac (right) in young (red) and mature (blue) tenocytes. (n=60 cells/group). (Mean ± SD, +: p<0.05 vs Young Control, #: p<0.05 vs Mature Control, %: p<0.05 vs Young 3H, $: p<0.05 vs Mature 3H, **: p<0.01, ***: p<0.001).

To assess age-dependent chromatin architecture and remodeling under inflammatory conditions, we employed super-resolution imaging of nuclear histone H2B (i.e., stochastic optical reconstruction microscopy, H2B-STORM) to characterize nanoscale chromatin organization through Voronoi tessellation and density mapping of individual H2B proteins ^22,38^. Heatmaps of nanoscale chromatin domain density revealed subtle baseline differences in genome architecture between young and mature tenocytes, with mature cells exhibiting more densely packed and numerous heterochromatin domains **(Fig. 1b)**. Following six hours of TNFα treatment (6H TNFα), chromatin condensation became markedly more pronounced in mature cells compared to their young counterparts, even when accounting for baseline differences **(Fig. 1b, c and Supplemental Fig. S1a)**.

To further investigate the driving molecular mechanisms behind genomic rearrangements under inflammatory conditions, we examined histone modifications associated with gene repression (trimethylation of histone H3 at lysine 27, H3K27me3 ^39^) and activation (acetylation of histone H3 at lysine 9, H3K9ac, and trimethylation of histone H3 at lysine 4, H3K4me3 ^40^). TNFα treatment elevated H3K27me3 levels in both age groups, with a significantly greater elevation in mature cells **(Fig. 1d)**. In contrast, the activating modification H3K9ac levels decreased in mature cells but increased in young cells after six hours of TNFα stimulation **(Fig. 1d)**. The upregulation of H3K9ac observed in young cells may counterbalance the elevated H3K27me3 levels, resulting in a less severe chromatin condensation response compared to mature cells ^41^. No significant changes were observed in H3K4me3 levels in either age group upon TNFα treatment **(Supplemental Fig. S1b)**.

Collectively, these findings indicate that mature tenocytes undergo a global loss of active (open) euchromatic regions and increase the density of repressed (closed) heterochromatic domains under inflammatory conditions, resulting in a more condensed chromatin state. In contrast, young cells exhibit greater chromatin state plasticity, suggesting that maturation may diminish the adaptive potential for chromatin reorganization and increase susceptibility to inflammation-induced dysfunction ^42^.

### 2.2 Age-Dependent Alterations in Inflammatory Pathway Activation in Tenocytes

Given that chromatin remodeling under inflammation was dependent on donor age, we next investigated how inflammatory signaling differed between young and mature tenocytes. To confirm activation of the canonical TNFα-NFκB signaling pathway ^43^, we performed immunofluorescence (IF) staining for NFκB after various durations of TNFα treatment **(Fig. 2a)**. NFκB activation was quantified by measuring the nuclear-to-cytoplasmic fluorescence intensity ratio per cell ^43^. A robust nuclear translocation of NFκB was observed in both age groups, peaking at 0.5 hours post-treatment **(Fig. 2a, b)**. However, no significant differences in NFκB translocation rates were observed between young and mature tenocytes **(Fig. 2b)**, indicating that initial NFκB activation occurs similarly across ages. Cell viability and proliferation were monitored up to 48 hours after TNFα exposure, showing no significant decline in either parameter compared to control conditions **(Supplemental Fig. S2a, b)**. Interestingly, TNFα enhanced cell proliferation in mature cells **(Supplemental Fig. S2b)**. Morphometric analysis revealed that mature cells exhibited larger cell spread areas and higher nuclear localization of yes-associated protein 1 (YAP) under baseline conditions. Upon TNFα stimulation, YAP nuclear accumulation increased only in young cells, while cell size remained unchanged in both groups **(Fig. 2a, c and Supplemental Fig. S2c)**.

**Figure 2:**
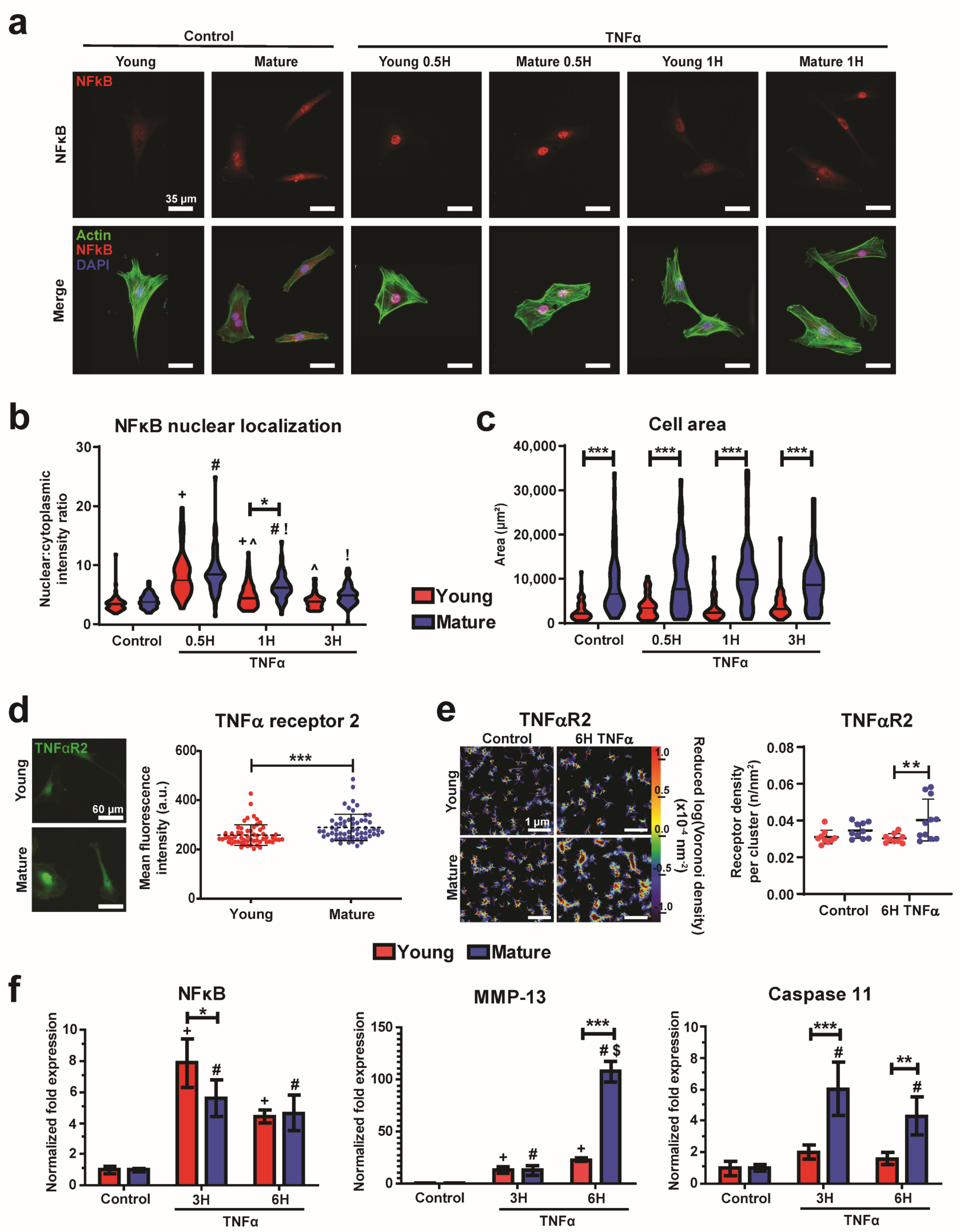
Inflammatory cues elicit age-dependent gene expression changes in mouse tenocytes. **(a)** Representative immunofluorescence (IF) images of young and mature tenocytes stained for NFκB (red), f-actin (green), and DNA (blue) at control or TNFα-treated conditions. **(b)** Quantification of the nuclear-to-cytoplasmic fluorescence intensity ratio of NFκB in young (red) and mature (blue) tenocytes following TNFα treatment (n=60 cells/group). **(c)** Cell area of young (red) and mature (blue) tenocytes following TNFα treatment (n=60 cells/group). **(d)** Representative IF images (left) and mean intensity quantification (right) for TNFα Receptor 2 (TNFαR2) in young and mature tenocytes under control conditions (n=60 cells/group). **(e)** Representative STORM density heatmaps (left) and quantification of nanoscale receptor cluster density (right) for TNFαR2 in young (red) and mature (blue) tenocytes under control and TNFα-treated conditions (n=11 cells/group, 3 ROIs per cell). **(f)** Fold-change in gene expression of NFκB (left), MMP-13 (middle), and Caspase 11 (right) in young (red) and mature (blue) tenocytes after TNFα treatment, normalized to control expression levels (n=3 technical replicates, normalized to GAPDH expression). (Mean ± SD, +: p<0.05 vs Young Control, #: p<0.05 vs Mature Control, ^: p<0.05 vs Young 0.5H, !: p<0.05 vs Mature 0.5H, $: p<0.05 vs Mature 3H, *: p<0.05, **: p<0.01, ***: p<0.001).

Because TNFα signaling is primarily mediated through the canonical TNFα Receptor 1 (TNFαR1) and TNFα Receptor 2 (TNFαR2) transmembrane receptor proteins ^44^, we next characterized the abundance and organization of these receptors. IF quantification showed significantly higher baseline expression of TNFαR2, but not TNFαR1, in mature cells **(Fig. 2d and Supplemental Fig. S2d)**. To examine receptor organization at the nanoscale, we performed super-resolution imaging and cluster analysis of TNFα receptor distributions. While TNFαR1 clustering was unaffected by treatment **(Supplementary Fig. S2e)**, TNFα stimulation induced a marked increase in TNFαR2 cluster density on the surface of mature cells **(Fig. 2e)**. Given that receptor clustering is known to potentiate downstream signal propagation ^44^, these findings suggest enhanced TNFα signaling magnitude in mature tenocytes.

Consistent with this, RT-PCR analysis demonstrated significant upregulation of inflammation-related genes, particularly MMP-13 and Caspase 11, in TNFα-treated mature tenocytes at the 6-hour timepoint, whereas young cells showed relatively minimal transcriptional induction **(Fig. 2f)**. Together, these results indicate that maturation enhances TNFα receptor organization and downstream inflammatory gene activation, potentially amplifying the inflammatory response despite comparable initial NFκB activation rates between age groups. These findings further reveal a transcriptional divergence between young and mature tenocytes following TNFα stimulation, underscoring an age-dependent differential response to inflammatory signaling in tendon.

### 2.3 Age-Specific Chromatin Accessibility and Pro-inflammatory Transcription Factor Activation in Inflammation-Stimulated Tenocytes

Given the age-dependent differences in chromatin remodeling under inflammatory conditions, we next investigated which genomic regions experienced differential chromatin accessibility during TNFα stimulation. To this end, we performed Assay for Transposase-Accessible Chromatin with sequencing (ATAC-seq) on tenocytes derived from young and mature donors. The 6-hour TNFα treatment time point was selected because it produced the strongest chromatin reorganization and transcriptional response observed in earlier analyses. Correlation among biological replicates per condition and peak distributions within consensus peaks confirmed dataset quality **(Supplemental Fig. S3a, b)**. Principal component analysis (PCA) of the top 500 differential peaks showed that TNFα treatment accounted for the majority of the dataset’s variance **(Supplemental Fig. S3c)**; however, excluding peaks greater than 3 kb away from a transcriptional start site (TSS) improved separation of age groups along principal component (PC) 2 **(Supplemental Fig. S3d)**.

Pairwise comparison of differentially accessible regions (DARs) between untreated young and mature cells (Young Control and Mature Control, respectively) revealed 8,085 regions with increased and 9,521 with decreased accessibility in young cells **(Supplemental Fig. S4a, b)**. The top-ranked increased and decreased DARs are listed in **Tables S1, S2**, and the corresponding gene annotation results are indicated in **Supplemental Fig. S4c, d**. Using peaks within 3 kb of the TSS (TSS-associated peaks), Kyoto Encyclopedia of Genes and Genomes (KEGG) pathway analysis featured differential enrichment in cytoskeletal organization and mechanobiological signaling pathways **(Supplemental Fig. S4e, f)**. The top-ranked transcription factor motifs identified using TSS-associated DARs included *Fra-1* and *Aft4* in young cells, and *HLF* and *NFĸB* in mature cells **(Supplemental Fig. S4g and Tables S3, S4)** ^45^.

In comparison, DARs generated by comparing young TNFα-treated and mature TNFα-treated groups (Young TNFα and Mature TNFα, respectively) revealed 5,271 regions with increased and 9,410 regions with decreased accessibility in young cells **(Supplemental Fig. S5a, b)**, with the top-ranked increased and decreased DARs listed in **Tables S5, S6**. Gene annotation revealed that peaks with increased accessibility were predominantly TSS-associated, whereas decreased peaks were more distal **(Supplemental Fig. S5c, d)**. KEGG enrichment analysis of TSS-associated DARs indicated altered accessibility of mTOR, Wnt, and PI3K signaling pathways **(Supplemental Fig. S5e, d)**.

To control for baseline accessibility differences, we next compared TNFα-treated samples to their respective controls within each age group (e.g., Young TNFα vs Young Control). Most TNFα-induced DARs in both age groups were upregulated, with promoter regions comprising ∼25% of all peaks **(Supplemental Fig. S6Aa-d and Supplemental Fig. S7a-d)**. The most significantly increased and decreased peaks are summarized in **Tables S7, S8** for young tenocytes, and **Tables S9, S10** for mature tenocytes. Interestingly, consensus peak regions showed distinct age-related responses following TNFα treatment: young tenocytes exhibited an overall reduction in peak accessibility, whereas mature cells showed increased accessibility **(Supplemental Fig. S6e and Supplemental Fig. S7e)**. KEGG analysis using TSS-associated DARs revealed alterations in multiple conserved signaling pathways across both age groups, including inflammation- and mechanobiology-related processes **(Fig. 3a, b; Supplemental Fig. S6f; Supplemental Fig. S7f)**.

**Figure 3:**
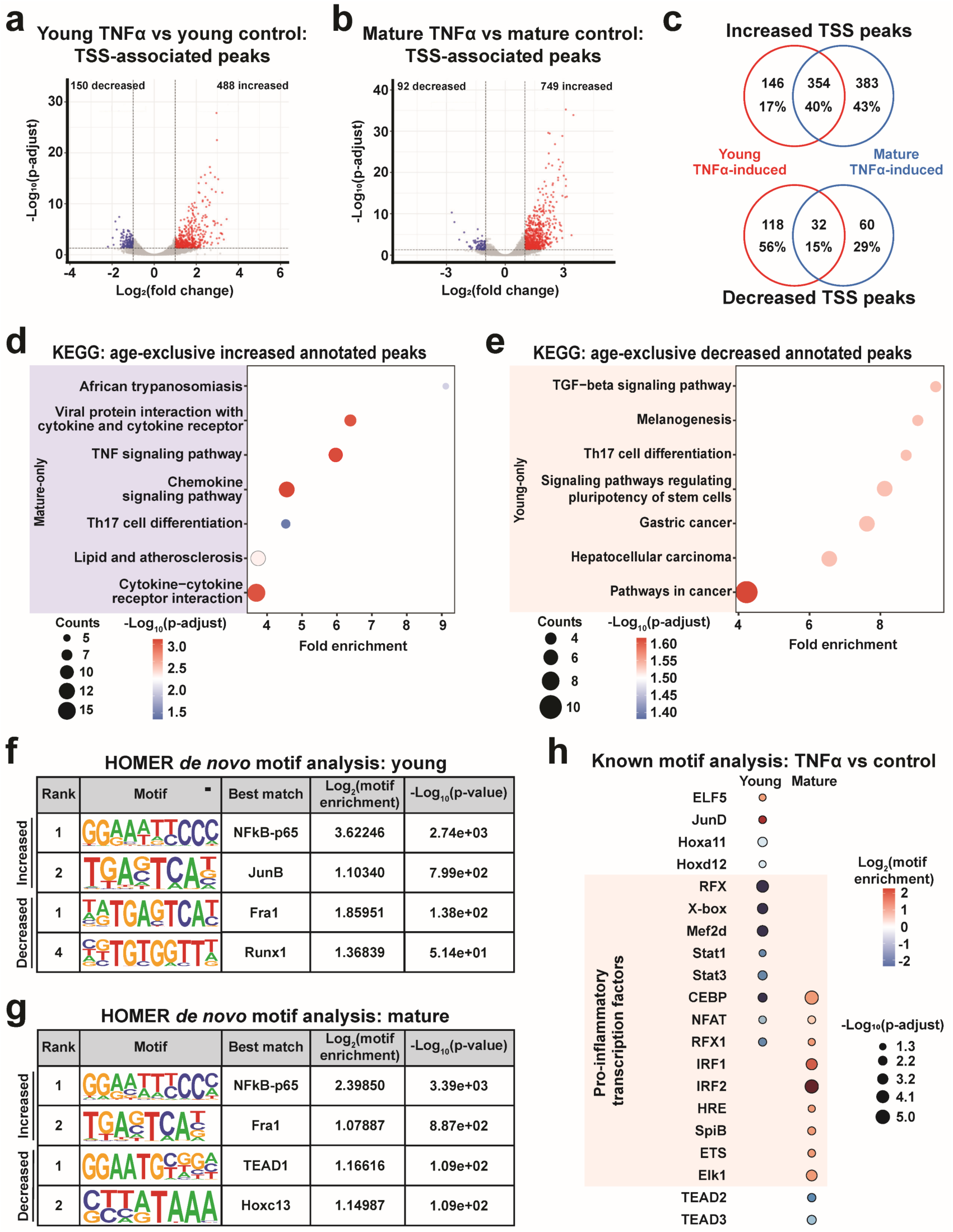
Mature tenocytes display unique chromatin accessibility and transcription factor motif activity under inflammatory stimulation. (a, **b)** Volcano plots of differentially accessible regions (DARs) within ±3 kb of transcription start sites (TSS-associated) comparing TNFα-treated and control conditions in young **(a)** and mature **(b)** tenocytes. Peaks with significantly increased (red) or decreased (blue) accessibility are indicated. **(c)** Schematic illustrating identification of age-exclusive TSS-associated peaks for young (red) and mature (blue) groups following TNFα treatment. **(d, e)** Kyoto Encyclopedia of Genes and Genomes (KEGG) pathway enrichment analysis of age-exclusive TSS-associated peaks showing pathways with increased **(d)** and decreased **(e)** accessibility. Color denotes significance, and circle size indicates gene count. Young and mature data are represented in red and blue, respectively. (f, g) *De novo* transcription factor binding motif enrichment of TSS-associated peaks from young **(f)** and mature **(g)** tenocytes associated with increased or decreased accessibility. **(h)** Age-exclusive known motif enrichment of TSS-associated peaks from young (left) and mature (right) tenocytes using previously annotated binding motif sequences. Color represents motif enrichment, circle size indicates significance, and pro-inflammatory factors are highlighted in pink. (n=2 biological replicates, TNFα treated for 6 hours, significance (DARs): |Log_2_(fold change)|>1 and p-adj<0.05, significance (*de novo* motifs): |Log_2_(motif enrichment)|>1 and p-value<1e-13).

To delineate age-specific responses, TSS-associated DARs were partitioned into age-exclusive (significant in one age group only) and shared (significant in both age groups) categories **(Fig. 3c)**. KEGG analysis of age-exclusive peaks with increased accessibility revealed significant pathway enrichment exclusively from mature-specific peaks **(Fig. 3d)**, including TNFα signaling, cytokine and chemokine signaling, and pro-inflammatory T helper 17 cell (Th17) differentiation **(Fig. 3d)**. In contrast, KEGG analysis of age-exclusive peaks with decreased accessibility showed enrichment only from young-exclusive peaks **(Fig. 3e)**, notably in TGF-β signaling and Th17 differentiation **(Fig. 3e)**. Collectively, these results show that mature tenocytes preferentially activate unique pro-inflammatory chromatin states, while young cells maintain a balanced immunomodulatory conformation. Although inflammation broadly alters chromatin accessibility, age-specific mechanisms distinctly regulate the magnitude and direction of the immune response ^46^.

We next investigated the nuclear factors responsible for driving age-dependent inflammatory pathway activation. To this end, we turned to *de novo* motif enrichment analysis using the Hypergeometric Optimization of Motif Enrichment (HOMER) platform ^45^. Using TSS-associated DARs identified from TNFα vs control comparisons in young and mature tenocytes, the top non-redundant transcription factor motifs are listed in **Tables S11, S12** for young tenocytes and **Tables S13, S14** for mature tenocytes. As expected, *NFĸB* was the most enriched motif for increased peaks in both groups, consistent with canonical inflammatory signaling. However, members of the AP-1 transcription factor family (e.g., *Fra1*, *JunB*, *c-Jun*) exhibited distinct age-dependent regulation under TNFα stimulation. Young cells appeared to increase the accessibility at *Jun* subfamily motifs but reduced accessibility at *Fra1* sites, whereas *Fra1* ranked as the second most enriched motif in mature cells **(Fig. 3f, g)**. AP-1 family transcription factors, including *Jun* and *Fra1*, can form diverse sets of heterodimers that influence sequence specificity and loci targeting within the nucleus ^47–49^. These results suggest that aging modulates AP-1 family activity through altered motif accessibility of specific heterodimer configurations.

Other *de novo* enriched motifs identified across both age groups included Hox family genes (i.e., *Hoxc13*), TEAD components (e.g., *Tead2*, *Tead1*), and CEBP family members (e.g., *Cebpa*, *Cebpg*) **(Fig. 3f, g)** ^50–52^. In addition to *de novo* motif enrichment, validated motifs identified from previously published chromatin immunoprecipitation data were used to determine known motif sequences present within TNFα-induced DARs as well. Significant differentially accessible known motifs within both age groups were compared to generate a list of age-exclusive known motifs for young and mature tenocytes with TNFα. Age-exclusive motifs associated with pro-inflammatory signaling and responses (e.g., RFX family, IRF family, *Stat1* ^53–55)^ were all associated with decreased accessibility within young cells and increased accessibility within mature cells **(Fig. 3h)**. Other transcription factors, including specific members of the Tead and Hox families, also showed age-exclusive accessibility changes.

Together, these results reveal distinct transcription factor accessibility landscapes between young and mature-aged tenocytes under inflammatory stimulation. The enhanced accessibility of AP-1 family factors (i.e., *Fra1*) and pro-inflammatory RFX and IRF motifs in mature cells suggest that age-dependent chromatin condensation at these loci drives differential transcriptional responses, contributing to the heightened inflammatory phenotype observed in aging tendon cells.

### 2.4 Age-dependent Transcriptional Reprogramming of Tenocytes in Response to Inflammatory Stimulation

Given the age-dependent differences in chromatin remodeling and epigenetic alterations under inflammatory conditions, we further investigated how inflammation differentially alters transcriptional programs in young and mature tenocytes. To achieve this, RNA-sequencing (RNA-seq) was performed using three biological replicates of young and mature tenocytes, with donor-matched cells treated with TNFα for 6 hours.

PCA showed a clear separation of TNFα-treated conditions across PC1, with additional separation by age across PC2 **(Fig. 4a)**. Differential analysis of control groups (Young Control and Mature Control) identified 273 upregulated and 467 downregulated differentially expressed genes (DEGs) **(Supplemental Fig. S8a, b and Supplemental Tables S15, S16)**. Gene ontology (GO) analysis indicated that these genes were enriched for vascular development factors, cytoskeletal components, and ECM regulation **(Supplemental Fig. S8c)**. In contrast, comparison of TNFα-treated samples (Young TNFα and Mature TNFα) revealed 208 upregulated and 316 downregulated DEGs **(Supplemental Fig. S9a, b and Supplemental Tables S17, S18)**. Young TNFα-treated cells showed elevated expression of ECM components (*Tnn* and *Col2a1*) and TGF-β signaling (*Gdf10* and *Lefty1*), whereas Mature TNFα-treated cells upregulated pro-inflammatory cues (*IL33* and *Cxcl12*) and chromatin structural proteins (*Hist1h2a*). GO enrichment further confirmed these divergent responses. Young cells activated pathways governing ECM maintenance and homeostasis (ECM organization, Extracellular structure) and resolution of inflammation (Regulation of inflammatory response, Regulation of angiogenesis, Regulation of IL-6 production), while mature cells exhibited increased expression of genes linked to immune activation (Complement activation, Leukocyte migration, Macrophage proliferation) **(Supplemental Fig. S9c)**. These findings indicate that inflammation drives an age-dependent shift from reparative processes in young cells to inflammatory propagation in mature cells ^9^.

**Figure 4:**
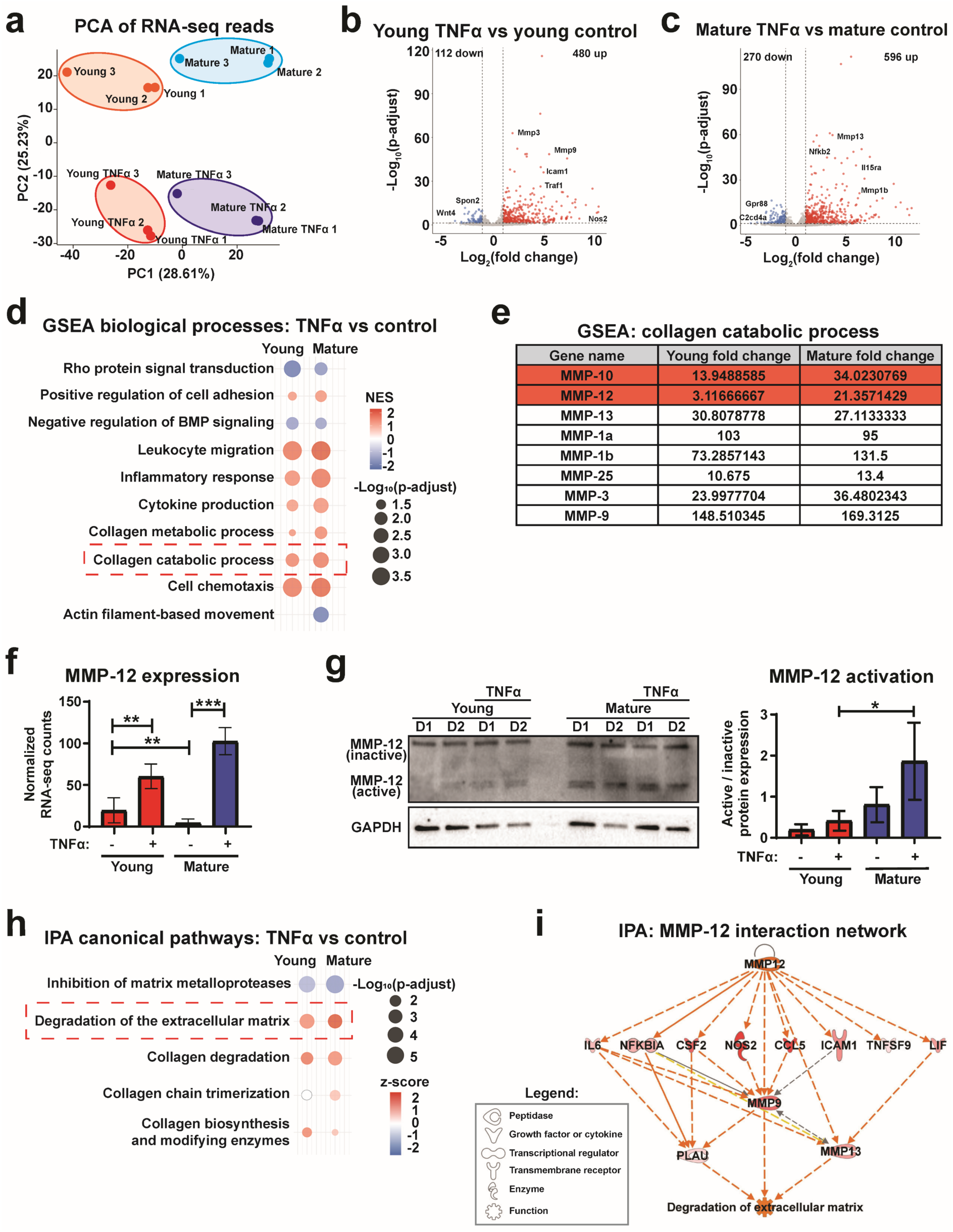
RNA-seq reveals an induction of matrix-degrading pathways in mature tenocytes following inflammatory stimulation. **(a)** Principal component analysis (PCA) of RNA-seq gene expression from young and mature tenocytes under control (blue and red, respectively) and TNFα-treated (6-hour; purple and green, respectively) conditions. **(b, c)** Volcano plots of differentially expressed genes (DEGs) comparing TNFα-treated to control tenocytes in young **(b)** and mature **(d)** groups. Significantly upregulated (red) and downregulated (blue) genes are indicated. **(d)** Gene set enrichment analysis (GSEA) of biological processes derived from DEGs in young (left) and mature (right) tenocytes comparing TNFα-treated versus control conditions. Color denotes normalized enrichment score, and circle size indicates significance. **(e)** List of DEGs within the “Collagen catabolic process” GSEA category, showing fold-change values between TNFα-treated and control cells. Genes highlighted in red exhibit >1.4-fold differences between young and mature comparisons. **(f)** Normalized RNA-seq counts for MMP-12 in young (red) and mature (blue) tenocytes after TNFα stimulation. (*: p-adj<0.05, ***: p-adj<0.001). **(g)** Western blot quantification (left) and representative blot image (right) showing the activation ratio of MMP-12 protein (active enzyme / inactive pro-enzyme) in young (red) and mature (blue) tenocytes following 6-hour TNFα treatment. (*: p<0.05, ***: p<0.001). **(h)** Ingenuity Pathway Analysis (IPA) of canonical pathways comparing TNFα-treated and control tenocytes in young (left) and mature (right) groups. Color represents z-score, and circle size indicates significance. **(i)** IPA interaction network illustrating MMP-12-mediated matrix degradation. Orange lines denote predicted activation, gray lines indicate indirect interactions, and yellow lines mark inconsistent expression effects. Darker symbol color represents higher confidence in predicted activation. (n=3 biological replicates per group, TNFα treated for 6 hours, significance: |Log_2_(fold change)|>1 and p-adj<0.05, Mean ± SD).

To dissect the contributions of TNFα separately from baseline age differences, DEGs were next identified by comparing TNFα-treated samples to their respective controls (e.g., Young TNFα and Young Control). The majority of TNFα-induced DEGs were upregulated in both age groups, with a greater number of significantly altered genes observed in mature tenocytes **(Fig. 4b, c; Supplemental Fig. S10a, b; Supplemental Tables S19-S22)**. Shared pathways enriched using GO and KEGG analyses among upregulated TNFα-induced DEGs included innate immune activation and chemokine signaling, whereas downregulated pathways were associated with cell identity and mesenchymal differentiation **(Supplemental Fig. S10c-e)**.

Notably, young tenocytes showed higher expression of ECM synthesis and collagen maturation processes, whereas mature cells preferentially upregulated catabolic and inflammatory pathways. Gene set enrichment analysis (GSEA) of TNFα-induced DEGs revealed enrichment of collagen remodeling (Collagen catabolic process) and immune cell infiltration (Leukocyte migration) pathways in mature cells **(Fig. 4d)**. Nearly all significantly expressed matrix metalloprotease (MMP) enzymes were elevated more in mature cells, with MMP-12 and MMP-10 in particular showing greater than 1.4-fold induction compared to young cells **(Fig. 4e)**. Normalized MMP-12 transcript counts confirmed its elevated transcriptional expression **(Fig. 4f)**, and western blot analysis confirmed increased activation of MMP-12 proteins, as shown by the higher ratio of active to inactive enzyme forms **(Fig. 4g)** ^56^. Ingenuity Pathway Analysis (IPA) corroborated these findings, displaying enhanced ECM degradation and reduced MMP inhibition in mature cells **(Fig. 4h)**. The “Degradation of the ECM” IPA pathway network further highlighted the central role of MMP-12 in inducing downstream proteolytic cascades **(Fig. 4i)**.

Analysis of predicted upstream regulators from IPA enriched for chromatin remodeling pathways revealed significant predicted activation only in mature tenocytes **(Supplemental Fig. S11a, b)**. Likewise, genes associated with the “Chromatin Organization” GO term overlapped with age-exclusive DEGs exclusively from mature cells **(Supplemental Fig. S11c)**. These data suggest that mature tenocytes displayed a unique gene expression response to TNFα treatment governing chromatin remodeling factors, potentially facilitating the global rearrangements described previously.

To further identify pathways uniquely activated in each age group, TNFα-induced DEGs were separated into age-exclusive and shared subsets **(Fig. 5a)**. KEGG and GO pathway enrichment analyses showed that mature-specific pathways included JAK-STAT, HIF-1, and FOXO signaling, while young-specific pathways included Hippo, NOTCH, and cellular adhesion signaling **(Supplemental Fig. S12a)**. GO terms further emphasized ECM maturation and organization in young cells compared to pro-inflammatory signaling in mature cells **(Fig. 5b, c)**. Within ECM-related processes, young-exclusive genes were predominantly upregulated, whereas mature-exclusive genes were largely downregulated **(Supplemental Fig. S12b-e)**, supporting a transition from homeostatic to catabolic transcriptional programs with aging ^16^.

**Figure 5:**
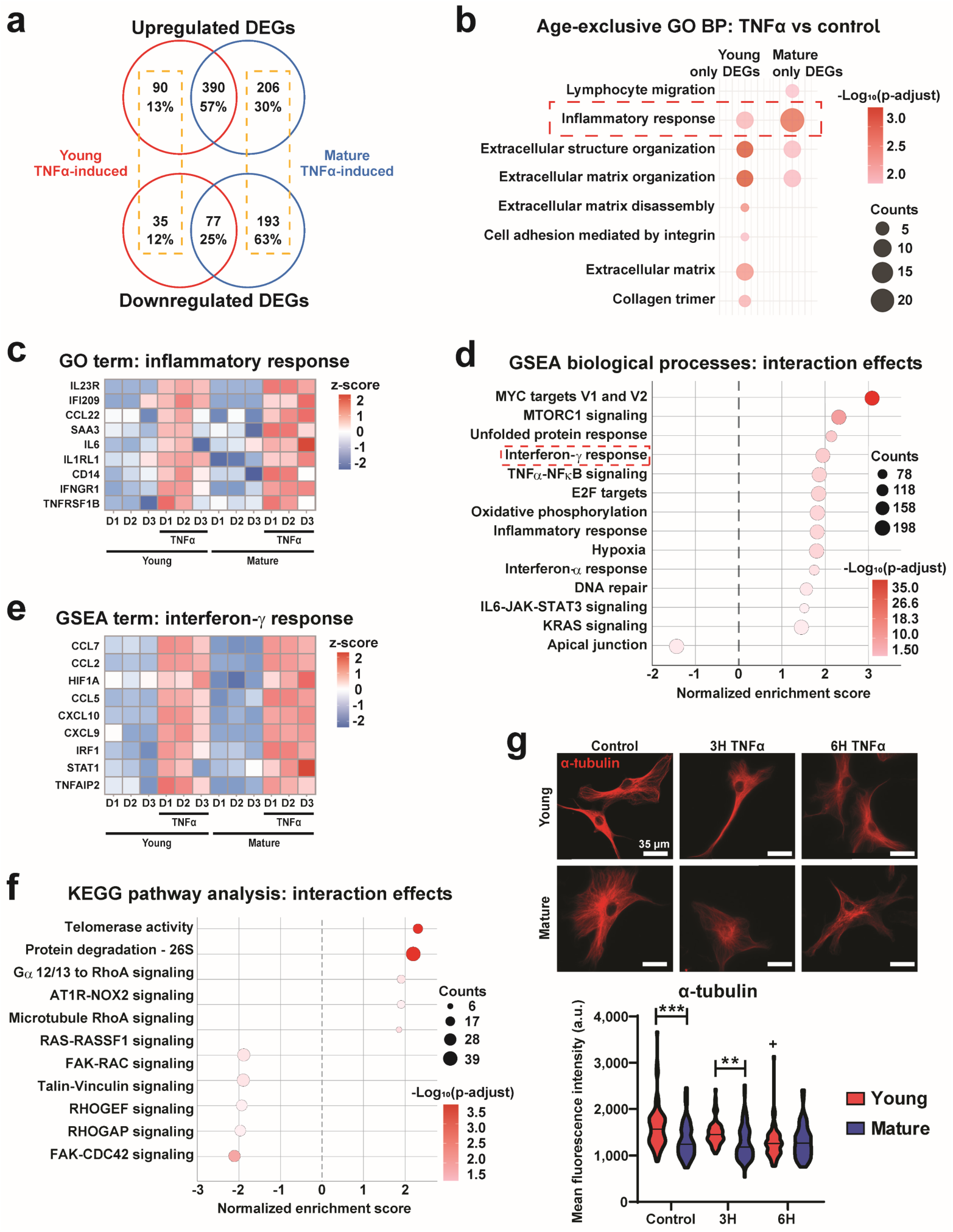
Age-exclusive inflammatory pathway activation in young and mature tenocytes. **(a)** Schematic illustrating the identification of age-exclusive differentially expressed gene (DEG) sets by comparing TNFα-treated and control groups in young (red) and mature (blue) tenocytes. Yellow boxes denote age-exclusive DEG groupings used for downstream pathway analyses. **(b)** Gene Ontology (GO) biological process (BP) enrichment analysis of age-exclusive DEGs comparing TNFα-treated and control tenocytes in young (left) and mature (right) groups. Color indicates statistical significance, and circle size represents gene counts. **(c)** Heatmap of age-exclusive DEGs within the GO term “Inflammatory response” for young and mature tenocytes under control and TNFα-treated conditions. Color represents z-score–scaled expression. **(d)** Pre-ranked Gene Set Enrichment Analysis (GSEA) of biological processes derived from two-way interaction analyses (Age × Treatment) of TNFα-induced DEGs. A positive normalized enrichment score (NES) indicates greater enrichment in mature tenocytes, color denotes significance, and circle size indicates gene counts. **(e)** Heatmap of TNFα-induced DEGs within the “Interferon-γ response” GSEA term for young and mature tenocytes. Color represents z-score–scaled expression. **(f)** Pre-ranked Kyoto Encyclopedia of Genes and Genomes (KEGG) pathway analysis based on two-way interaction results (Age × Treatment) of TNFα-induced DEGs. A positive NES reflects greater enrichment in mature tenocytes, color indicates significance, and circle size denotes gene counts. **(g)** Representative immunofluorescence images and mean fluorescence intensity quantification of α-tubulin in young (red) and mature (blue) tenocytes under control and TNFα-treated conditions. (n=60 cells/group, mean ± SD, +: p<0.05 vs Young Control, **: p<0.01, ***: p<0.001). (n=3 biological replicates, TNFα treated for 6 hours, significance: |Log_2_(fold change)|>1 and p-adj<0.05).

A complementary two-way interaction analysis identified genes that responded to TNFα in both age groups but with different magnitudes of change. The 19 genes showing the most significant interaction effects are listed in **Table S23**. The resulting interaction statistic was used for rank ordering of GSEA and KEGG enrichment. The pre-ranked GSEA identified multiple inflammation-related pathways with positive enrichment scores (Interferon-γ Response, NFĸB Signaling), indicating a larger increase in expression was observed within mature TNFα-induced DEGs **(Fig. 5d, e)**. Other processes with stronger induction in mature tenocytes include mTORC1 signaling, DNA repair, and Hypoxia. KEGG enrichment of 2-way interaction-associated pathways indicated a strong decrease in multiple mechanoregulatory factors in young cells, including microtubule dynamics, focal adhesions, and Rho signaling **(Fig. 5f)**. Consistent with this, immunofluorescence analysis confirmed decreased microtubule intensity after TNFα treatment only in young cells **(Fig. 5g)**.

Taken together, RNA-seq analyses revealed that mature tenocytes exhibit heightened expression and activation of MMPs and pro-inflammatory signaling pathways, while young tenocytes favor novel ECM synthesis and maintenance. This age-dependent transcriptional reprogramming leads to a shift toward matrix catabolism and chronic inflammation in mature cells, potentiating a loss of tendon homeostasis, structural integrity, and repair capacity ^10,57^.

### 2.5 Consequences of Inflammatory Induction with Tenocyte Age

Thus, mature tendon cells show distinct molecular and epigenetic characteristics compared to their young counterparts even at baseline, and these differences likely underlie their heightened sensitivity to inflammatory stimuli. The epigenetic landscape, defined by chromatin accessibility and histone modification patterns, regulates the ability of transcription factors and other regulatory elements to bind DNA ^23^. This, in turn, modulates the magnitude and duration of cellular responses to inflammatory cues, even when upstream signaling and nuclear translocation remain comparable ^37,58^. Our findings demonstrate that the epigenetically primed state of mature tenocytes promotes further chromatin compaction and repression of tenocyte-specific gene programs upon inflammatory challenge. This leads to enhanced matrix degradation and reduced ECM synthesis, ultimately impairing the regenerative capacity of mature and aged tendon tissue ^59^. Such maladaptive responses reinforce a degenerative feedback loop, leading to further inflammatory propagation and degeneration, which progressively compromises the tendon’s structure and healing potential **(Fig. 6)** ^60,61^.

**Figure 6:**
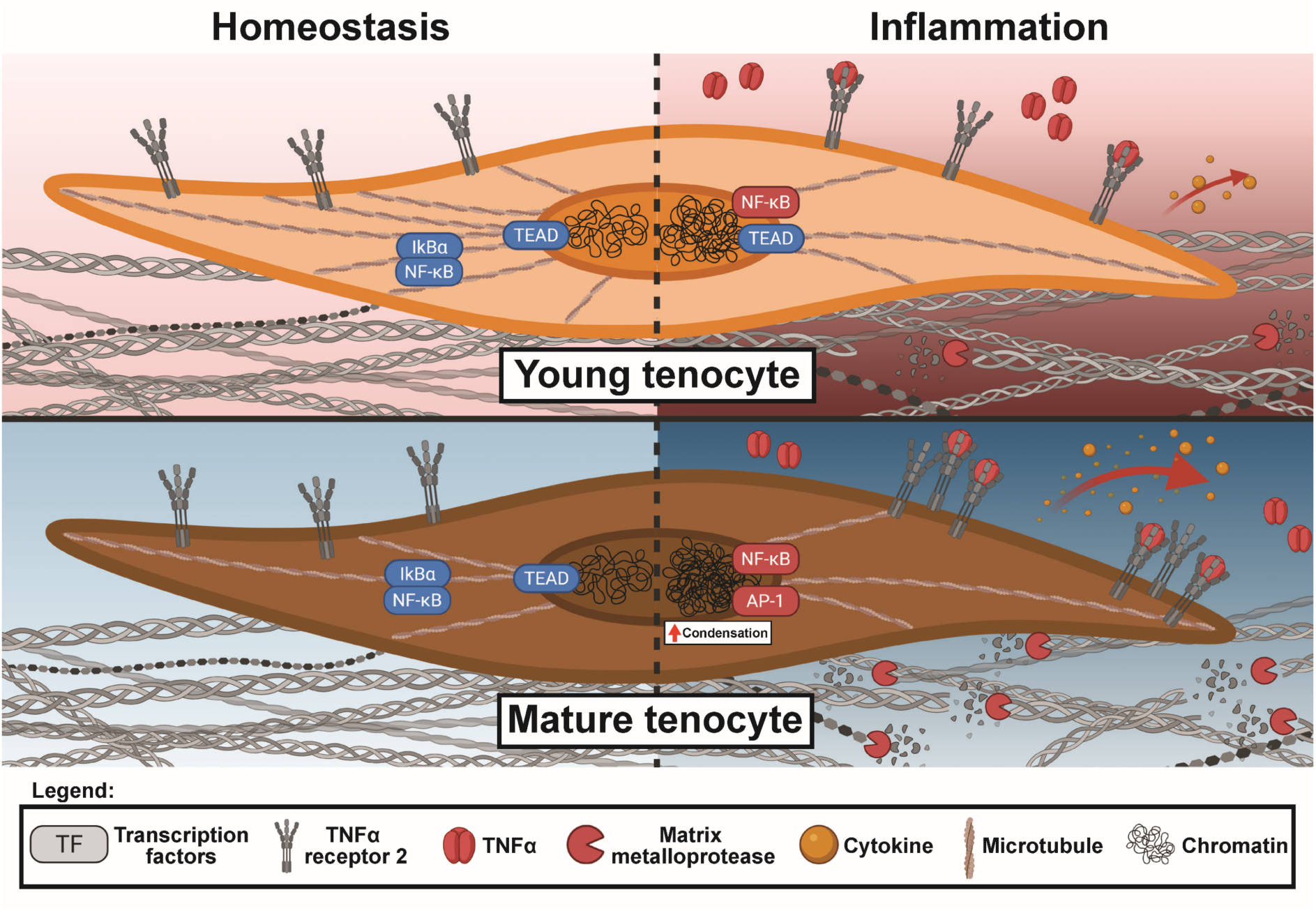
Summary of the cumulative effects of tenocyte aging on inflammatory responsiveness. Schematic illustration of the proposed mechanism underlying enhanced inflammatory sensitivity in mature tenocytes. Panels compare homeostatic (left) and inflammatory (right) conditions in young (top) and mature (bottom) tenocytes. Red elements represent pro-inflammatory signals or factors, whereas blue elements denote anti-inflammatory or homeostatic components. The diagram summarizes how maturation and aging shift the tenocyte response to inflammatory stimulation from a balanced, reparative state to one characterized by heightened chromatin compaction, amplified cytokine signaling, and increased matrix catabolism.

## 3. Discussion

Aging profoundly alters the tendon microenvironment and cellular phenotype, rendering tendon tissue more susceptible to chronic inflammation and impaired repair ^1,15–17^. In this study, we demonstrate that age-dependent chromatin remodeling underlies distinct transcriptional and functional responses to inflammatory stimuli in tenocytes. Inflammation and aging are intricately interconnected, forming a self-reinforcing feedback loop that drives cellular and tissue dysfunction across multiple organ systems ^62–64^. Age-associated changes in intracellular signaling and systemic immune regulation promote chronic, low-grade tissue inflammation ^61,65^, which disrupts normal healing cascades following acute injury ^12,66^ and contributes to the diminished regenerative potential of aged tissues ^32,64,67,68^. In the context of tendon, chronic inflammation is associated with impaired mechanical strength and resilience, compromising tissue integrity and force transmission efficiency ^69–71^. Although these outcomes are partly driven by global alterations in immune system activity and specificity ^72,73^, our data identify stromal cell-intrinsic mechanisms within aged tendon tissue that contribute to this phenotype. Specifically, we show that mature tenocytes themselves become active amplifiers of inflammation through enhanced transcriptional activation of cytokines, chemokines, and catabolic enzymes ^74^.

Mechanistically, our findings indicate that maturation alters both the signaling architecture and structural organization of inflammatory pathway components. Elevated expression of multiple inflammatory mediators in mature cells reflects an increased amplitude of inflammatory signaling with age ^75^. In addition, mature tenocytes exhibit enhanced clustering of TNFα Receptor 2 upon ligand engagement, which likely concentrates downstream signaling complexes and amplifies the magnitude of inflammatory response despite comparable NFκB nuclear translocation rates between age groups ^44^. This receptor-level sensitization, combined with elevated transcription of inflammatory mediators, suggests that mature tendon stromal cells play an active role in perpetuating inflammation rather than passively responding to it. Such amplification may contribute to the delayed resolution of inflammation observed in mature or aged tendon healing ^9,12^.

The primary physiological role of tenocytes is to maintain ECM homeostasis and respond to extrinsic mechanical demands through balanced anabolic and catabolic activity ^25,76^. During normal tendon function, microdamage to collagen fibrils occurs routinely and is often efficiently repaired without compromising mechanical integrity ^77^. However, sustained low-grade inflammatory signaling can disrupt these conventional repair cues, leading to a catabolic imbalance ^8^. Over time, the accumulation of unresolved microdamage in the tissue eventually drives a transition point beyond which tendon integrity and function decline ^13,77^. Our RNA-seq data support this model, showing that mature tenocytes under inflammatory stress exhibit elevated expression of matrix-degrading enzymes, including catalytically active MMPs, alongside reduced ECM synthesis compared to young cells. This imbalance shifts ECM homeostasis towards degeneration, driving progressive loss of tissue integrity. Furthermore, the upregulation of pro-fibrotic factors such as fibronectin in mature tenocytes may exacerbate fibrosis, further limiting the regenerative potential of the tissue ^78,79^. Chronic inflammation also promotes immune cell recruitment, enhancing matrix degradation and tissue remodeling through immune cell–derived proteases ^80,81^. Thus, both systemic inflammaging and cell-intrinsic reprogramming in aged tenocytes contribute to a feed-forward loop of matrix breakdown and persistent inflammation.

At the epigenetic level, our study reveals that genome organization is a central determinant of these age-divergent responses. Consistent with previous findings in other cell types ^82–84^, mature tenocytes display increased heterochromatin density and reduced chromatin plasticity at baseline compared to young cells. This more compact nuclear state may constrain the dynamic remodeling capacity of the genome, limiting transcriptional flexibility and solidifying high accessibility at pro-inflammatory loci ^85–87^. In parallel, our ATAC-seq analysis identified increased chromatin accessibility in mature tenocytes at motifs associated with pro-inflammatory transcription factors such as AP-1 (*Fra-1*), RFX, and IRF families ^48,53,54^, whereas young cells retained accessibility at loci involved in ECM maintenance and repair (e.g., *Tead*). In particular, elevated *Fra-1* binding activity may drive aberrant inflammatory responses, as AP-1 family factors regulate proliferation, stress adaptation, and apoptosis ^88,89^. Consistently, we observed elevated caspase activation in mature cells, suggesting that heightened AP-1 activity may couple inflammatory signaling to cell death and tissue degeneration.

Notably, we also observed differential regulation of chromatin regions associated with T helper 17 (Th17) differentiation pathways between young and mature tenocytes. Given that Th17 responses are central to many autoimmune and chronic inflammatory diseases ^46,90,91^, these findings suggest that mature tenocytes may adopt epigenetic states that favor autoimmune-like chronic inflammation. This aligns with our observation of increased expression of inflammatory cytokines and chemokines, suggesting that mature tenocytes may contribute directly to immune recruitment and persistence within tendon lesions.

While this study establishes a mechanistic link between maturation, chromatin remodeling, and inflammation, several limitations should be acknowledged. Our experiments were conducted in vitro using two-dimensional culture systems, which do not fully recapitulate the native mechanical and structural environment of tendon tissue. Tendons are highly anisotropic and significantly softer than standard culture substrates ^92^. These physical properties likely influence tenocyte mechanotransduction and inflammatory behavior. Nevertheless, prior evidence suggests that the observed pro-inflammatory and catabolic phenotypes are largely driven by signaling and epigenetic mechanisms rather than substrate stiffness alone ^22,25^. Moreover, the use of TNFα as a single inflammatory stimulant provides a controlled but simplified model of inflammation; in vivo, a complex mixture of cytokines, growth factors, and immune cell paracrine signals interact simultaneously ^36^. Despite this, our RNA-seq data revealed broad activation of multiple inflammatory pathways, supporting the physiological relevance of TNFα-induced responses. Future work incorporating advanced-aged (geriatric) animal models and *ex vivo* tendon systems will be crucial to validate these findings under native biomechanical and cellular contexts.

In summary, our data demonstrate that aging predisposes tenocytes to a primed chromatin state, which amplifies inflammatory signaling and matrix catabolism upon stimulation. This epigenetically reinforced sensitivity, driven by altered chromatin architecture and transcription factor accessibility, shifts tendon homeostasis from a reparative state to a degenerative trajectory. Understanding how chromatin-level regulation interfaces with mechanical and inflammatory signaling provides a critical foundation for developing therapies targeting epigenetic and transcriptional modulators to restore tendon health in aging populations.

## 4. Conclusions

This study establishes a mechanistic connection between age-related epigenetic remodeling in tenocytes and the diminished healing potential observed in aged tendon tissue ^59^. We demonstrate that tenocytes derived from young and mature animals display fundamentally distinct transcriptional and chromatin accessibility responses to identical inflammatory stimuli, driven in part by differential epigenetic priming. Mature tenocytes preferentially adopt a pro-inflammatory and catabolic transcriptional state, characterized by enhanced matrix degradation and cellular stress responses, which may provide a molecular framework for the chronic degeneration and impaired regenerative outcomes observed clinically in aging tendons. These findings further identify matrix-degrading enzymes, including MMP-12, as potential therapeutic targets for mitigating tendon degeneration in aging or inflammation-prone conditions. More broadly, our results suggest that selective modulation of age-dependent chromatin and transcriptional programs may offer a strategy to rebalance inflammatory signaling and restore tissue homeostasis. Future studies aimed at defining, targeting and reprogramming these age-exclusive regulatory pathways will be essential for developing rejuvenation-focused strategies that enhance repair capacity in inflamed or aged tendon tissues.

## 5. Materials and methods

### 5.1 Cell Isolation

Pure CD1, pure C57BL/6, or mixed CD1/C57BL/6 male mice were used as tenocyte donors. Young mice (4 weeks old) and mature mice (45–50 weeks old) were designated as the “young” and “mature” groups, respectively. Briefly, tail tendon fascicles were surgically harvested from donor mice following established protocols ^93^. The isolated fascicles were digested in an enzymatic media containing 0.4% (w/v) collagenase IV (Worthington LS004188), 0.3% (w/v) dispase II (Thermo 17105041) in DMEM, high glucose culture media (Thermo 11965084). Digestion was carried out at 37°C with shaking for 2 hours. The digestion solution was diluted 30-fold with basal culture media (DMEM, high-glucose media supplemented with 1% (v/v) PS solution and 10% (v/v) fetal bovine serum (Corning MT35010CV)) and centrifuged, with the resulting cell-tissue pellet passed through a 100 µm mesh (Thermo 352360) and plated in tissue culture-treated plates (6-well: Falcon 087721B or 10cm: CellTreat CT229621) in fresh basal media.

### 5.2 Cell Culture and Inflammatory Treatment

Cell cultures were maintained below 80% confluency, with media changes performed every 2 days as needed. All experiments were conducted using cells at passage 0 (P0) or 1 (P1). Prior to treatment, tenocytes were cultured for two days in basal medium to allow for stabilization. Inflammatory signaling was induced by supplementing the culture medium with 20 ng/mL recombinant mouse TNFα (Sigma T7539-50-UG) during a media change ^26^. The inflammatory stimulus remained in the culture media for either 0.5, 1, 3, 6, or 24 hours before fixation or extraction and analysis, depending on the experimental group.

### 5.3 Tenocyte Viability and Proliferation Assays

Tenocyte viability under inflammatory conditions was assessed using the LIVE/DEAD Viability Kit (Thermo L3224). Tenocytes were seeded onto slide glass (EMS 71887-87) and cultured for up to 48 hours under inflammatory or control conditions using basal media containing phenol red-free DMEM (Thermo 21063029). At 0, 6, 24, and 48 hours, assay reagents were applied to cells according to the manufacturer’s protocol, and samples were immediately imaged using the Leica DM6000 Widefield 20× air objective. Cell viability was quantified as the ratio of live (green-positive, red-negative) cells to total cell count for each condition.

Cell proliferation was determined using the Cell Counting Kit-8 (Sigma C859K87). Tenocytes were cultured in phenol red-free basal media within 6-well plates for 0, 6, 24, or 48 hours in inflammatory or control conditions before adding the CCK8 reagent. Plates were incubated for 2 hours, protected from light, before reading the absorbance at 450 nm. Proliferation values were normalized to the 0-hour control for each age group.

### 5.4 Stochastic Optical Reconstruction Microscopy (STORM) Imaging

Tenocytes were fixed within chambered slide glass culture devices (Thermo 155411) using a cold 50:50 MeOH-EtOH solution for 5 minutes (mins) at room temperature (RT) following TNFα treatment or control culture conditions. Samples were blocked with BlockAid solution (Thermo B10710) for 1 hour at RT before incubating overnight at 4°C with the primary antibody (1:50 dilution for anti-Histone H2B (Thermo PA5-115361), 1:100 dilution for anti-TNFαR1 (Enzo ADI-CSA-815-D) or anti-TNFαR2 (Abcam AB15563). A custom donkey anti-rabbit secondary antibody (Jackson 711-005-152) conjugated with activator-reporter dye pairs of AF-405 and AF-647 was applied for 1 hour at RT ^22^. Immediately before imaging, a MEM-based imaging buffer was prepared and added to the chambers to facilitate photoswitching events ^94^. The imaging buffer consisted of 36mM HCl (Lab Alley HCASA1N-1L), 99.8mM cysteamine (Sigma 30070-10G), 5% (w/v) glucose (Thermo A16828-36), 0.56mg/mL glucose oxidase (Sigma G2133-10KU), 0.4mM NaCl (Sigma 20-159), 0.05mg/mL catalase (Sigma C40) and 80µM Tris buffer (Sigma 648314-100ML) in DPBS (Thermo 14190136) ^94^. Imaging was performed on the Nanoimager S (ONi) (100× oil immersion objective, NA 1.49, sCMOS Hamamatsu Ocra-Flash camera) with TIRF inclination. Cell membrane- or nuclear-stained regions were imaged for 25,000 frames at a 15ms exposure time while gradually increasing the laser power to maintain a consistent photoblinking density.

### 5.5 STORM Analysis

Localization of single-molecule blinking events and drift correction were performed in NimOS (ONi) to reconstruct an overall image, and rendering was done using a custom ImageJ plugin, Insight3 ^95^. Localization data were further analyzed using previously published MATLAB scripts implementing Voronoi tessellation-based clustering to quantify nanoscale localization density ^22^. Dense chromatin regions were defined as domains within the 41st–80th percentile of Voronoi area distributions, and chromatin condensation was assessed by the inverse of the 80th percentile reduced Voronoi area value per nucleus (n=10 nuclei per condition) ^22^. Percentage changes in chromatin density were calculated by comparing 80th percentile dense chromatin values between treatment and control groups. For visualization, density heatmaps were generated using a custom MATLAB script that color-coded Voronoi polygons based on inverse reduced areas ^22^. For TNFα receptor analyses, localization clusters were identified from membrane-associated regions of interest (ROIs), and receptor density was calculated as the average localization count per cluster across multiple cells (n = 10 cells per condition, 3 cytoplasmic ROI per cell).

### 5.6 Immunofluorescence (IF) Imaging

Following TNFα treatment or control culture, tenocytes were fixed using 4% paraformaldehyde solution (Thermo J19943) for 30 mins at RT on slide glass. Cells were then permeabilized using a solution of 0.5% (v/v) Triton X-100 (Sigma X100-100ML), 314.6 mM sucrose (Sigma S0389-500G), and 6.4 mM MgCl_2_ (Sigma 208337-1KG) in DPBS for 5 mins at RT, followed by incubation in a blocking solution of 3% bovine serum albumin (BSA) (Sigma A1933-5G) in DPBS for 1 hour at RT.

Primary antibodies were diluted in a solution of 3% BSA in DPBS and incubated overnight at 4°C. Primary antibodies used included anti-H3K27me3 (Thermo PIPA531817, 1:400), anti-H3K4me3 (Thermo MA5-11199, 1:400), anti-H3K9ac (Thermo MA5-11195, 1:400), anti-TNFαR1 (Enzo ADI-CSA-815-D, 1:333), anti-TNFαR2 (Abcam 1031239-3, 1:333), anti-YAP (Santa Cruz Bio sc-101199, 1:300), anti-αtubulin (Abcam ab18251, 1:1000), and anti-NFĸB (Thermo 51-0500, 1:200). Goat anti-rabbit secondary antibodies with AF-549 (Thermo A-11035) or AF-488 (Thermo A-11008) were added to the samples for 1 hour at RT. Secondary antibody staining of YAP samples was performed using anti-mouse AF-549 (A-11005). After secondary antibody staining, NFĸB- and YAP-stained samples were additionally treated with Phalloidin-488 (Thermo A12379) (1:300 dilution in 3% BSA solution) for 1 hour at 37°C. Prolong gold DAPI dye (Thermo P36935) was applied to samples immediately before imaging. Tenocytes were imaged using the Leica DM6000 Widefield 20× air objective, capturing multiple fields of view per sample.

### 5.7 Immunofluorescence (IF) Analysis

IF image analysis was performed using ImageJ. The nuclear ROI was selected based on the DAPI (nuclear) channel, while the cytoplasmic ROI was defined using the phalloidin (cytoskeleton) channel to delineate the cell area. Data reported as mean intensity per ROI. For quantification of the nuclear-to-cytoplasmic (N:C) intensity ratio, the cytoplasmic ROI was defined to exclude the nuclear region. Mean fluorescence intensity within each ROI was measured, and the N:C ratio was calculated by dividing nuclear by cytoplasmic intensity values for each cell. Cell area was determined from the phalloidin-stained region without subtracting the nuclear ROI.

### 5.8 Reverse Transcription Polymerase Chain Reaction (RT-PCR) and Analysis

Total RNA was extracted and purified from tenocyte cultures using the Direct-zol RNA Microprep kit with Tri Reagent (Zymo Research R2063) following TNFα treatment (3 or 6 hours) or control culture conditions. RNA concentration was quantified using a Nanodrop One spectrophotometer (Thermo ND-ONE-W) and subsequently used for cDNA synthesis with the ProtoScript II kit (New England Biolabs E6560L). RT-PCR was performed using Fast SYBR Green quantification (Thermo 4385616) on the QuantStudio 3 (Thermo A28136). Ct values for target genes were normalized to the housekeeping gene (i.e., GAPDH ^96^) for each condition (ΔCt), and these values were then normalized to control group gene expression (ΔΔCt) within each age group before fold change was calculated (primers listed in **Table S24**). Statistics were performed on ΔΔCt values.

### 5.9 Assay for Transposase-Accessible Chromatin with Sequencing (ATAC-seq)

Tenocytes (n = 2 per group from 2 different donors) were collected using 0.25% Trypsin-EDTA (Thermo 25200056) after 6 hours of TNFα treatment or control culture conditions. A total of 100,000 cells were used for tagmentation and library preparation using an ATAC-seq kit (Active Motif 53150). Purified libraries were quantified using a Nanodrop One spectrophotometer, and library size distribution was measured using the high-sensitivity DNA bioanalyzer kit (Agilent 5067-4626) on the Agilent 2100 Bioanalyzer system. Libraries were sequenced by Azenta (GENEWIZ) using 2×150 bp, 375 Gb configuration, with 60 million reads per sample and 5% PhiX normalization spike-in. Upstream analysis, including read trimming, alignment, QC, and peak annotation, was performed in R (Bowtie2, MACS3, ChIPseeker_1.42.1). Downstream analysis of reads, including differential analysis, utilized DESeq2, EnhancedVolcano_1.18.0, clusterProfiler_4.8.3, pheatmap_1.0.12.

### 5.10 ATAC-seq Differential Analysis

Differentially accessible regions (DARs) were determined within each age group, comparing TNFα to control conditions, and within each treatment condition, comparing young to mature groups. Significance was defined as a p-adjust value < 0.05 and a fold change threshold of |Log_2_(Fold Change)| > 1 to identify DARs. Pairwise DARs were analyzed separately based on the directionality of the fold change (i.e., increased or decreased peaks) or analyzed after combining all peaks per comparison. Following gene annotation, significant peaks within 3 kb of transcription start sites (TSSs) were retained and analyzed separately as TSS-associated peaks. Kyoto Encyclopedia of Genes and Genomes (KEGG) analysis was performed on TSS-associated peaks with FDR < 0.05 and a minimum pathway size set to 4. Age-exclusive DAR lists were determined by comparing TSS-associated DAR lists derived from the TNFα vs Control comparisons of young and mature tenocytes and selecting annotated genes present in only one age group. Age-exclusive DARs were used for subsequent KEGG pathway enrichment analysis of upregulated and downregulated regions, separately.

### 5.11 Motif Enrichment Analysis

Motif enrichment analysis of DARs was performed using Hypergeometric Optimization of Motif Enrichment (HOMER_5.1), utilizing TSS-associated DARs identified from TNFα vs control (young and mature) and young vs mature (control) comparisons as input ^45^. Top-ranked non-redundant *de novo* motifs with a similarity score > 0.9 and a p-value < 1e^-13^ were considered for analysis. Known motif analysis was performed using TNFα vs control (young and mature) conditions, and significant results were defined as possessing a |Log_2_(Motif Enrichment)| > 1 and p-adjust < 0.05, where Motif Enrichment is calculated as the percentage of target sequences with a motif divided by the percentage of background sequences with a motif. Significantly regulated known motif results were then compared between the young and mature groups to identify age-exclusive known motifs present in one age group.

### 5.12 RNA Sequencing (RNA-seq)

RNA samples were extracted from 6-hour TNFα-treated and control cell cultures using TRIzol reagent (Thermo 15596026) (n = 3 per group from 3 different donors). RNA concentrations were determined using a Nanodrop spectrophotometer. Library preparation, quality control, and subsequent sequencing were performed by Azenta (GENEWIZ). Sequencing was conducted at a depth of 20 million paired-end reads (2x150bp) with ERCC spike-in controls. Initial trimming and alignment were performed by Azenta, and downstream analysis was performed in R (DESeq2_1.40.2, EnhancedVolcano_1.18.0, clusterProfiler_4.8.3, fgsea_1.32.2, and pheatmap_1.0.12) and Ingenuity Pathway Analysis (IPA) (Qiagen).

### 5.13 RNA-seq Differential Analysis

Significance was defined as a p-adjust value < 0.05 and a fold change threshold of |Log_2_(Fold Change)| > 1 to determine differentially expressed genes (DEGs). DEGs were identified within each age group, comparing TNFα conditions to control conditions, and within each treatment condition, comparing young to mature individuals. Gene ontology (GO) analysis of DEGs was performed using Biological Processes, Cellular Component, or Molecular Function term categories with FDR < 0.05 and a minimum pathway size set to 4. KEGG, IPA, and gene set enrichment analysis (GSEA) analyses were performed with FDR < 0.05 and a minimum pathway size set to 4. Age-exclusive DEG lists were determined by comparing DEG lists derived from the TNFα vs control comparisons of young and mature tenocytes and selecting DEGs present within only one age group. Age-exclusive DEGs were used for subsequent pathway enrichment analysis using KEGG and GO databases. IPA upstream regulators identified using TNFα vs control DEG lists for each age group were compared to GO terms of interest to determine specific pathways overrepresented within the predicted upstream regulators lists, including the “Chromatin organization” GO term.

Two-way interaction analysis was performed in R (DESeq2_1.40.2) using all reads and the variables of age and treatment. The resulting test statistic was used to rank all genes for downstream GSEA and KEGG analyses, indicating pathways associated with distinct magnitudes of TNFα response between age groups. The 19 genes with |Log_2_(Fold Change)| > 1 and p-adjust < 0.4 from interaction analysis were clustered into 4 categories of shared response behavior to both variables.

### 5.14 Protein Extraction and Western Blot

Whole protein lysate was extracted from tenocytes under control and inflammatory (6-hour) conditions using RIPA buffer (Thermo 89900) (n = 3 per group from three different donors). Total protein concentration was determined using the Qubit 4 BR Protein Assay Kit (Thermo A50668). Protein samples were denatured at 95°C for 10 mins (Biorad 1610747). 7.4µg of protein was loaded per lane of 4-20% MP TGX gels (Biorad 4561095) and run for 1.2 hours at constant 110V in Tris-Glycine-SDS running buffer (Biorad 1610772). Protein transfer occurred with constant 2.5A, 25V conditions for 7 mins semi-dry using 0.2µm nitrocellulose membranes (Biorad 1704270) and transfer buffer (Biorad 10026938) with 20% ethanol (Thermo A4094).

Membranes were blocked using 3% BSA in Tris-buffered saline (Biorad 1706435) with 1% Tween 20 (Thermo AAJ20605AP) (TBST) with rocking for 1 hour at RT. Primary antibodies for MMP-12 (Thermo PA5-13181) and GAPDH (Proteintech 10494-1-AP) were diluted in TBST 1:2000 and 1:3000, respectively, and incubated overnight with rocking at 4°C. Secondary anti-rabbit conjugated to HRP enzyme (Thermo PRW4011) was diluted 1:3000 in TBST and incubated for 1 hour at RT. All washing steps were performed for 10 mins 3x with TBST. GAPDH membranes were incubated with Clarity western ECL substrate (Biorad 170-5061) covered from light for 5 mins before imaging for chemiluminescence using the ChemiDoc XRS+ system (Biorad), and MMP-12 membranes were incubated with Clarity Max western ECL substrate (Biorad 1705062).

### 5.15 Western Blot Analysis

Quantification of band intensity was performed in ImageLab (Biorad) and ImageJ. The background signal of gels directly above and below the band of interest was subtracted from the band area intensity for all targets. The resulting band intensities for GAPDH were used to normalize protein loading content per lane between samples, and MMP-12 band values were scaled accordingly. 2 MMP-12 bands were analyzed per lane, corresponding to the inactive pro-enzyme form at 54kDa and the activated enzyme form at 44kDa ^56^. The ratio of 44 kDa to 54 kDa band intensities was reported as an indicator of the activation status of MMP-12 per sample.

### 5.16 Statistical Analysis

Outliers were identified and excluded using the 1.5 × interquartile range method. Data normality was assessed using the Shapiro-Wilk test. For comparisons between two conditions, a two-tailed Student’s t-test was applied to normally distributed data, while the Mann-Whitney test was used for non-normal distributions. For comparisons among more than two conditions, a two-tailed one-way analysis of variance (ANOVA) with Tukey post-hoc test was used for normally distributed data, or a non-parametric one-way ANOVA for non-normal data. To evaluate the effects of treatment and age simultaneously, a two-tailed two-way ANOVA with Tukey’s post-hoc testing was performed. Statistical significance was defined as a p-value < 0.05 with α = 0.8. All data reported as mean + standard deviation with sample size indicated.

## 6. Data availability

All sequencing data is publicly available in an NCIB Sequence Read Archive repository (PRJNA1370051). Any additional datasets generated during the current study are available from the corresponding author on reasonable request.

## Acknowledgments

We would like to acknowledge the use of the Penn Center for Musculoskeletal Disorders Biomechanics Core (NIH P30 AR069619), the University of Pennsylvania Perelman School of Medicine Cell and Developmental Biology Microscopy Core (SCR 022373), the Hypergeometric Optimization of Motif Enrichment software (HOMER_5.1) (RRID:SCR_010881), the National Institutes of Health (R01 AR079224, R01 HL163168, P50 AR080581, T32 AR007132), and the National Science Foundation Science and Technology Center for Engineering Mechanobiology (CMMI-1578571). Schematic figures generated using BioRender.com (https://BioRender.com/dm5lwkk).

## Author Contributions

T.E.B., E.Y.Z., I.K.K., N.A.D., I.J., and S.C.H. contributed to the design of the experiments. T.E.B. and E.Y.Z. collected data. T.E.B., E.Y.Z., S.K., B.I.T., X.J., I.K.K., N.A.D., I.J., and S.C.H. contributed to method development. T.E.B., E.Y.Z., S.K., and B.I.T. performed data analysis. T.E.B., E.Y.Z., S.K., B.I.T., I.K.K., N.A.D., I.J., and S.C.H. interpreted the data and contributed to the text. Figures were created by T.E.B., S.K., and B.I.T.

## Data and Materials Correspondence

Direct material and data inquiries to Su Chin Heo.

## Supplemental Materials

**Supplemental Figure S1:**
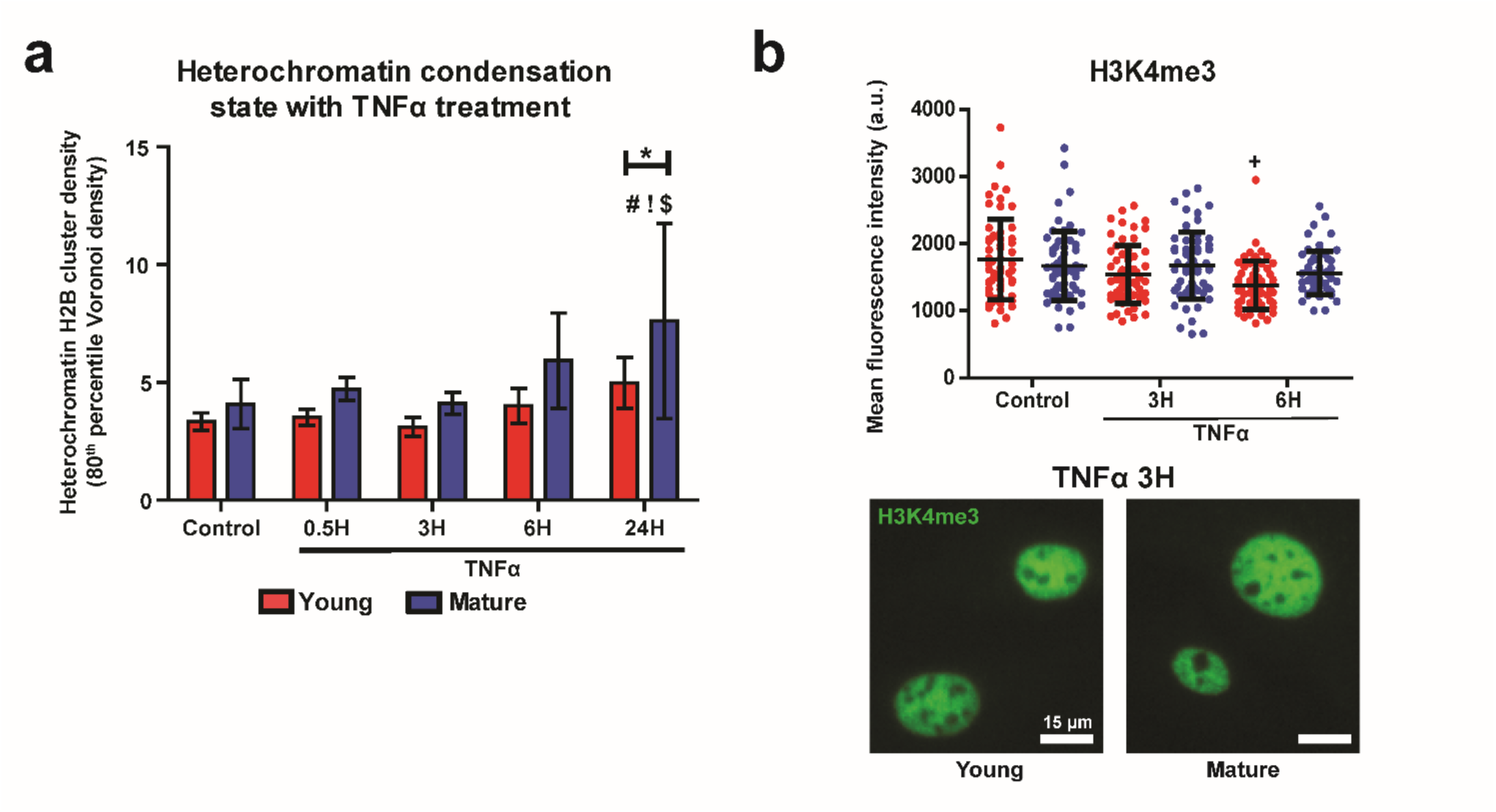
Epigenetic characterization of aged tenocytes under inflammatory treatment. **(a)** Heterochromatin domain density of H2B-STORM localizations reported as the 80th percentile of the Voronoi density cumulative distribution function per cell, normalized to cell area. Shown for control and TNFα-treated conditions in young (red) and mature (blue) tenocytes. (n=16 cells/group). **(b)** Immunofluorescence (IF) mean intensity quantification (top) and representative images (bottom) for histone modification H3K4me3 in young (red) and mature (blue) tenocytes. (n=60 cells/group). (Mean ± SD, +: p<0.05 vs Young Control, #: p<0.05 vs mature Control, !: p<0.05 vs mature 0.5H, $: p<0.05 vs mature 3H, *: p<0.05).

**Supplemental Figure S2:**
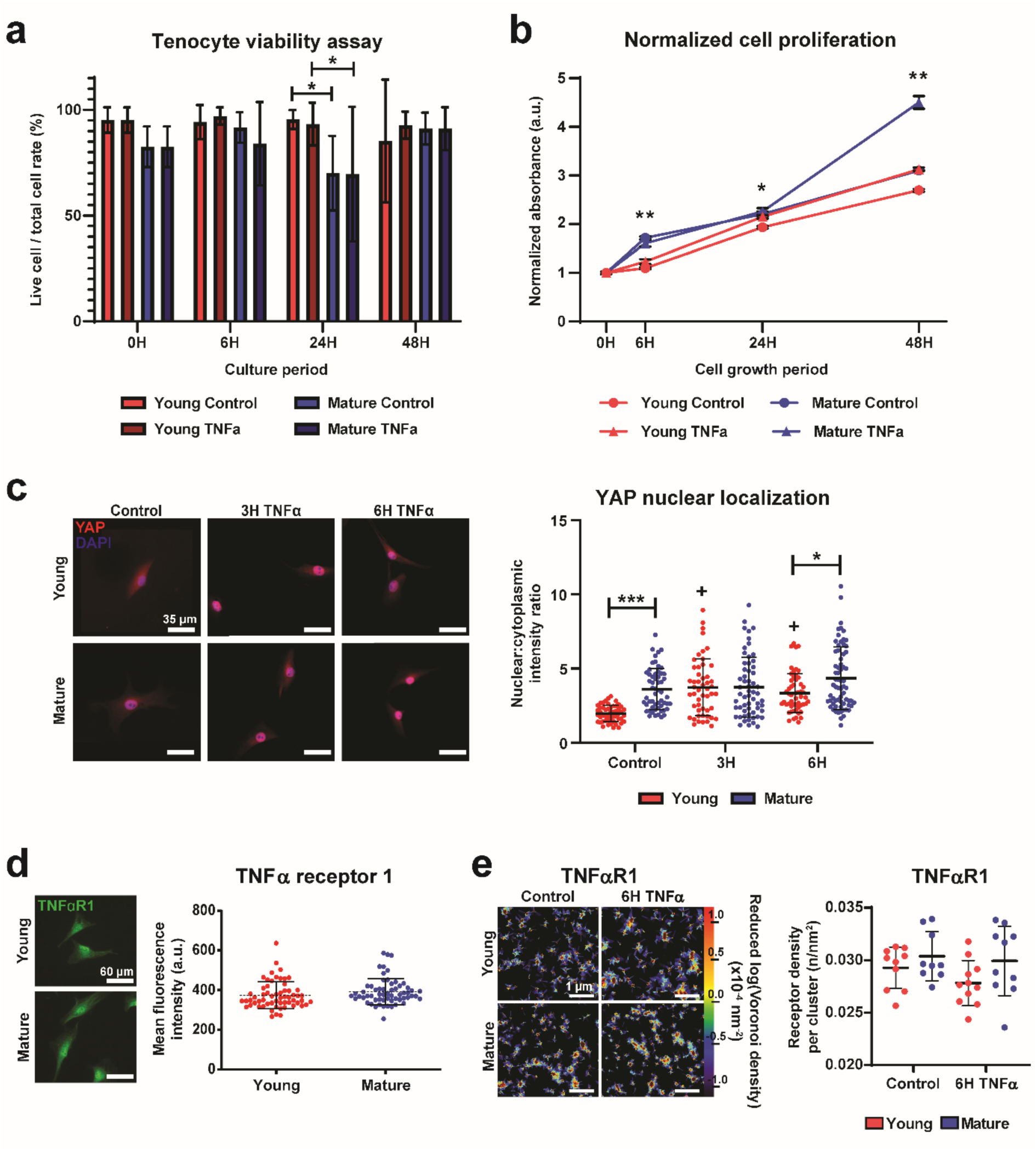
Tenocytes remain viable and mechanobiologically active under inflammatory conditions. **(a)** Viability assay using Live/Dead staining of tenocytes under various control and inflammatory timepoints reported as the percentage of viable cells observed. Young groups colored in red and mature in blue, with TNFα-treated groups in darker colors. (n=100 cells/group). **(b)** Cell proliferation assay of young (red) and mature (blue) tenocytes at control (circle) and inflammatory (triangle) conditions. Values normalized to 0-hour (0H) results for each group. (n=2 technical replicates). **(c)** Representative IF images (left) and nuclear to cytoplasmic ratio (N:C) quantification (right) of YAP in young (red) and mature (blue) control and inflammatory-treated cells. (n=60 cells/group). **(d)** Representative IF images (left) and mean intensity quantification (right) for TNFα Receptor 1 in young and mature tenocytes at control conditions. (n=60 cells/group). **(e)** Representative STORM density heatmaps (left) and quantification of nanoscale receptor cluster density (right) for TNFα Receptor 1 in young (red) and mature (blue) tenocytes at control and inflammatory conditions. (n=11 cells/group, n=3 ROIs/cell). (Mean ± SD, +: p<0.05 vs Young Control, *: p<0.05, **: p<0.01, ***: p<0.001).

**Supplemental Figure S3:**
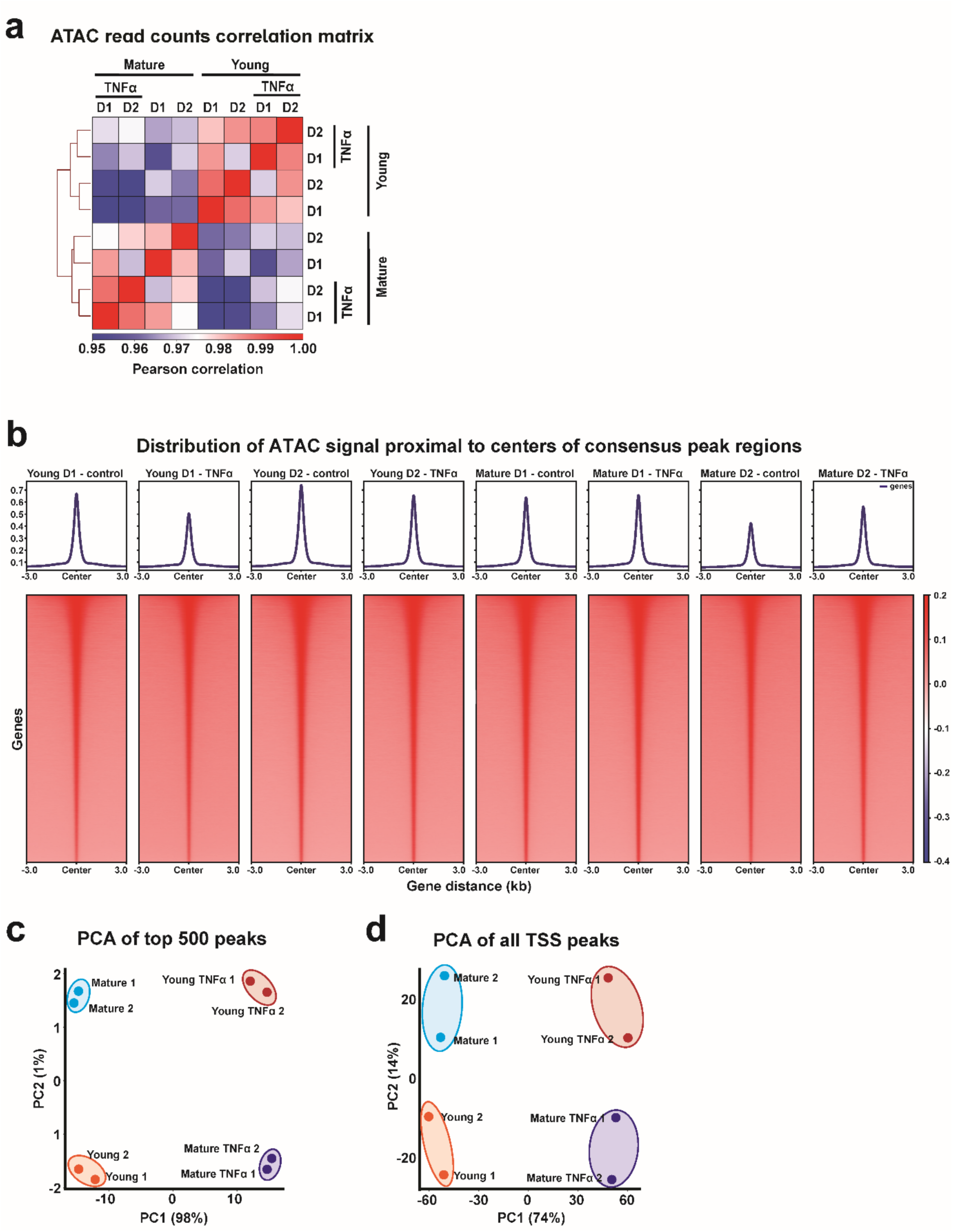
Quality control results of ATAC-seq reads. **(a)** Sample correlation matrix using ATAC-seq read counts in young and mature tenocytes under control and TNFα-treated conditions. Color indicates Pearson correlation coefficient. **(b)** ATAC-seq read distribution profiles across each biological donor for control and TNFα-treated groups within 3 kb of consensus peak regions. **(c)** Principal component analysis (PCA) plot of the top 500 differential peaks comparing control and TNFα-treated young and mature tenocytes. **(d)** PCA plot of all transcription start site-associated (TSS-associated) peaks across control and TNFα-treated conditions in young and mature groups. (n=2 biological replicates per condition, TNFα treated for 6 hours, significance: |Log_2_(fold change)|>1 and p-adj<0.05).

**Supplemental Figure S4:**
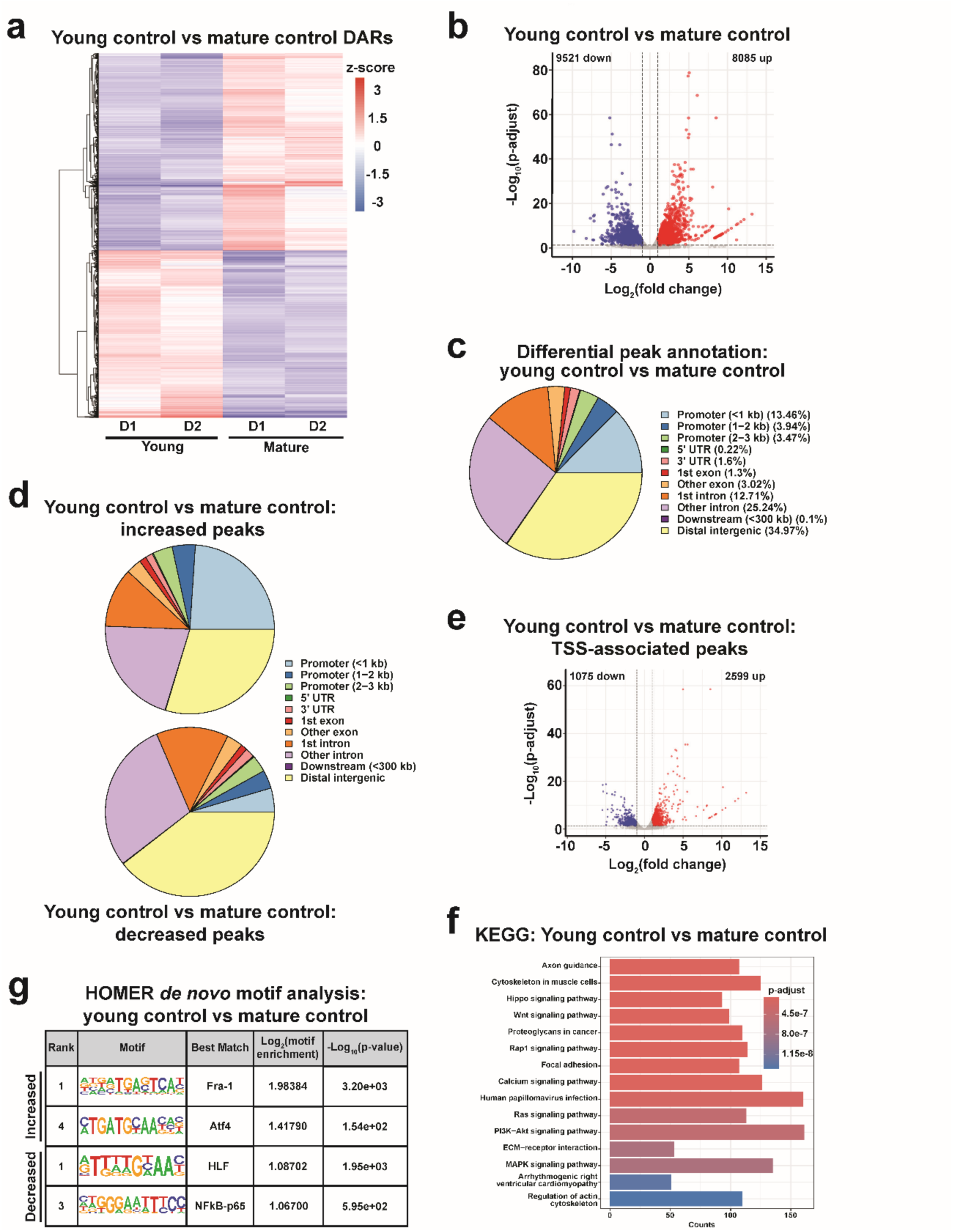
ATAC-seq analysis comparing chromatin accessibility between young and mature control tenocytes. **(a)** Heatmap of differentially accessible regions (DARs) between Young Control and Mature Control tenocytes. Color represents z-score–scaled accessibility. **(b)** Volcano plot of all DARs between Young Control and Mature Control. Peaks with significantly increased (red) or decreased (blue) accessibility are indicated. **(c)** Gene annotation of DARs comparing Young Control and Mature Control. **(d)** Gene annotation results for increased (top) and decreased (bottom) peaks between Young Control and Mature Control group. **(e)** Volcano plot of all TSS-associated peaks between Young Control and Mature Control. Genes with significantly increased (red) or decreased (blue) accessibility are indicated. **(f)** Kyoto Encyclopedia of Genes and Genomes (KEGG) pathway enrichment analysis of TSS-associated DARs comparing Young Control and Mature Control. Color denotes statistical significance. **(g)** *De novo* motif enrichment analysis of increased (top) and decreased (bottom) TSS-associated peaks between Young Control and Mature Control tenocytes.(n=2 biological replicates per condition, significance (DARs): |Log_2_(fold change)|>1 and p-adj<0.05, significance (*de novo* motifs): |Log_2_(motif enrichment)|>1 and p-value<1e^-13^).

**Supplemental Figure S5:**
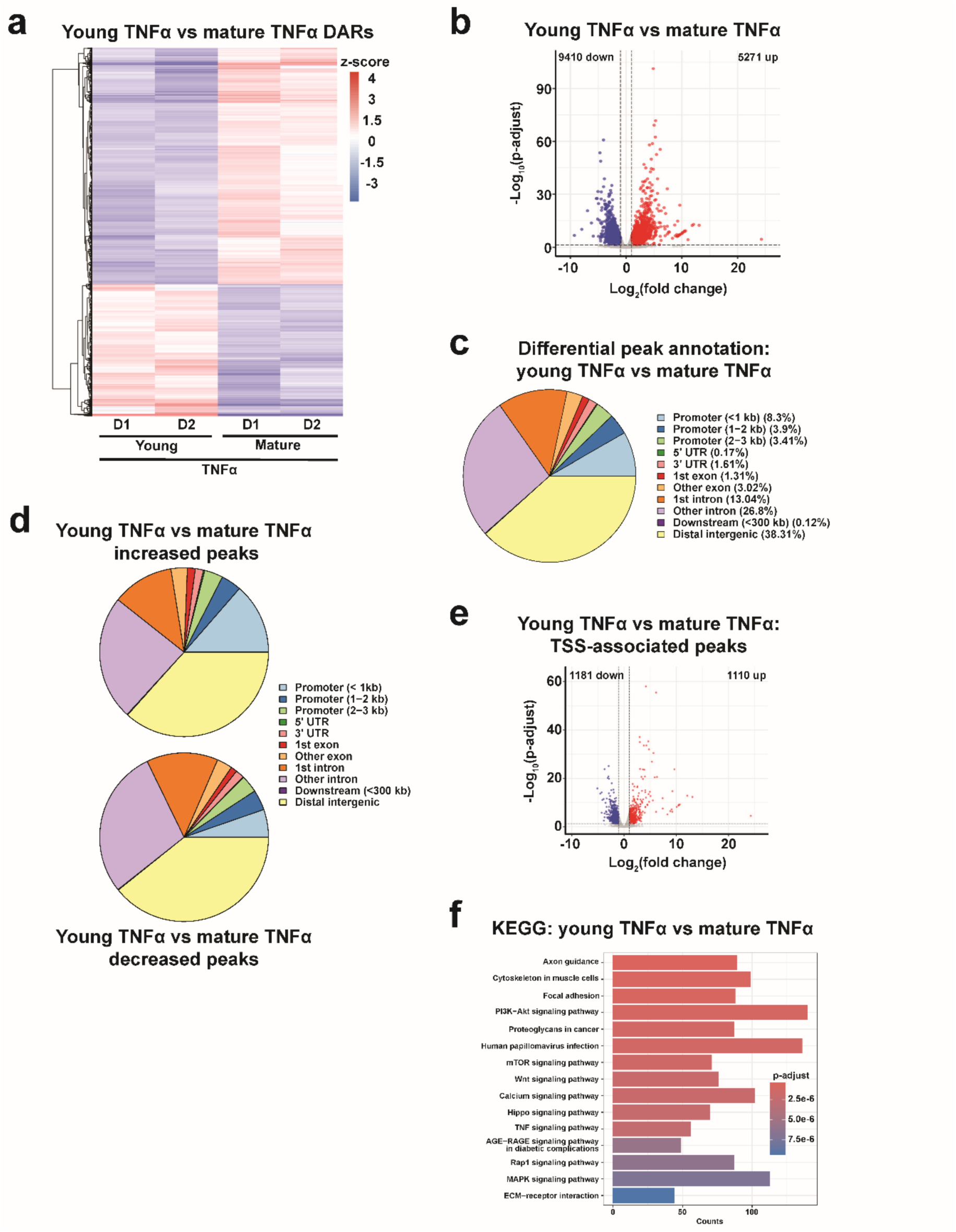
ATAC-seq analysis comparing chromatin accessibility between young and mature tenocytes under TNFα treatment. **(a)** Heatmap of DARs between Young TNFα and Mature TNFα tenocytes. Color represents z-score–scaled accessibility. **(b)** Volcano plot of all DARs between Young TNFα and Mature TNFα. Peaks with significantly increased (red) or decreased (blue) accessibility are indicated. **(c)** Gene annotation of DARs comparing Young TNFα and Mature TNFα groups. **(d)** Gene annotation results for increased (top) and decreased (bottom) peaks between Young TNFα and Mature TNFα. **(e)** Volcano plot of TSS-associated peaks between Young TNFα and Mature TNFα. Genes with significantly increased (red) or decreased (blue) accessibility are indicated. **(f)** KEGG pathway enrichment analysis of TSS-associated DARs comparing Young TNFα and Mature TNFα tenocytes. Color denotes statistical significance. (n=2 biological replicates per condition, TNFα treated for 6 hours, significance: |Log_2_(fold change)|>1 and p-adj<0.05).

**Supplemental Figure S6:**
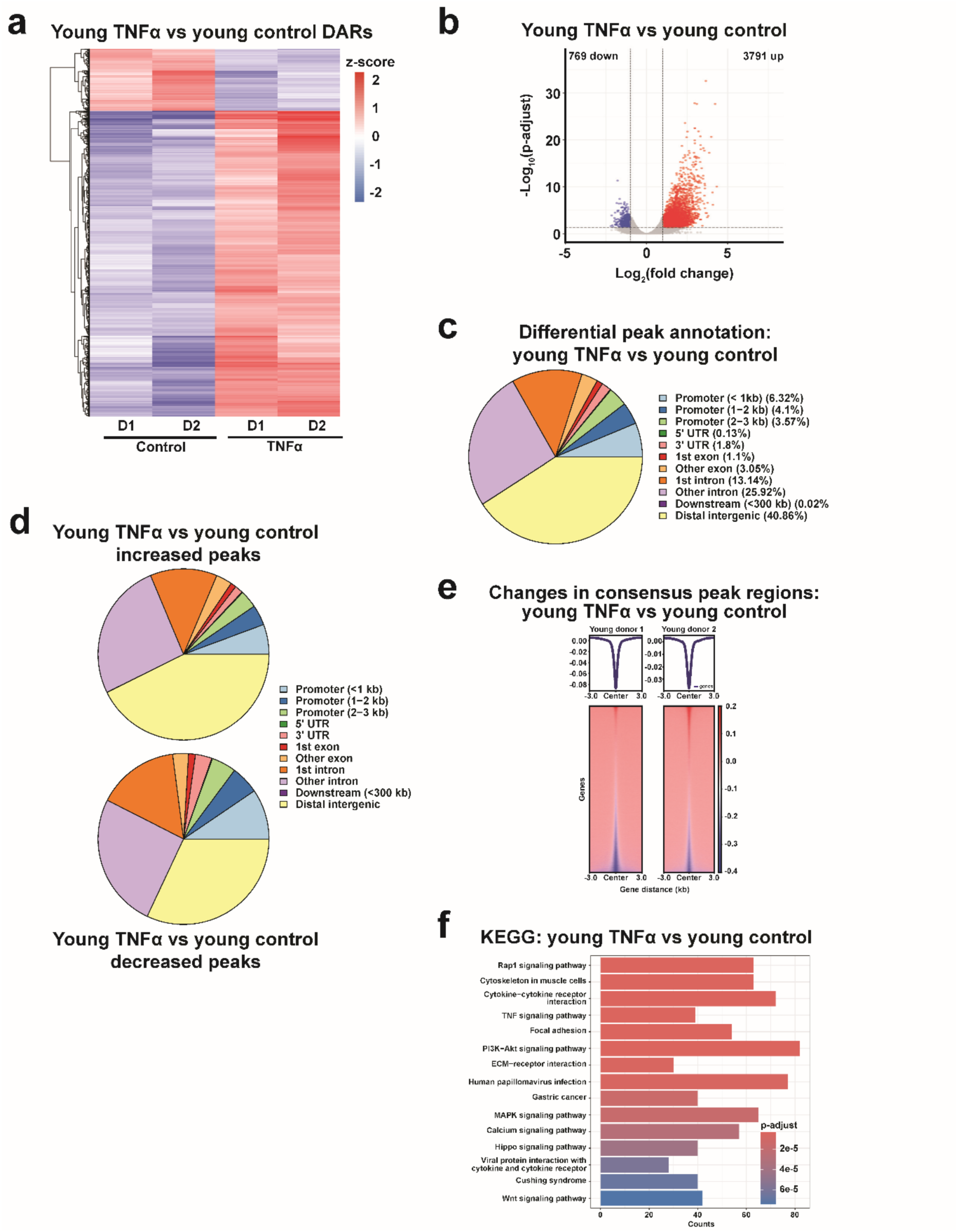
ATAC-seq analysis comparing chromatin accessibility between TNFα-treated and control young tenocytes. **(a)** Heatmap of DARs between Young TNFα and Young Control tenocytes. Color represents z-score–scaled accessibility. **(b)** Volcano plot of all DARs between Young TNFα and Young Control. Peaks with significantly increased (red) or decreased (blue) accessibility are indicated. **(c)** Gene annotation of DARs comparing Young TNFα and Young Control groups. **(d)** Gene annotation results for increased (top) and decreased (bottom) peaks between Young TNFα and Young Control. **(e)** Differential peak enrichment plot of consensus peak regions between Young TNFα and Young Control. Color indicates accessibility changes. **(f)** KEGG pathway enrichment analysis of TSS-associated DARs comparing Young TNFα and Young Control tenocytes. Color denotes statistical significance. (n=2 biological replicates per condition, TNFα treated for 6 hours, significance: |Log_2_(fold change)|>1 and p-adj<0.05).>1 and p-adj<0.05).

**Supplemental Figure S7:**
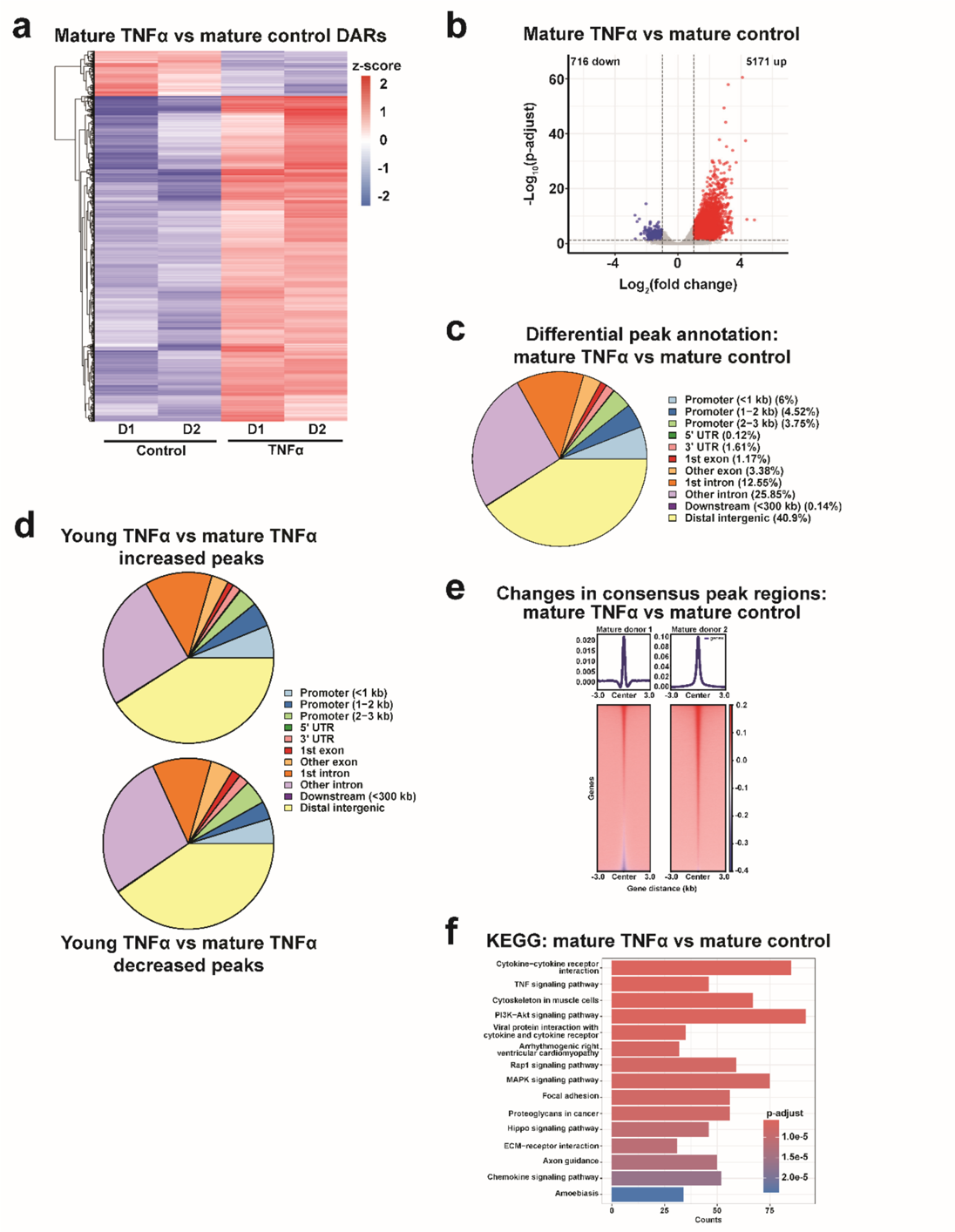
ATAC-seq analysis comparing chromatin accessibility between TNFα-treated and control mature tenocytes. **(a)** Heatmap of DARs between Mature TNFα and Mature Control tenocytes. Color represents z-score–scaled accessibility. **(b)** Volcano plot of all DARs between Mature TNFα and Mature Control. Peaks with significantly increased (red) or decreased (blue) accessibility are indicated. **(c)** Gene annotation of DARs comparing Mature TNFα and Mature Control groups. **(d)** Gene annotation results for increased (top) and decreased (bottom) peaks between Mature TNFα and Mature Control. **(e)** Differential peak enrichment plot of consensus peak regions between Mature TNFα and Mature Control. Color indicates accessibility changes. **(f)** KEGG pathway enrichment analysis of TSS-associated DARs comparing Mature TNFα and Mature Control tenocytes. Color denotes statistical significance. (n=2 biological replicates per condition, TNFα treated for 6 hours, significance: |Log_2_(fold change)|>1 and p-adj<0.05).

**Supplemental Figure S8:**
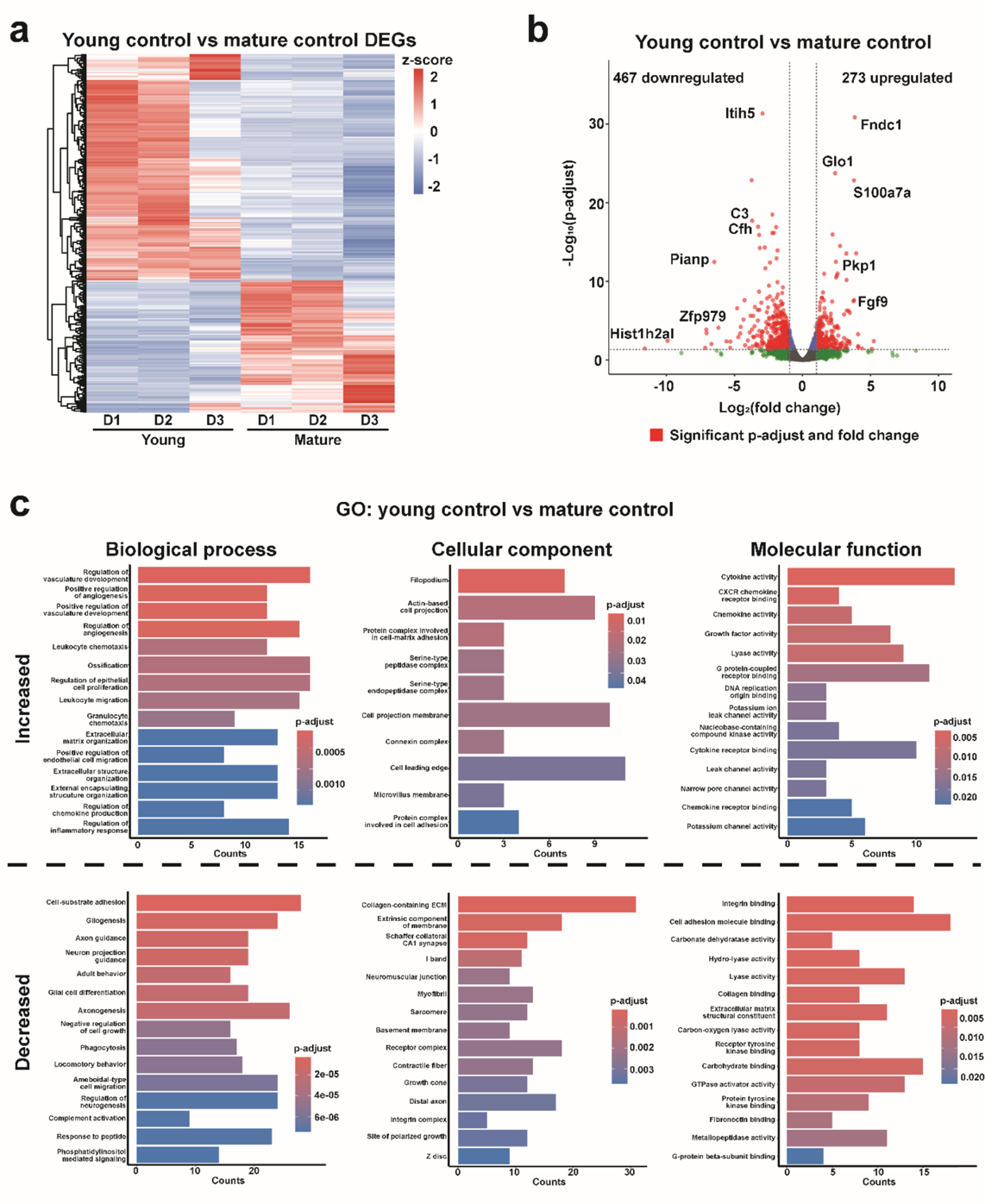
RNA-seq analysis comparing gene expression between young and mature control tenocytes. **(a)** Heatmap of differentially expressed genes (DEGs) comparing Young Control and Mature Control tenocytes. Color represents z-score–scaled expression levels. **(b)** Volcano plot of DEGs between Young Control and Mature Control groups, with significantly altered genes indicated in red. **(c)** Gene Ontology (GO) enrichment analysis of DEGs comparing Young Control and Mature Control tenocytes, categorized by Biological Process (left), Cellular Component (middle), and Molecular Function (right). Terms with increased (top) or decreased (bottom) prevalence in young control cells are colored by statistical significance. (n=3 biological replicates, significance: |Log_2_(fold change)|>1 and p-adj<0.05).

**Supplemental Figure S9:**
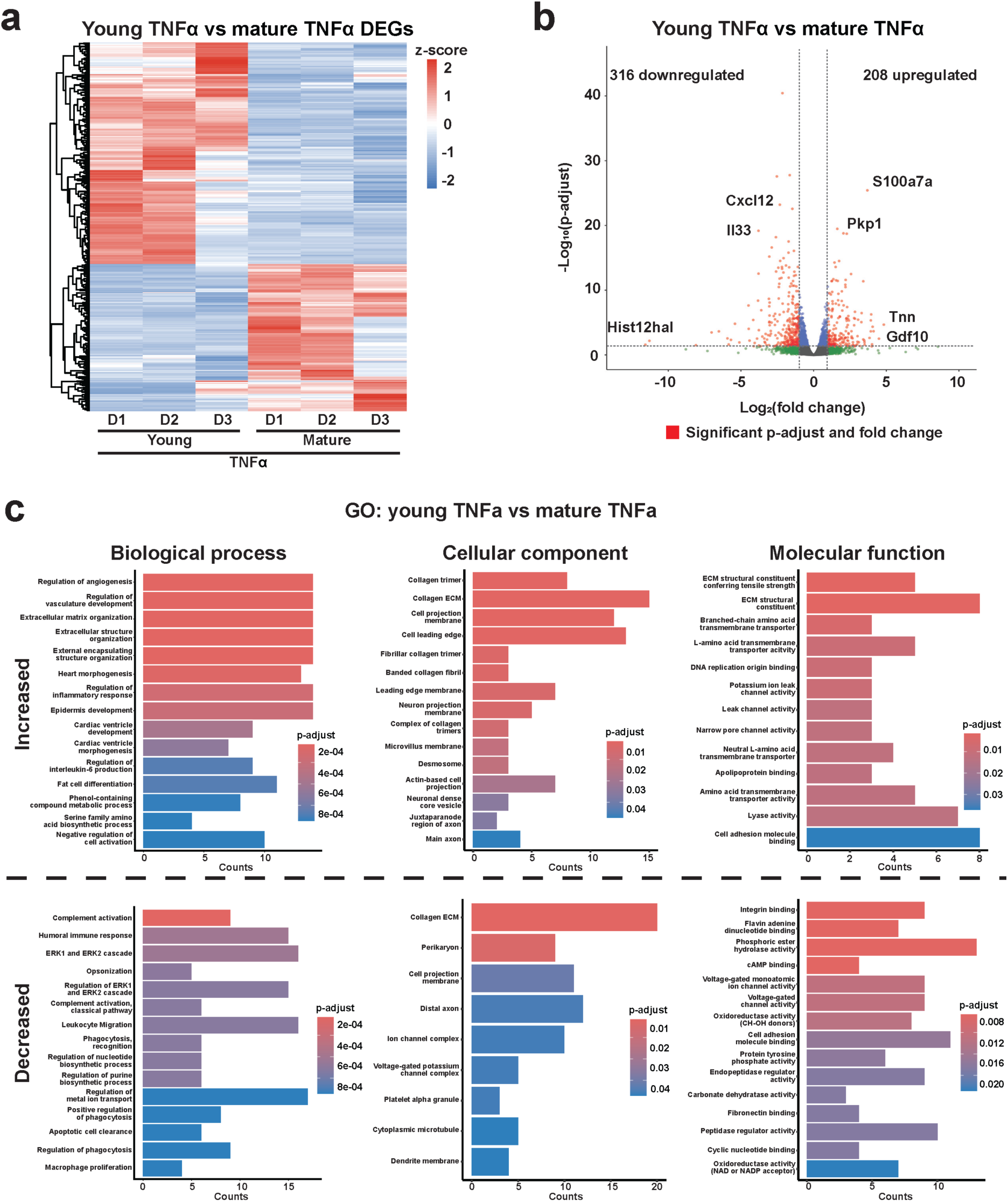
RNA-seq analysis comparing gene expression between young and mature tenocytes under TNFα stimulation. **(a)** Heatmap of DEGs comparing Young TNFα and Mature TNFα tenocytes. Color represents z-score–scaled expression levels. **(b)** Volcano plot of DEGs between Young TNFα and Mature TNFα groups, with significantly altered genes highlighted in red. **(c)** GO enrichment analysis of DEGs comparing Young TNFα and Mature TNFα tenocytes, categorized by Biological Process (left), Cellular Component (middle), and Molecular Function (right). Terms with increased (top) or decreased (bottom) prevalence in young TNFα-treated cells are colored according to statistical significance. (n=3 biological replicates, TNFα treated for 6 hours, significance: |Log_2_(fold change)|>1 and p-adj<0.05).

**Supplemental Figure S10:**
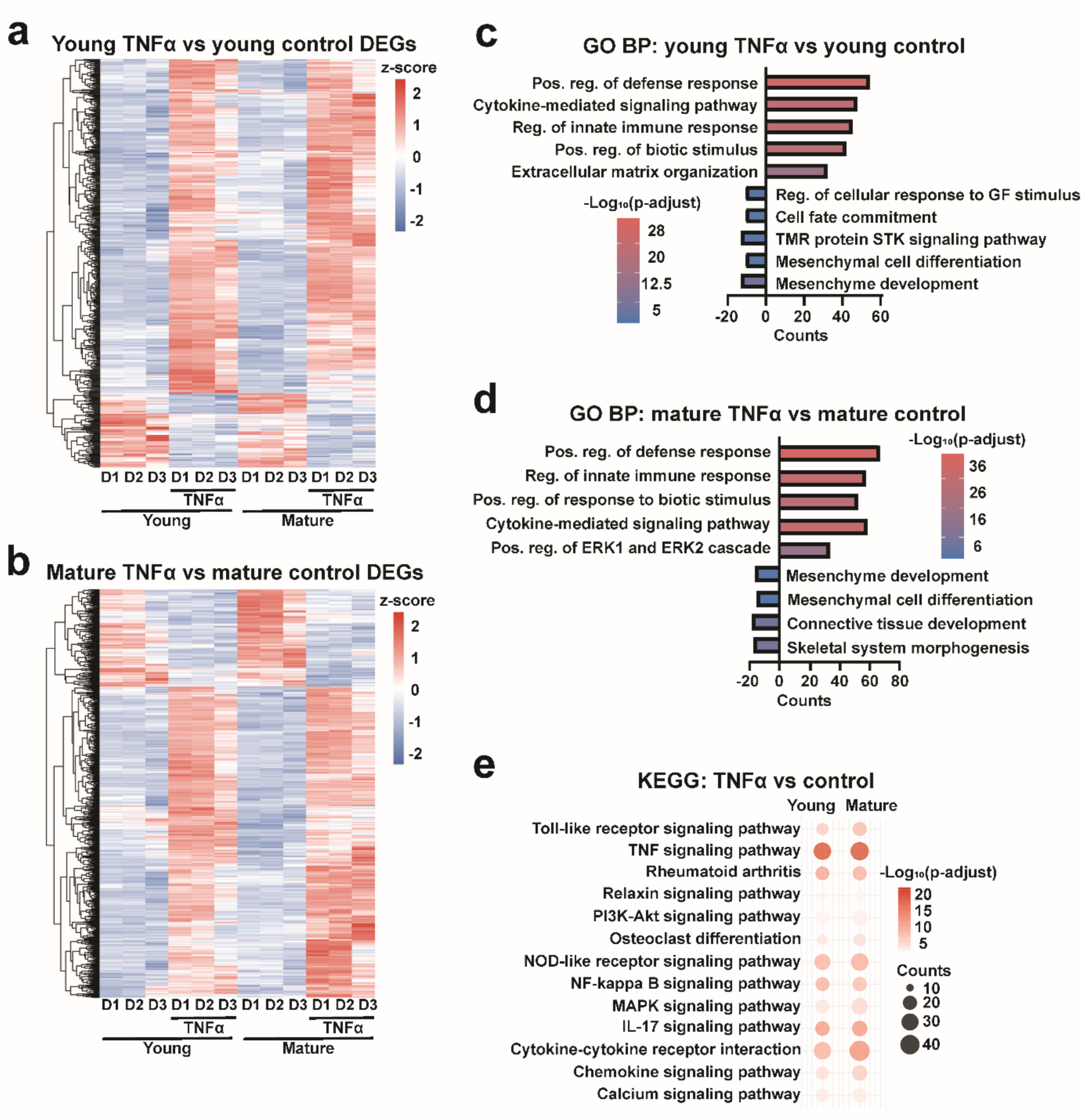
RNA-seq analysis comparing TNFα-treated and control tenocytes across age groups. (a, **b)** Heatmaps of DEGs comparing Young TNFα versus Young Control **(a)** and Mature TNFα versus Mature Control **(b)** groups. Color represents z-score–scaled expression levels. **(c, d)** GO enrichment analysis of DEGs within the Biological Process (BP) category for Young TNFα versus Young Control **(c)** and Mature TNFα versus Mature Control **(d)** comparisons. Color denotes significance, and the direction of counts indicates gene upregulation (positive) or downregulation (negative). **(e)** KEGG pathway enrichment analysis of DEGs comparing TNFα-treated to control tenocytes in young (left) and mature (right) groups. Color indicates significance, and circle size represents gene count. (n=3 biological replicates, TNFα treated for 6 hours, significance: |Log_2_(fold change)|>1 and p-adj<0.05). **(a, b)** Predicted upstream regulators identified by Ingenuity Pathway Analysis (IPA) from age-exclusive TNFα-induced DEGs overlapping with the GO term “Chromatin Remodelers.” Results are shown for young **(a)** and mature **(b)** tenocytes, with color representing statistical significance. **(c)** Heatmap of age-exclusive DEGs within the GO term “Chromatin Organization” for young and mature tenocytes following TNFα stimulation. Color represents z-score–scaled expression levels. (n=3 biological replicates, TNFα treated for 6 hours, significance: |Log_2_(fold change)|>1 and p-adj<0.05).

**Supplemental Figure S11:**
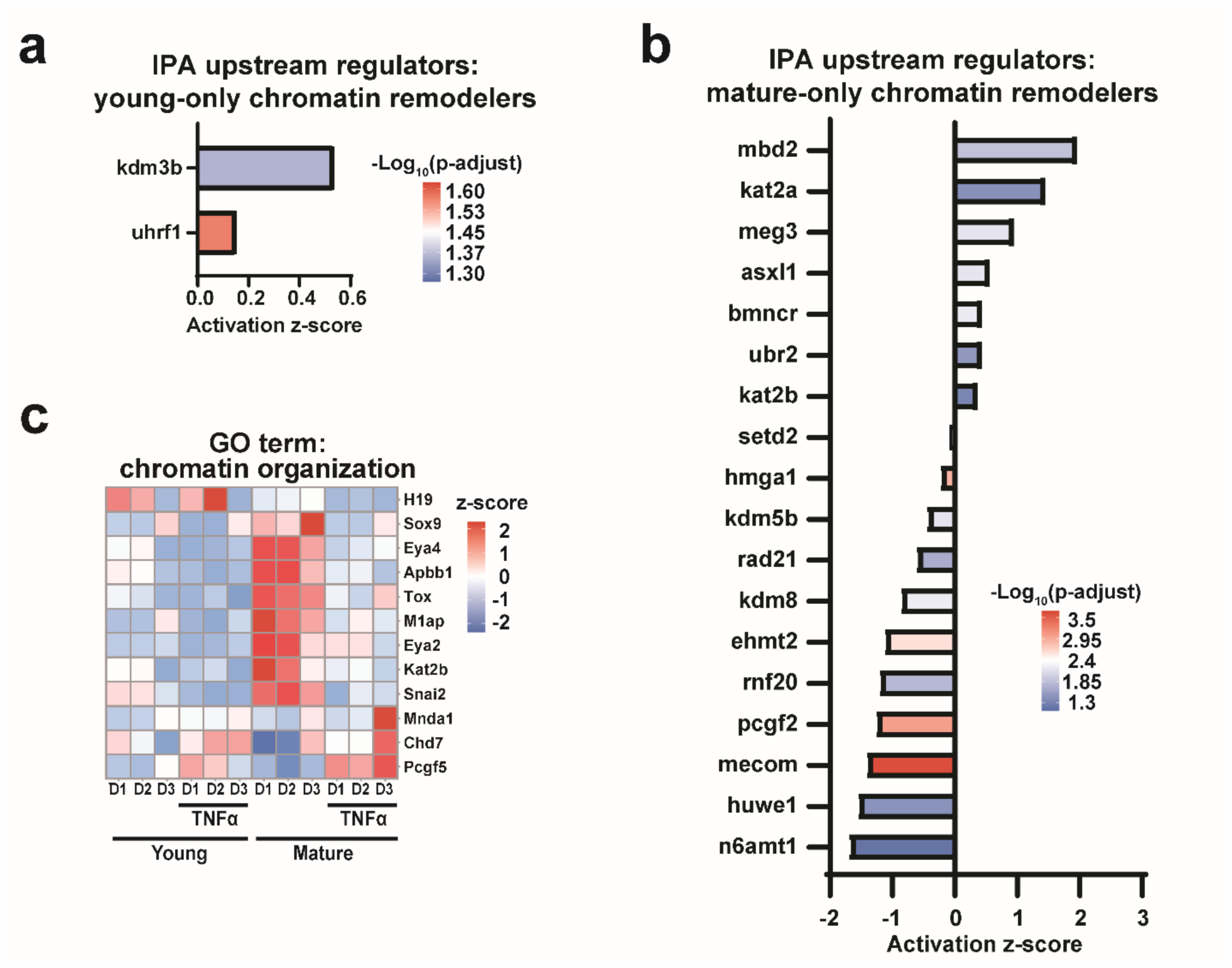
Chromatin state modifiers associated with inflammatory stimulation. (a, **b)** Predicted upstream regulators identified by Ingenuity Pathway Analysis (IPA) from age-exclusive TNFα-induced DEGs overlapping with the GO term “Chromatin Remodelers.” Results are shown for young **(a)** and mature **(b)** tenocytes, with color representing statistical significance. **(c)** Heatmap of age-exclusive DEGs within the GO term “Chromatin Organization” for young and mature tenocytes following TNFα stimulation. Color represents z-score–scaled expression levels. (n=3 biological replicates, TNFα treated for 6 hours, significance: |Log2(fold change)|>1 and p-adj<0.05).

**Supplemental Figure S12:**
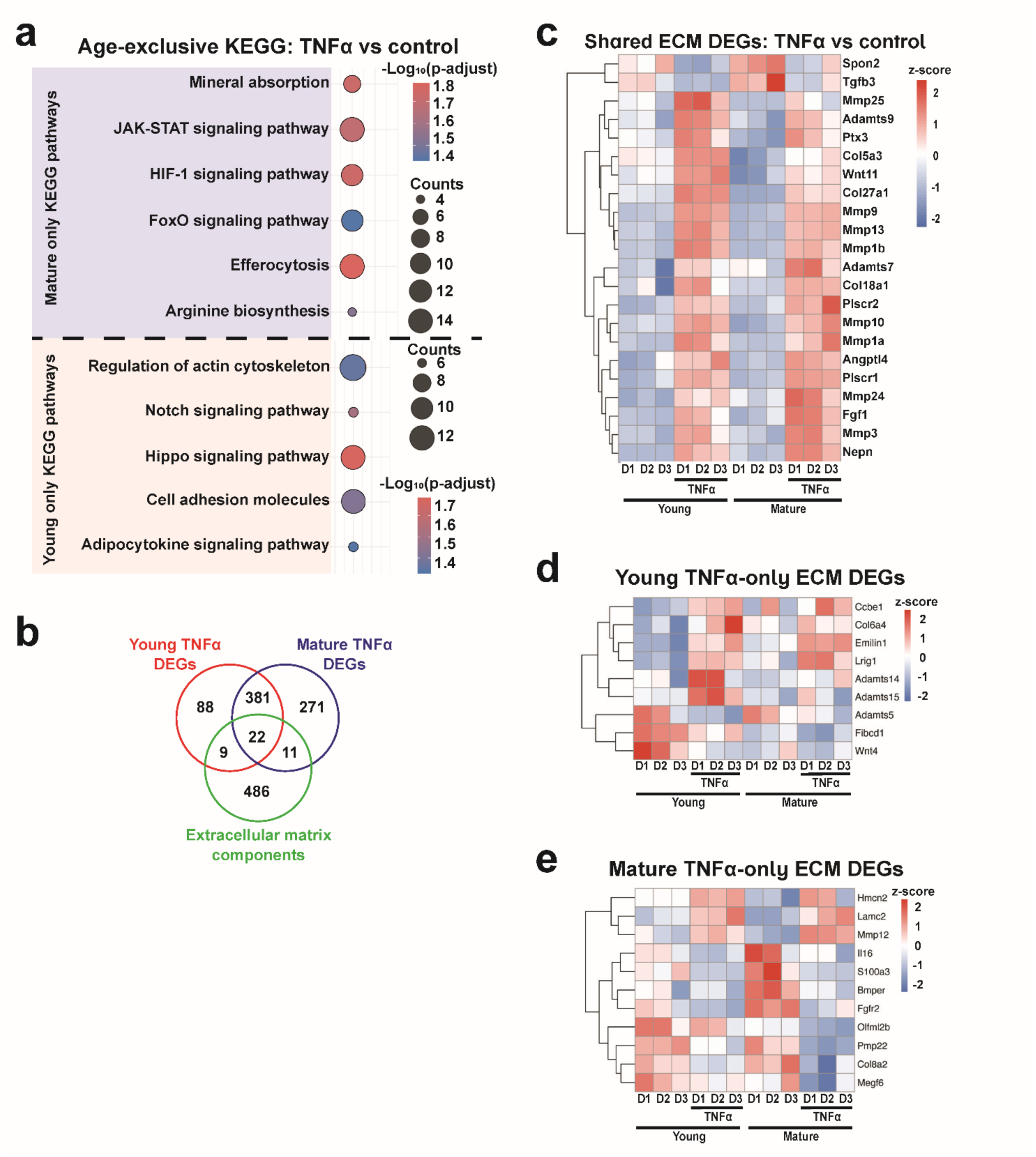
Altered extracellular matrix remodeling following TNFα stimulation in young and mature tenocytes. **(a)** KEGG pathway enrichment analysis of age-exclusive DEGs comparing TNFα-treated and control tenocytes in young and mature groups. Color denotes statistical significance, and circle size represents gene count. Age-exclusive results for young and mature tenocytes are highlighted in red and blue, respectively. **(b)** Diagram showing TNFα-induced DEGs in young (red) and mature (blue) tenocytes overlapping with the GO term “Extracellular Matrix Components and Regulators” (green). **(c-e)** Heatmaps of ECM-related DEGs significantly regulated in both age groups (shared) **(c)**, or exclusive to young **(d)** or mature **(e)** tenocytes following TNFα treatment. Color represents z-score–scaled expression. (n=3 biological replicates, TNFα treated for 6 hours, significance: |Log_2_(fold change)|>1 and p-adj<0.05).

**Table S1:**
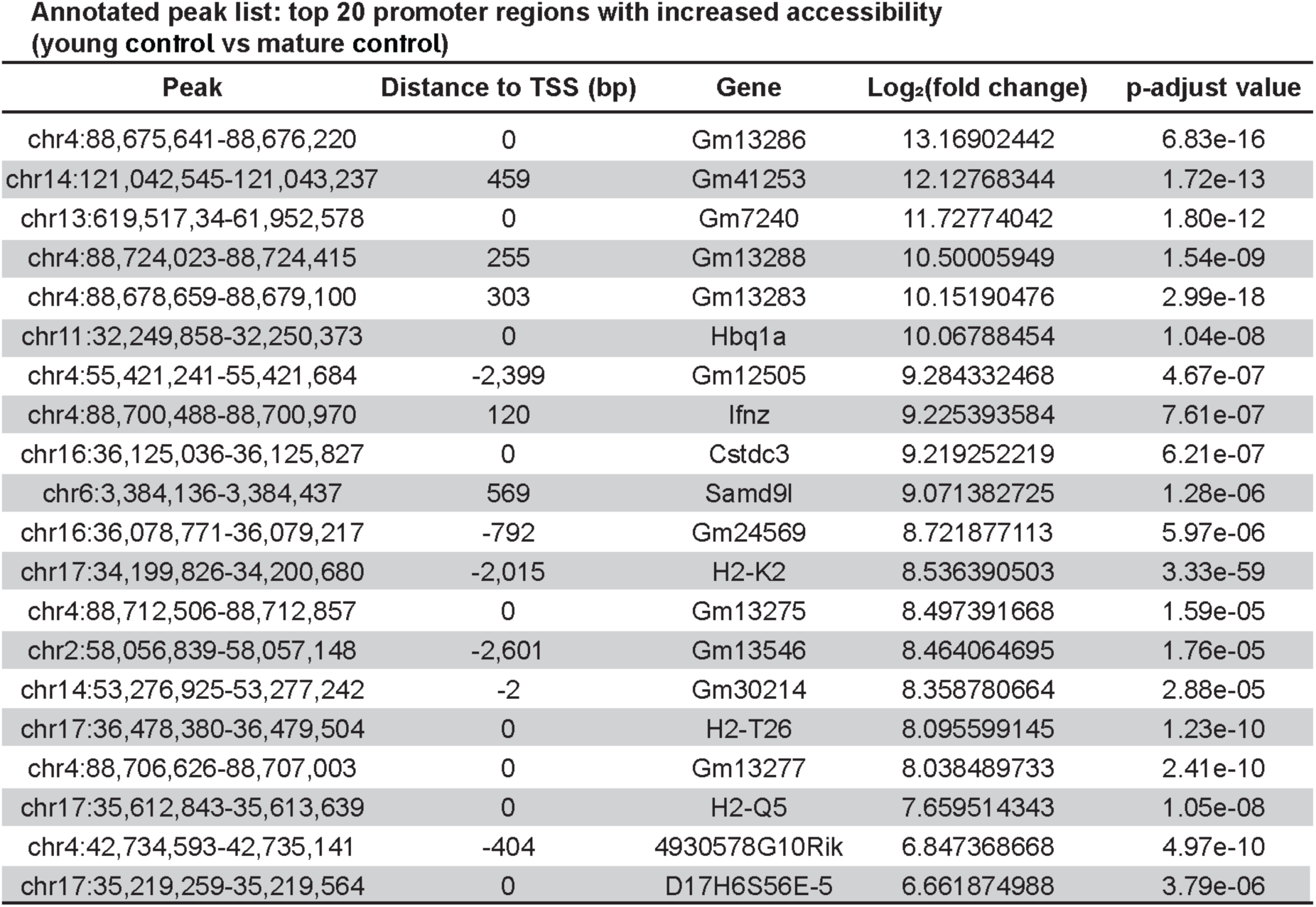
Top 20 transcription start site-associated (TSS-associated) differentially accessible regions (DARs) with increased accessibility in Young Control compared to Mature Control tenocytes.

**Table S2:**
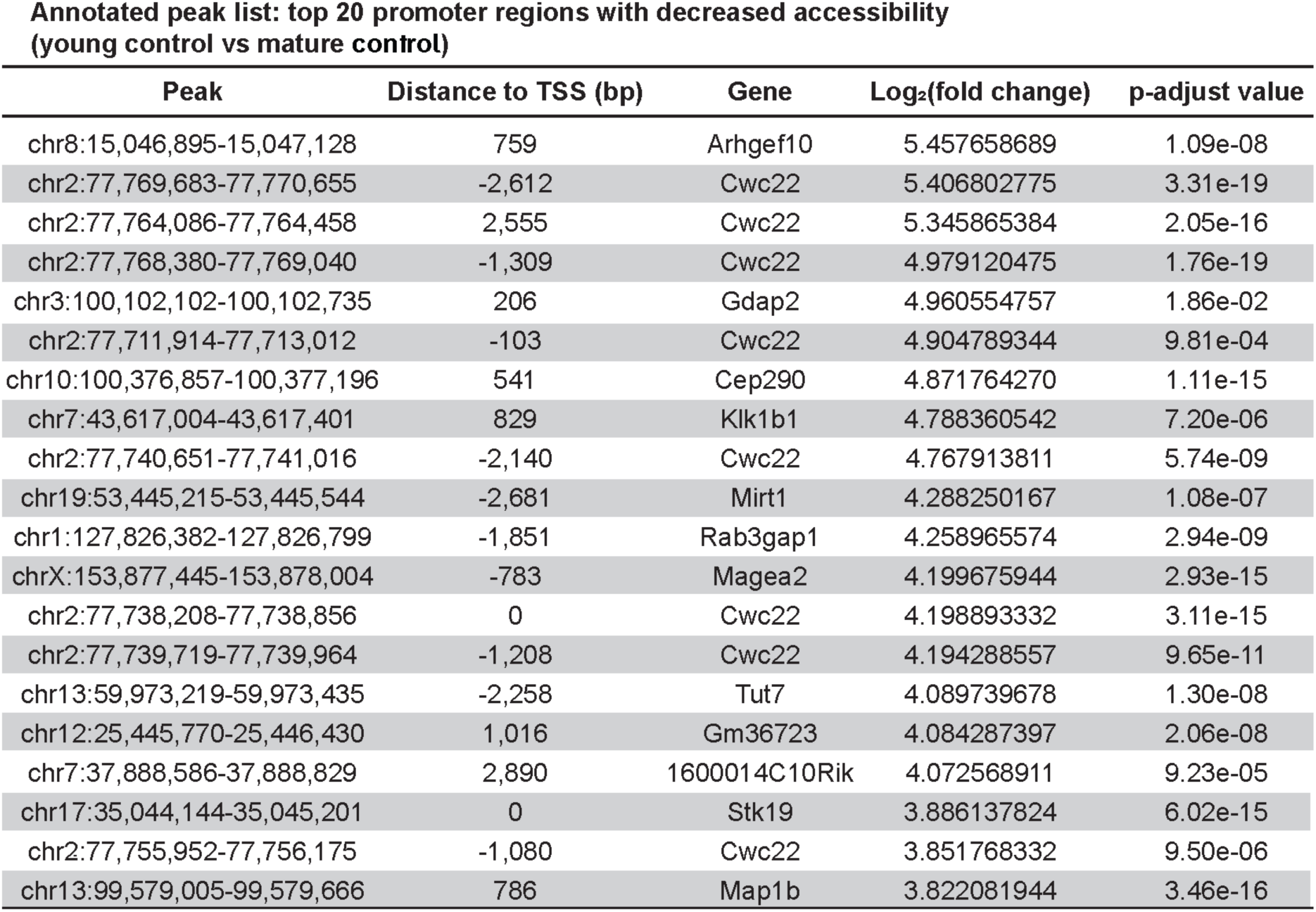
Top 20 TSS-associated DARs with decreased accessibility in Young Control compared to Mature Control tenocytes.

**Table S3:**
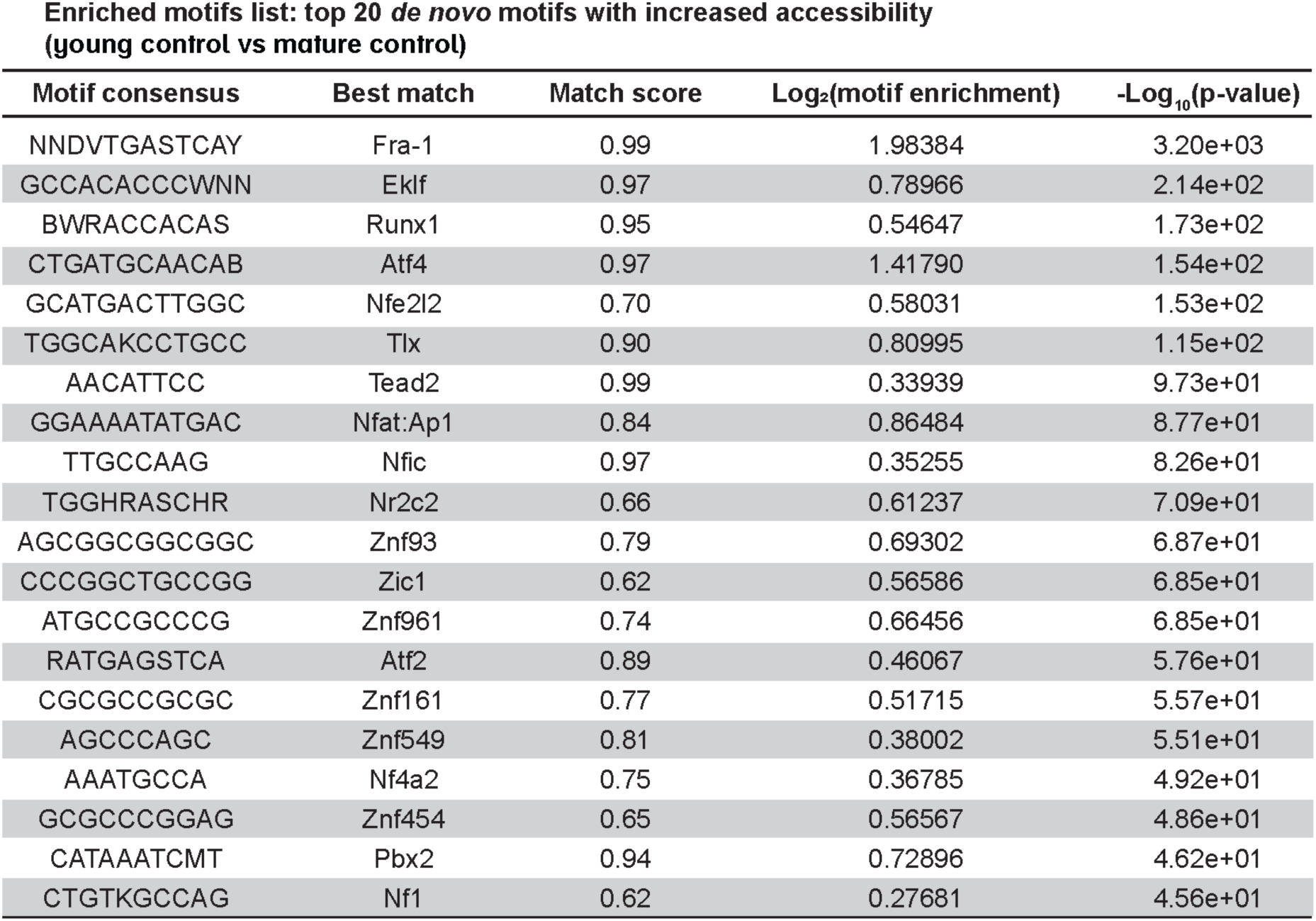
Top 20 differentially accessible transcription factor motifs with increased accessibility in Young Control compared to Mature Control tenocytes.

**Table S4:**
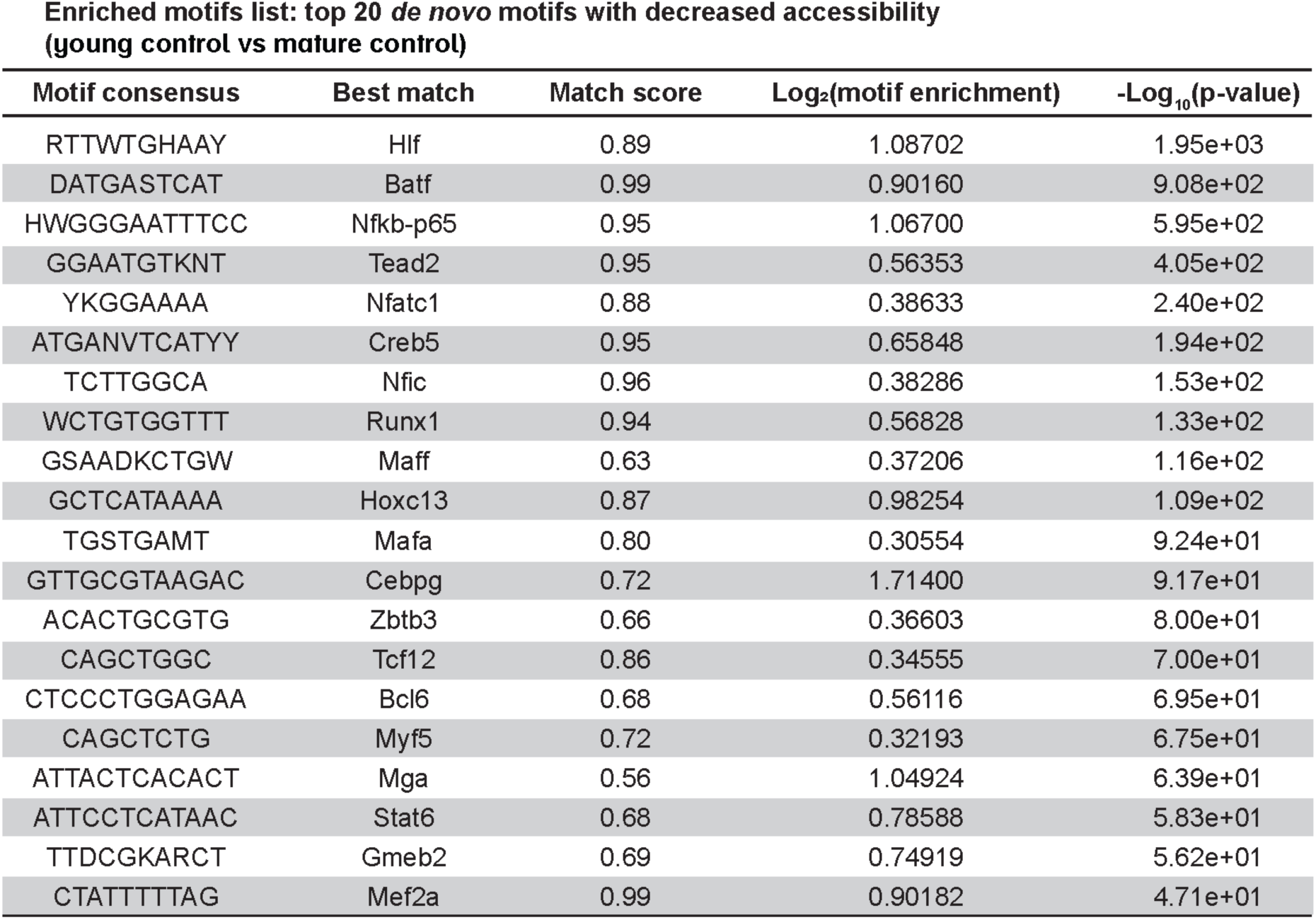
Top 20 differentially accessible transcription factor motifs with decreased accessibility in Young Control compared to Mature Control tenocytes.

**Table S5:**
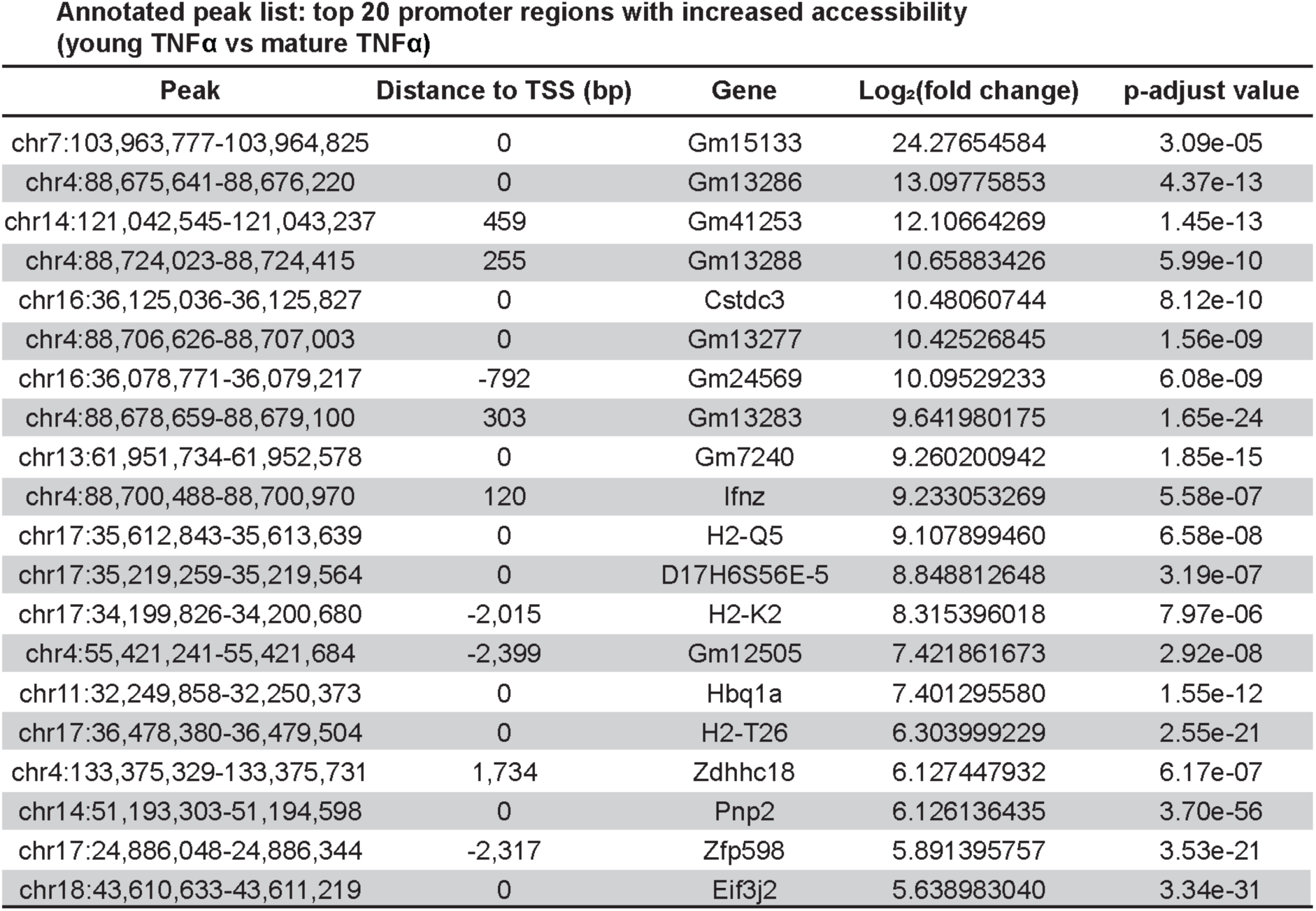
Top 20 TSS-associated DARs with increased accessibility in Young TNFα compared to Mature TNFα tenocytes.

**Table S6:**
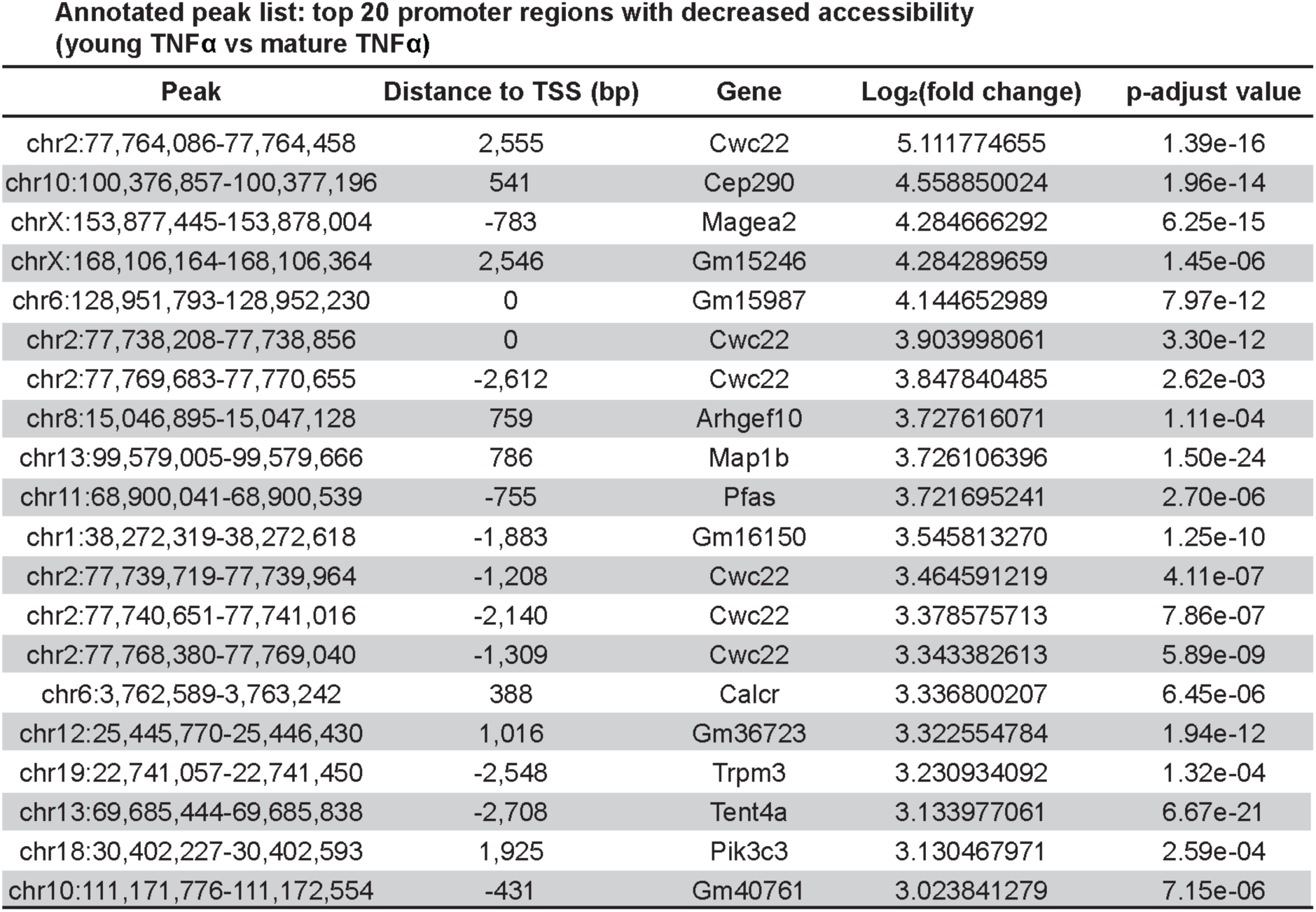
Top 20 TSS-associated DARs with decreased accessibility in Young TNFα compared to Mature TNFα tenocytes.

**Table S7:**
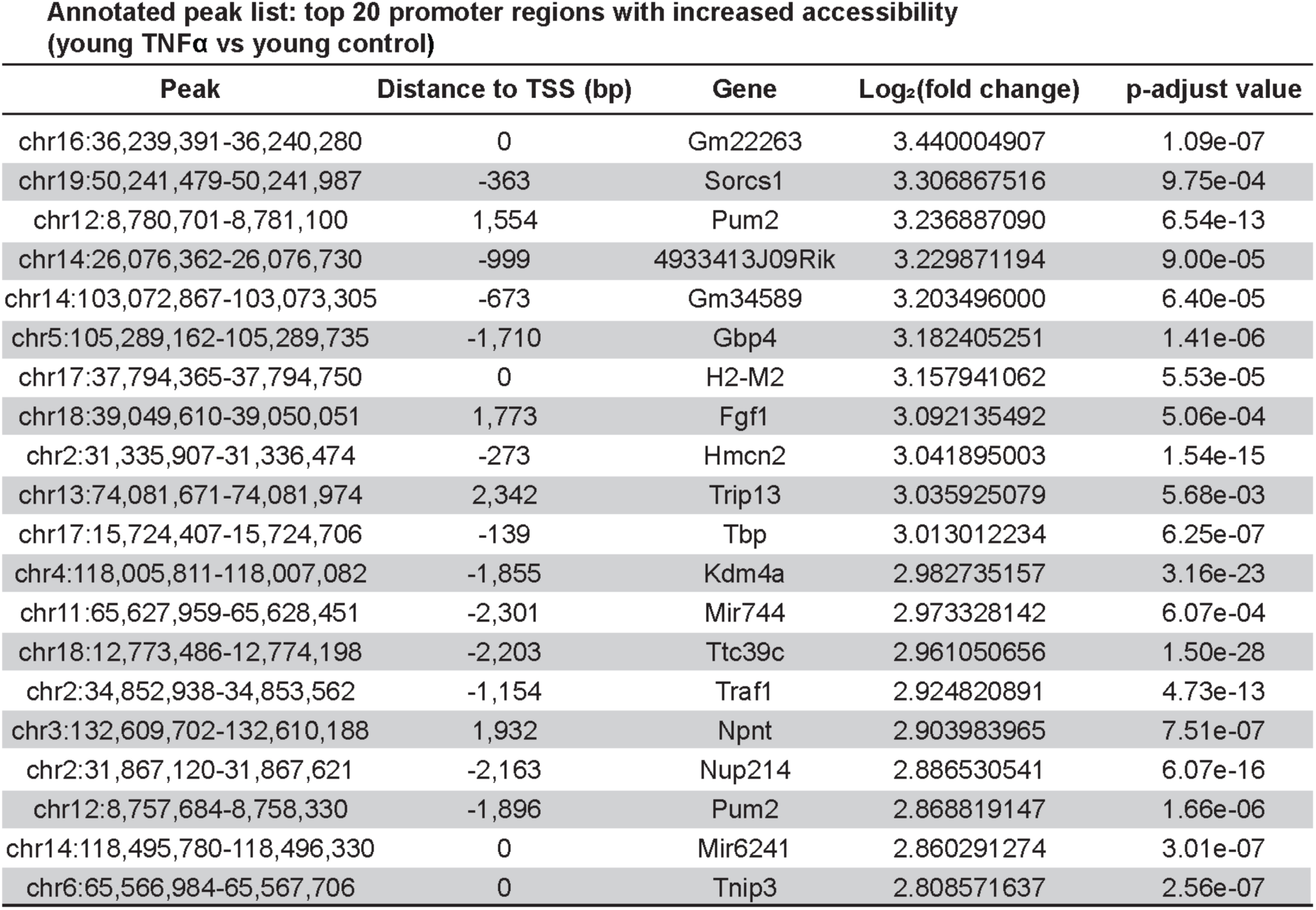
Top 20 TSS-associated DARs with increased accessibility in Young TNFα compared to Young Control tenocytes.

**Table S8:**
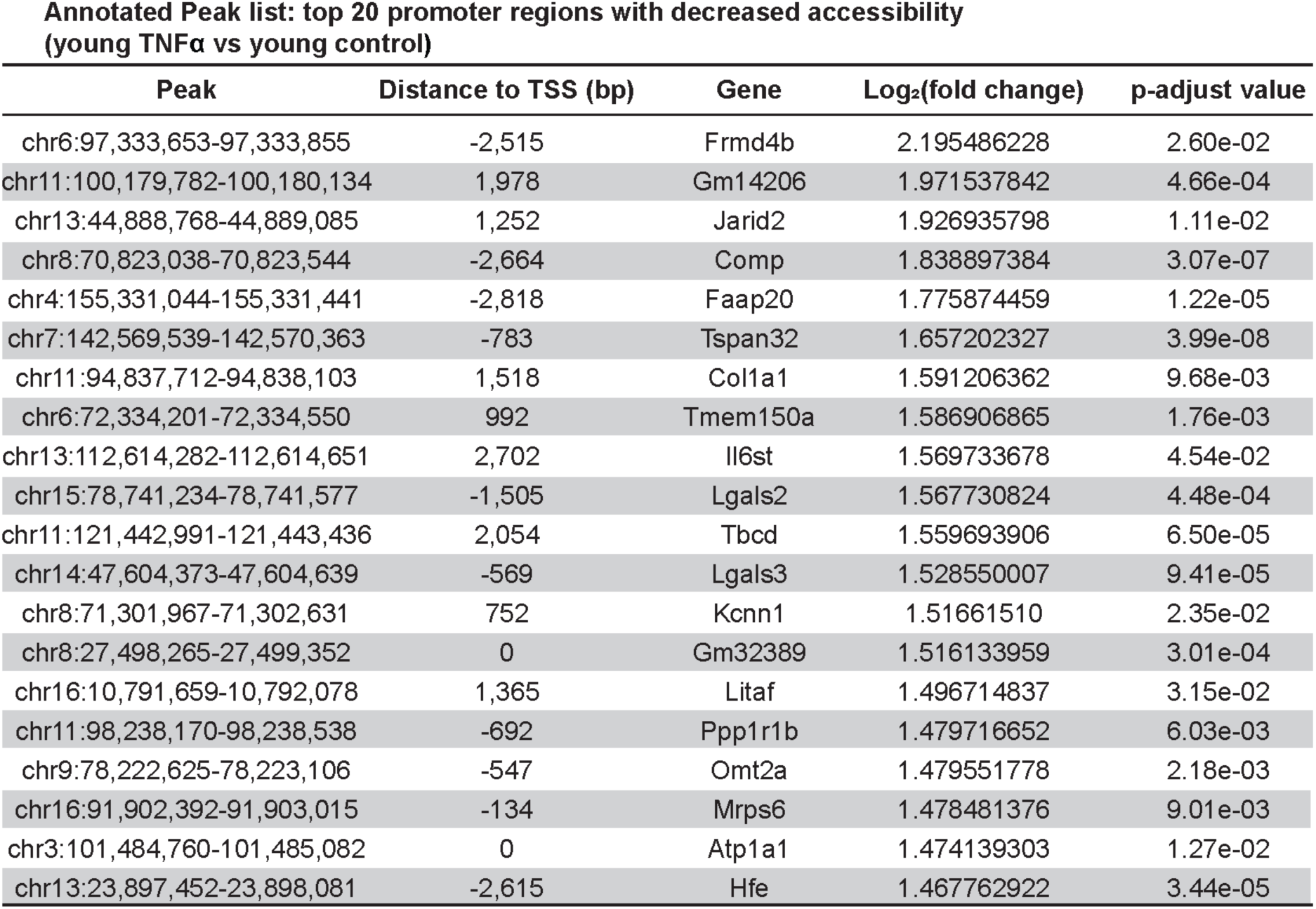
Top 20 TSS-associated DARs with decreased accessibility in Young TNFα compared to Young Control tenocytes.

**Table S9:**
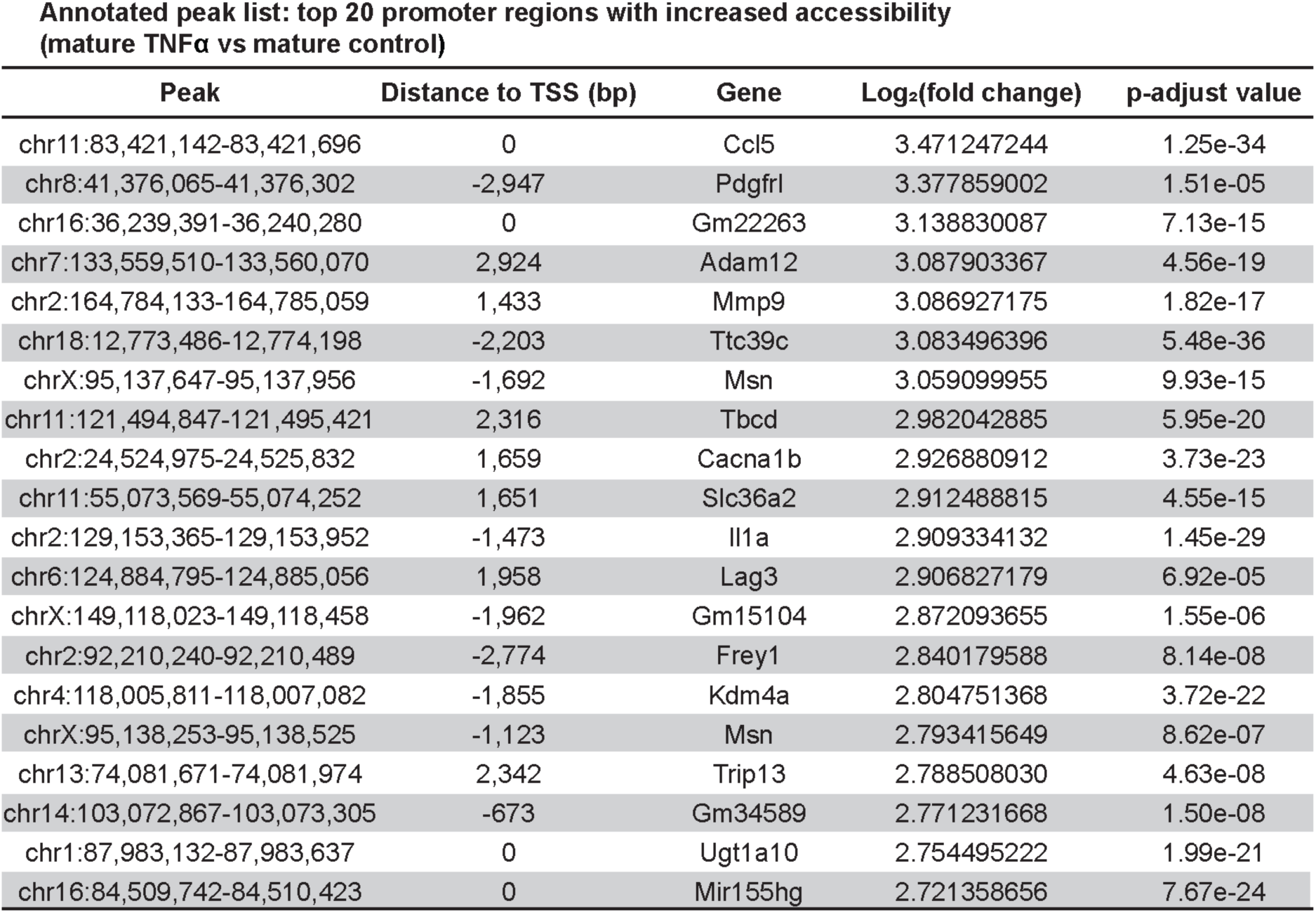
Top 20 TSS-associated DARs with increased accessibility in Mature TNFα compared to Mature Control tenocytes.

**Table S10:**
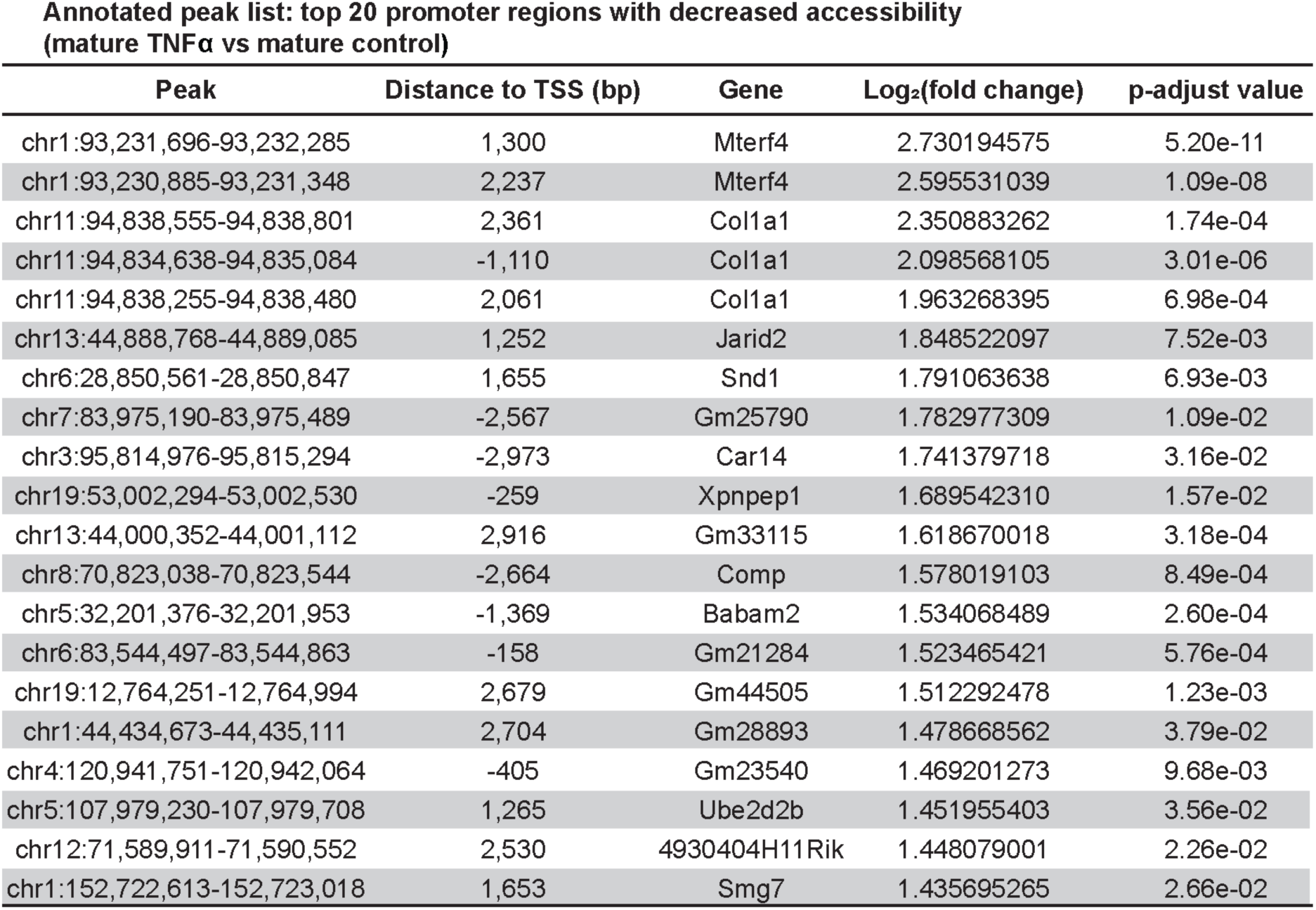
Top 20 TSS-associated DARs with decreased accessibility in Mature TNFα compared to Mature Control tenocytes.

**Table S11:**
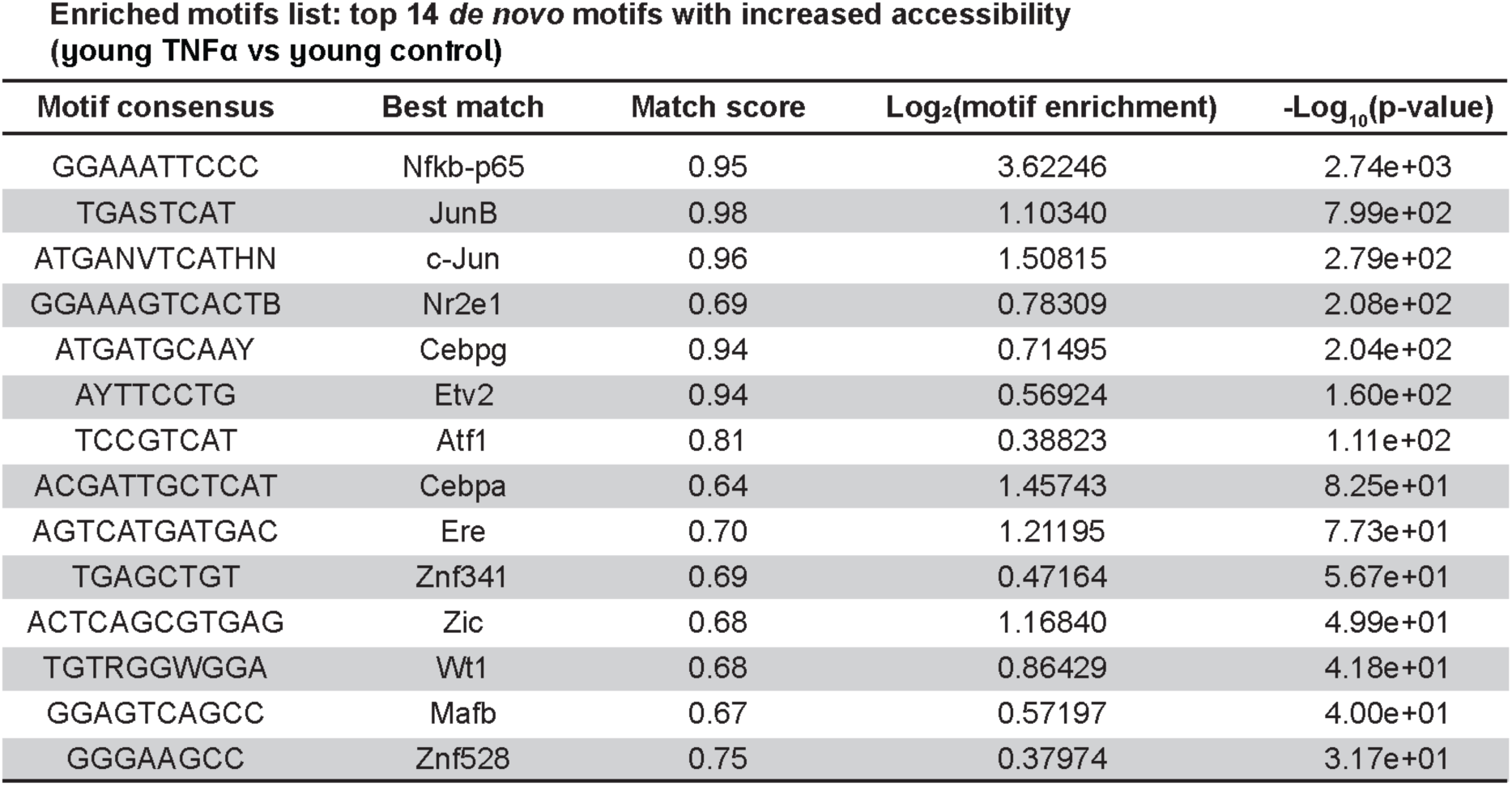
Top 14 differentially accessible transcription factor motifs with increased accessibility in Young TNFα compared to Young Control tenocytes.

**Table S12:**
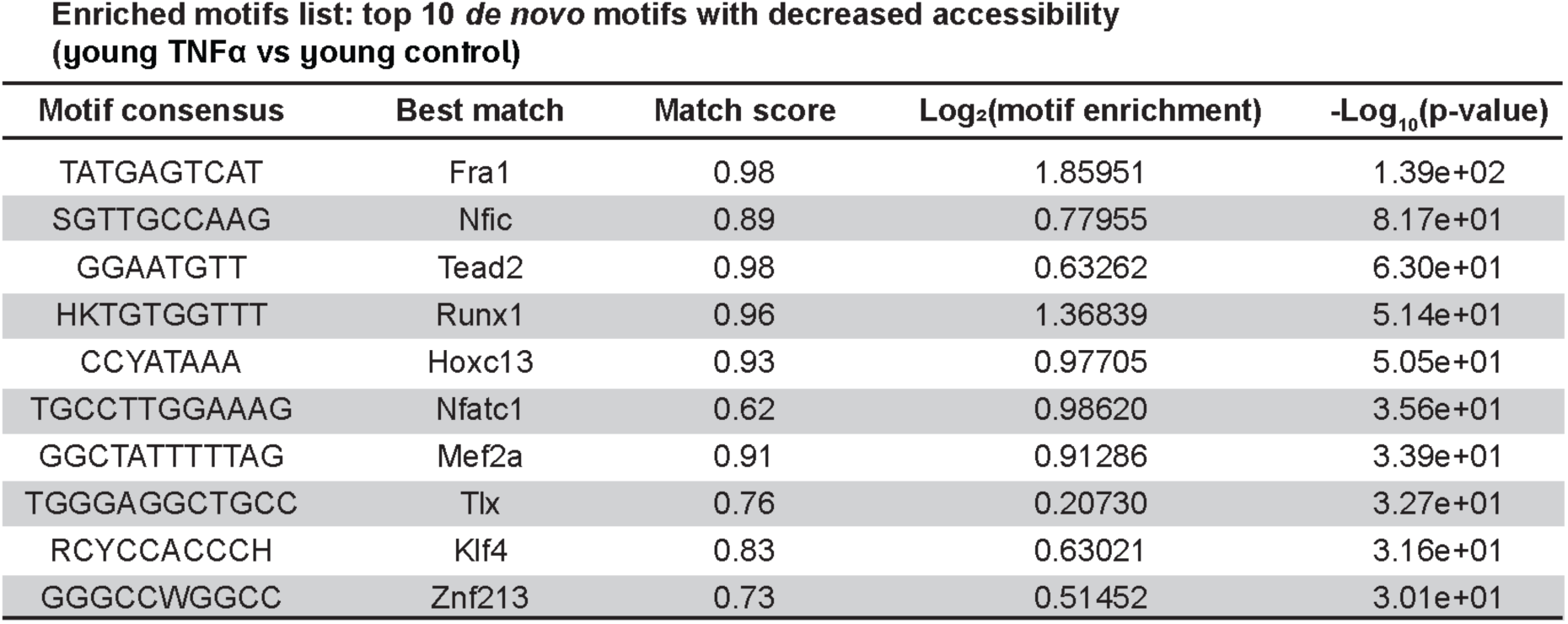
Top 10 differentially accessible transcription factor motifs with decreased accessibility in Young TNFα compared to Young Control tenocytes.

**Table S13:**
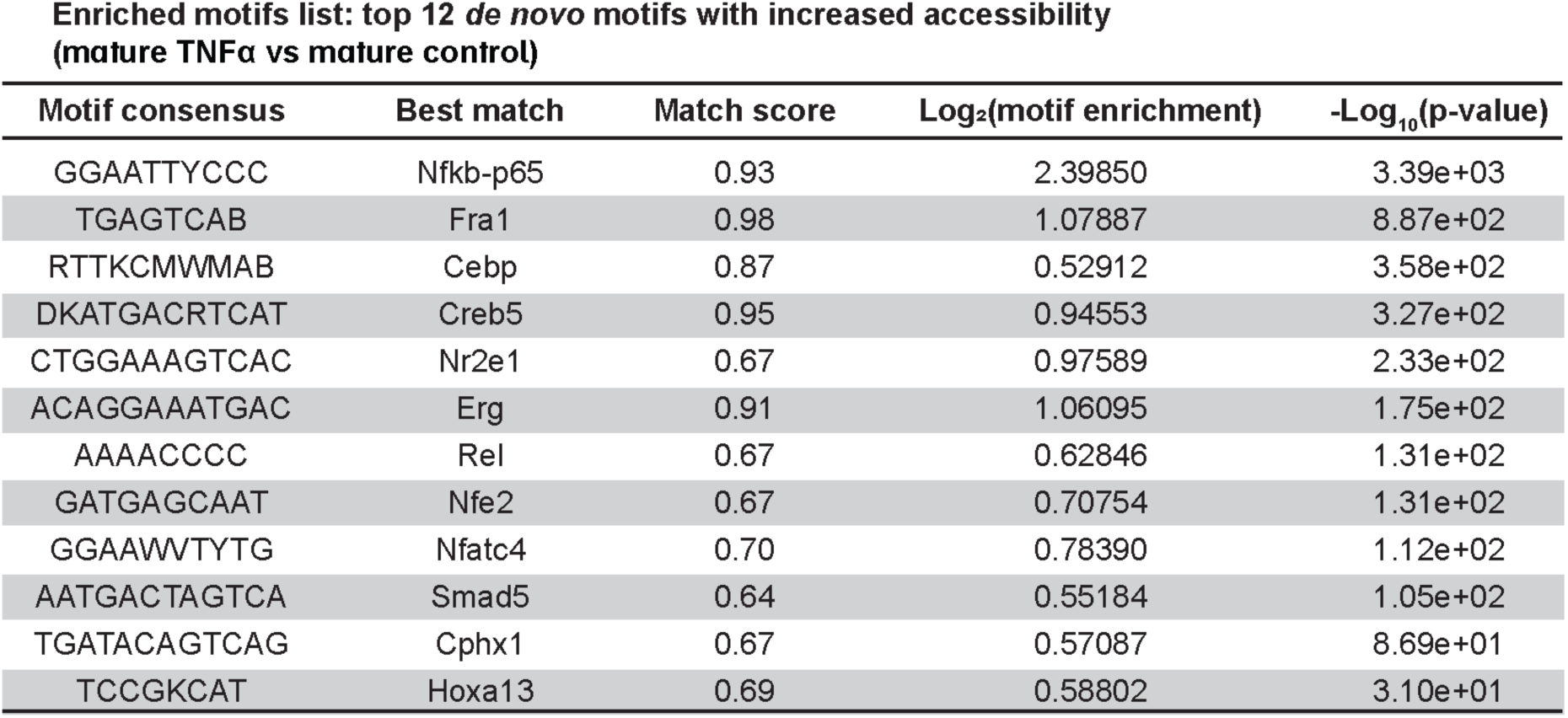
Top 12 differentially accessible transcription factor motifs with increased accessibility in Mature TNFα compared to Mature Control tenocytes.

**Table S14:**
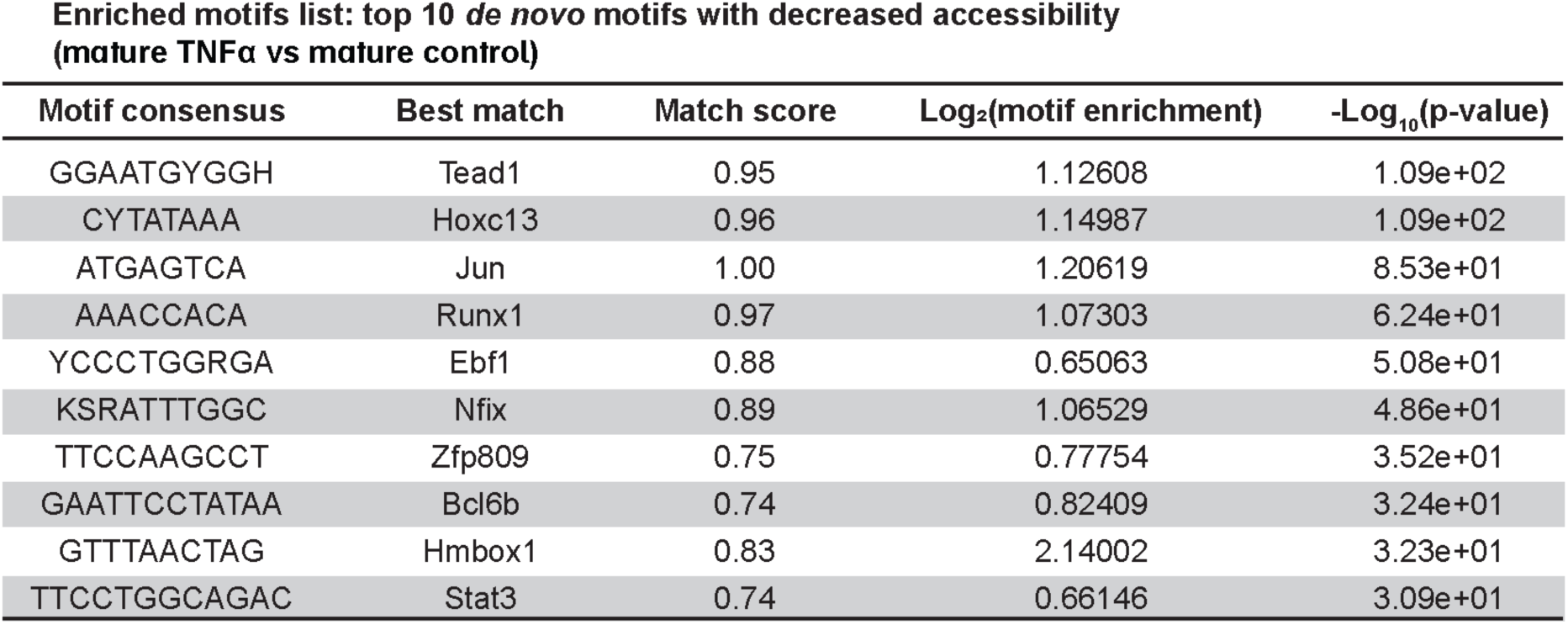
Top 10 differentially accessible transcription factor motifs with decreased accessibility in Mature TNFα compared to Mature Control tenocytes.

**Table S15:**
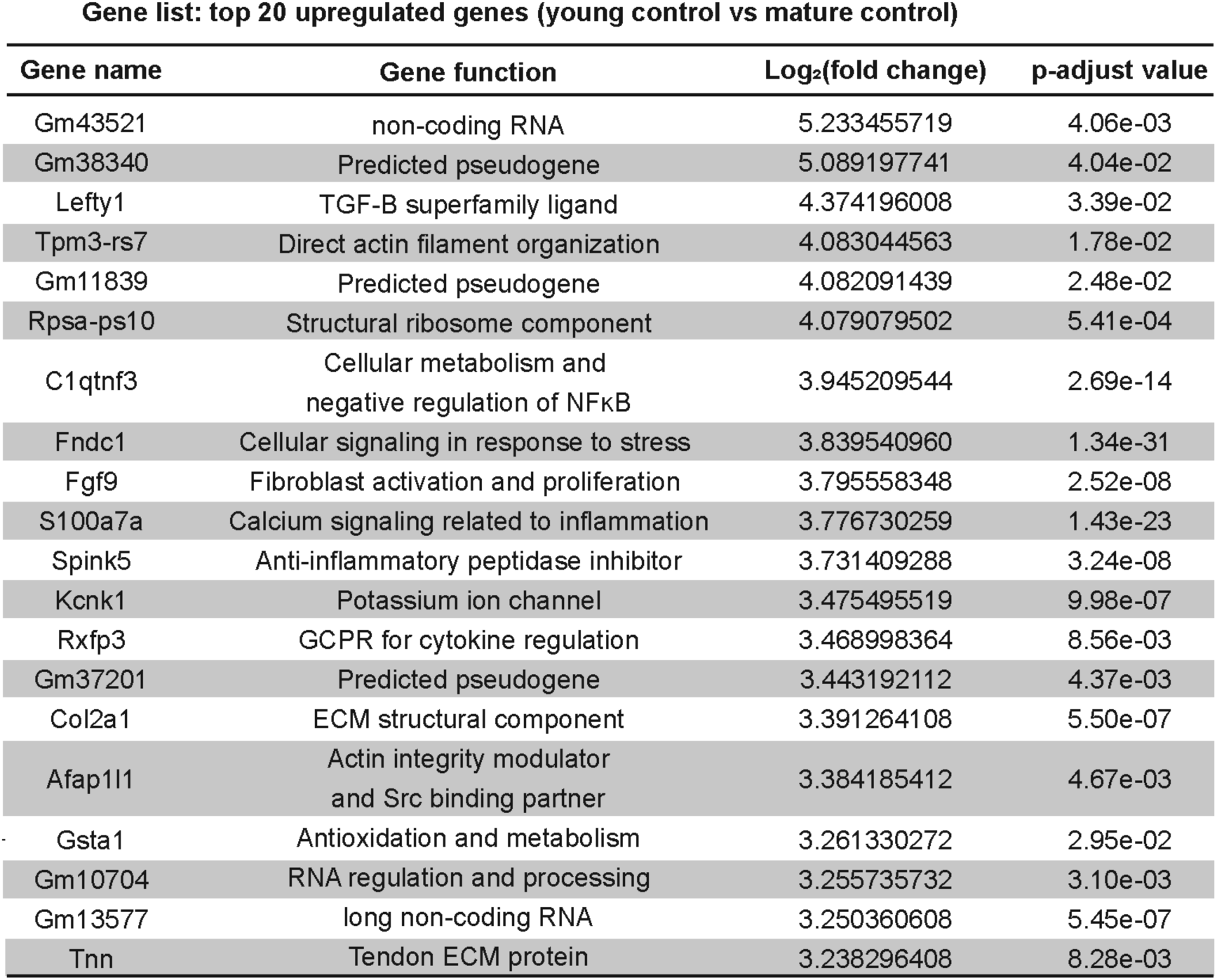
Top 20 differentially expressed genes with increased expression in Young Control compared to Mature Control.

**Table S16:**
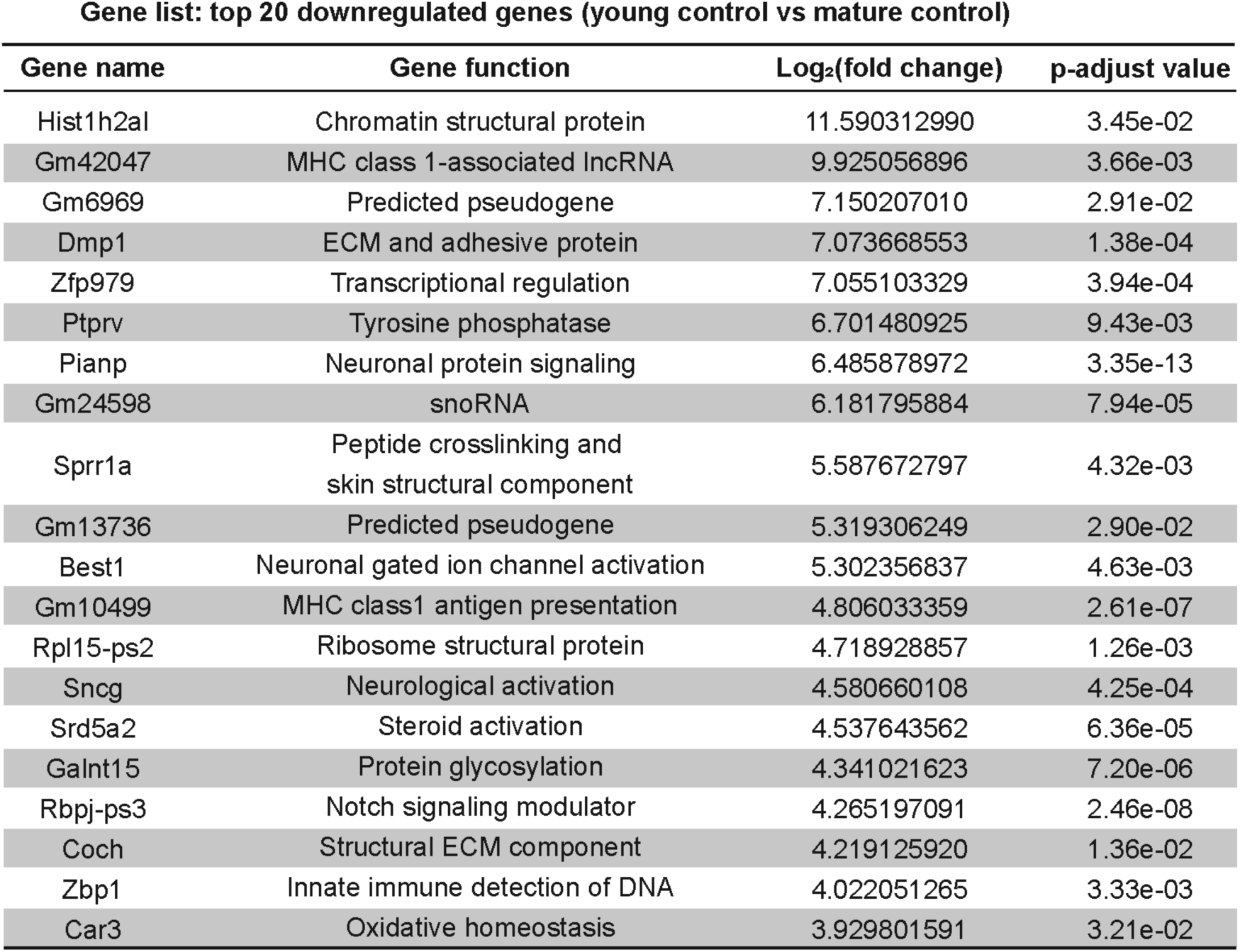
Top 20 differentially expressed genes with decreased expression in Young Control compared to Mature Control.

**Table S17:**
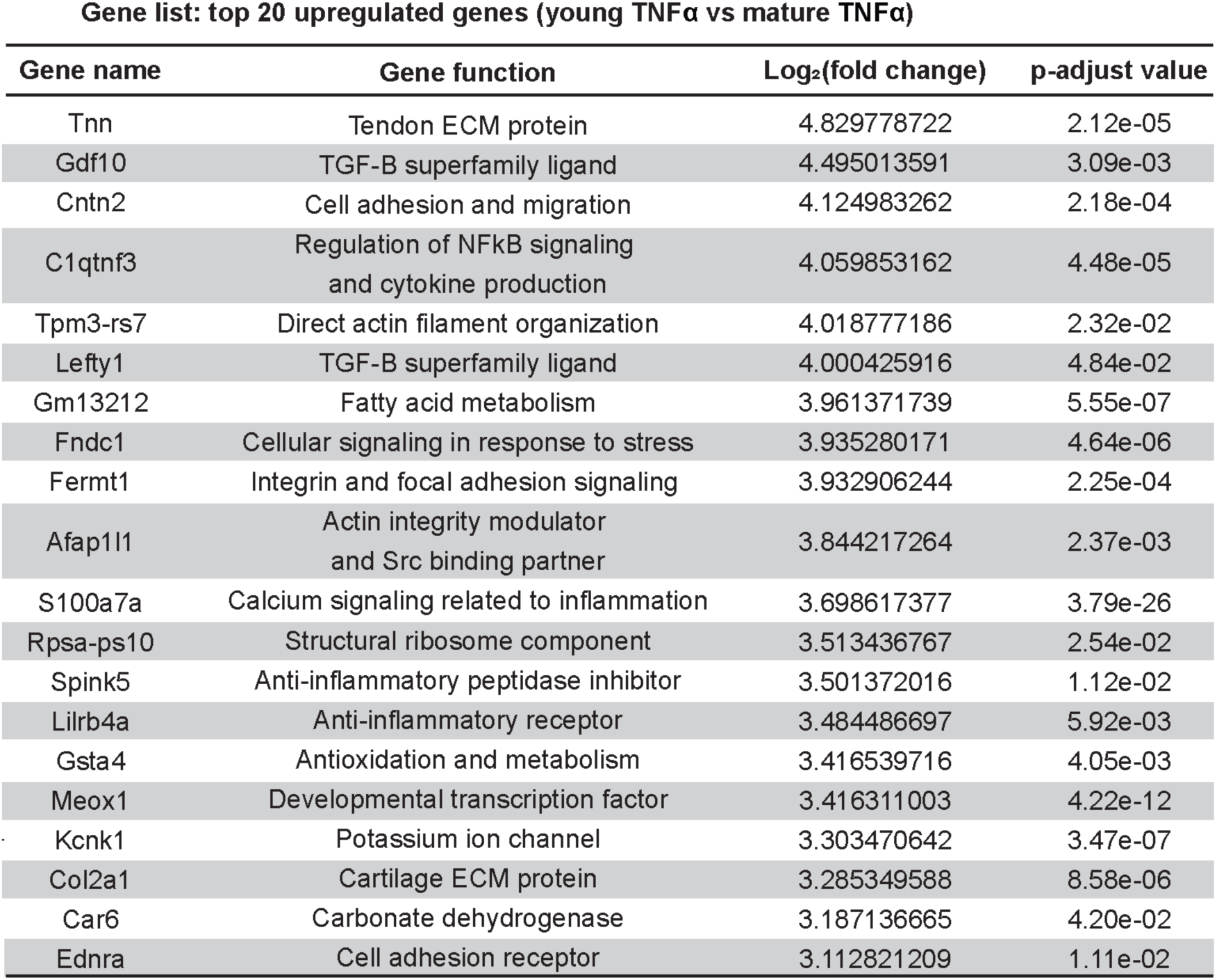
Top 20 differentially expressed genes with increased expression in Young TNFα compared to Mature TNFα.

**Table S18:**
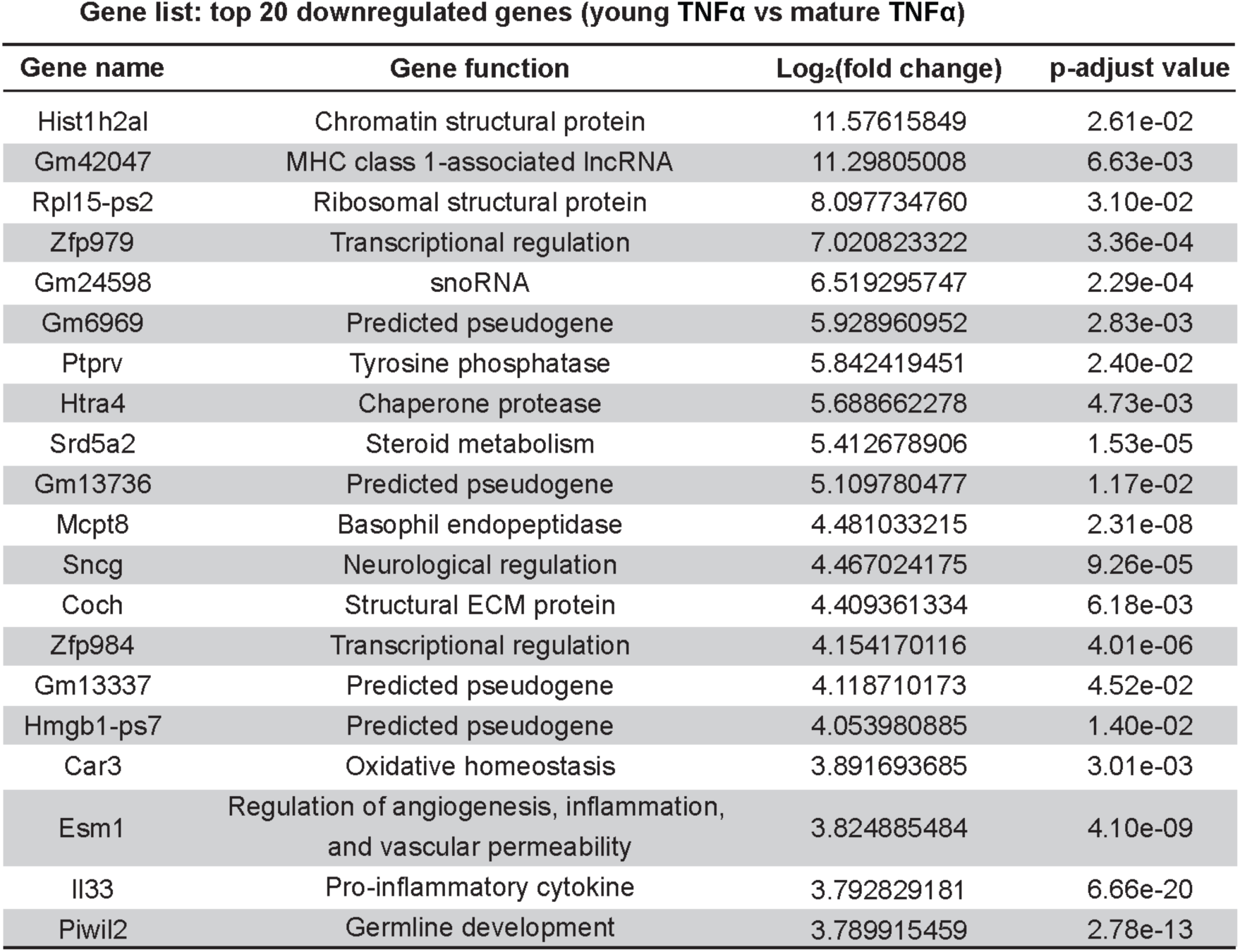
Top 20 differentially expressed genes with decreased expression in Young TNFα compared to Mature TNFα.

**Table S19:**
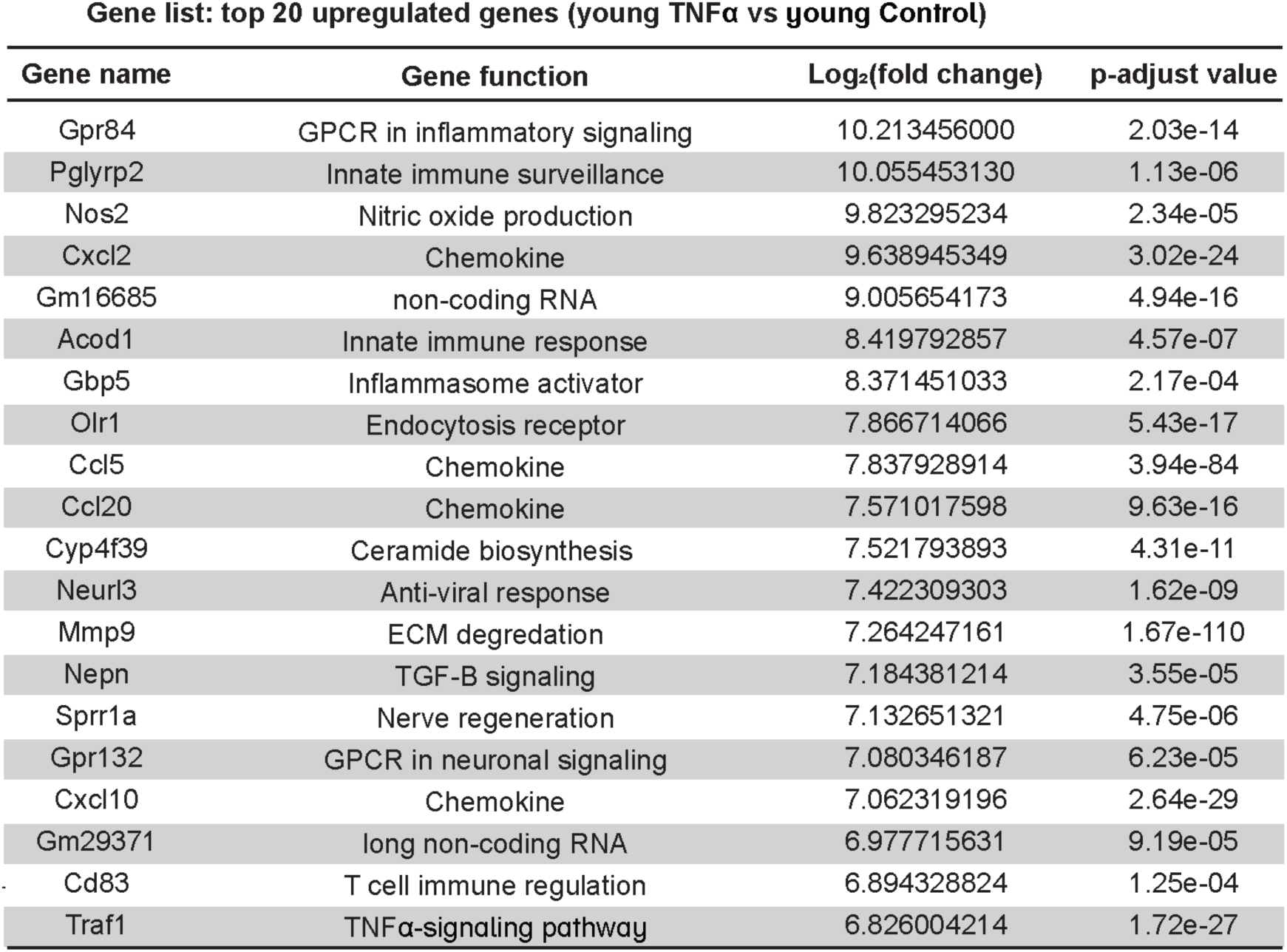
Top 20 differentially expressed genes with increased expression in Young TNFα compared to Young Control.

**Table S20:**
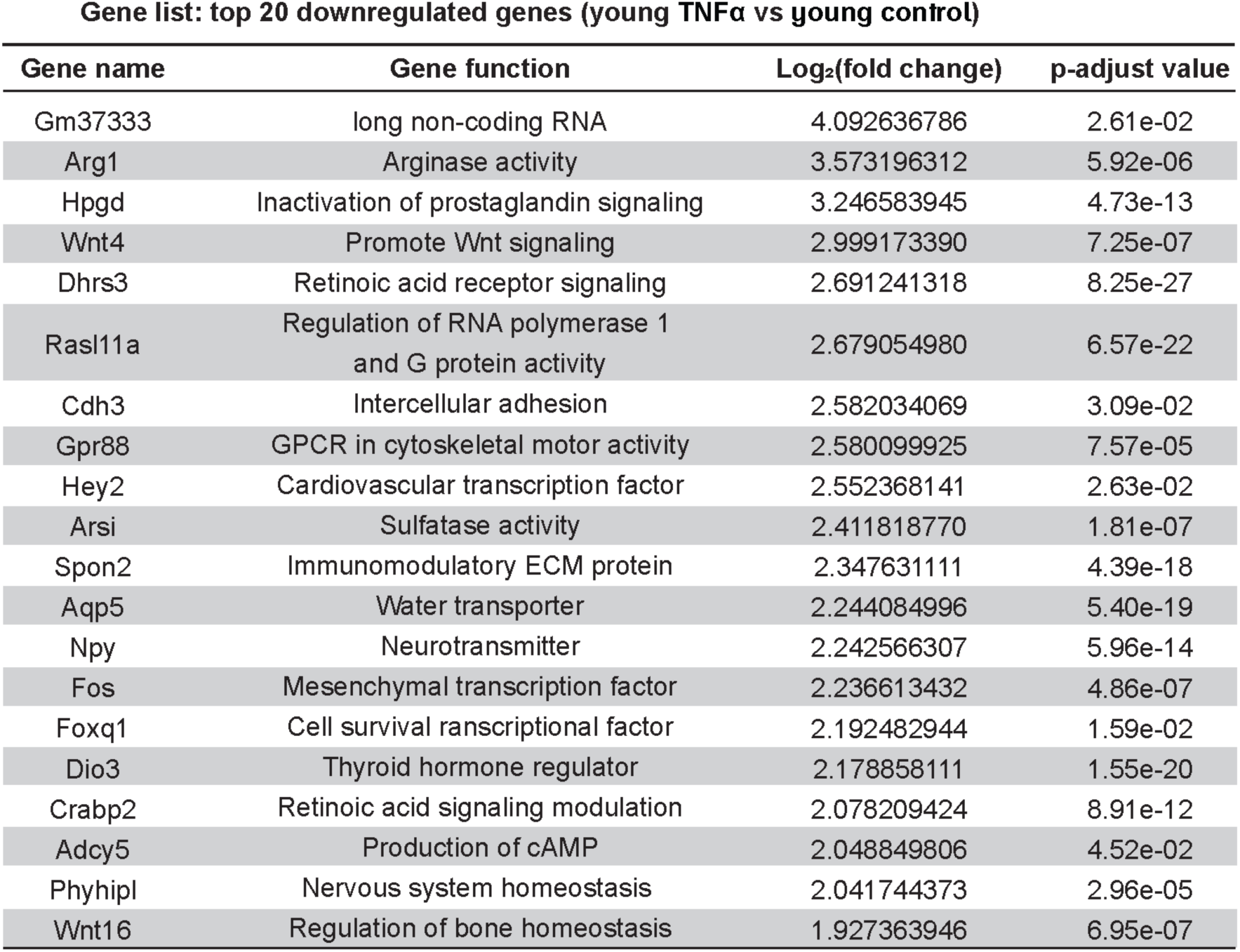
Top 20 differentially expressed genes with decreased expression in Young TNFα compared to Young Control.

**Table S21:**
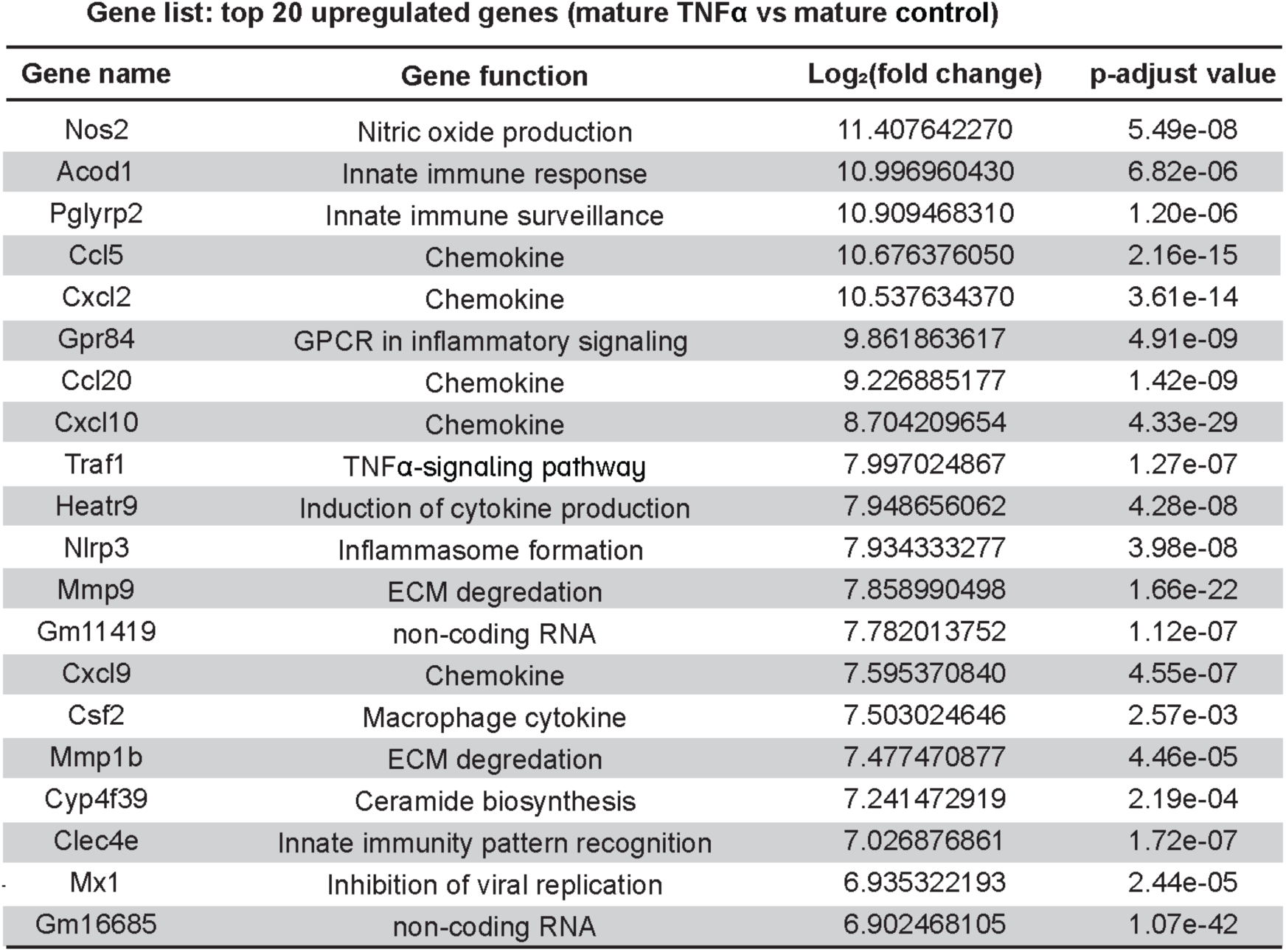
Top 20 differentially expressed genes with increased expression in Mature TNFα compared to Mature Control.

**Table S22:**
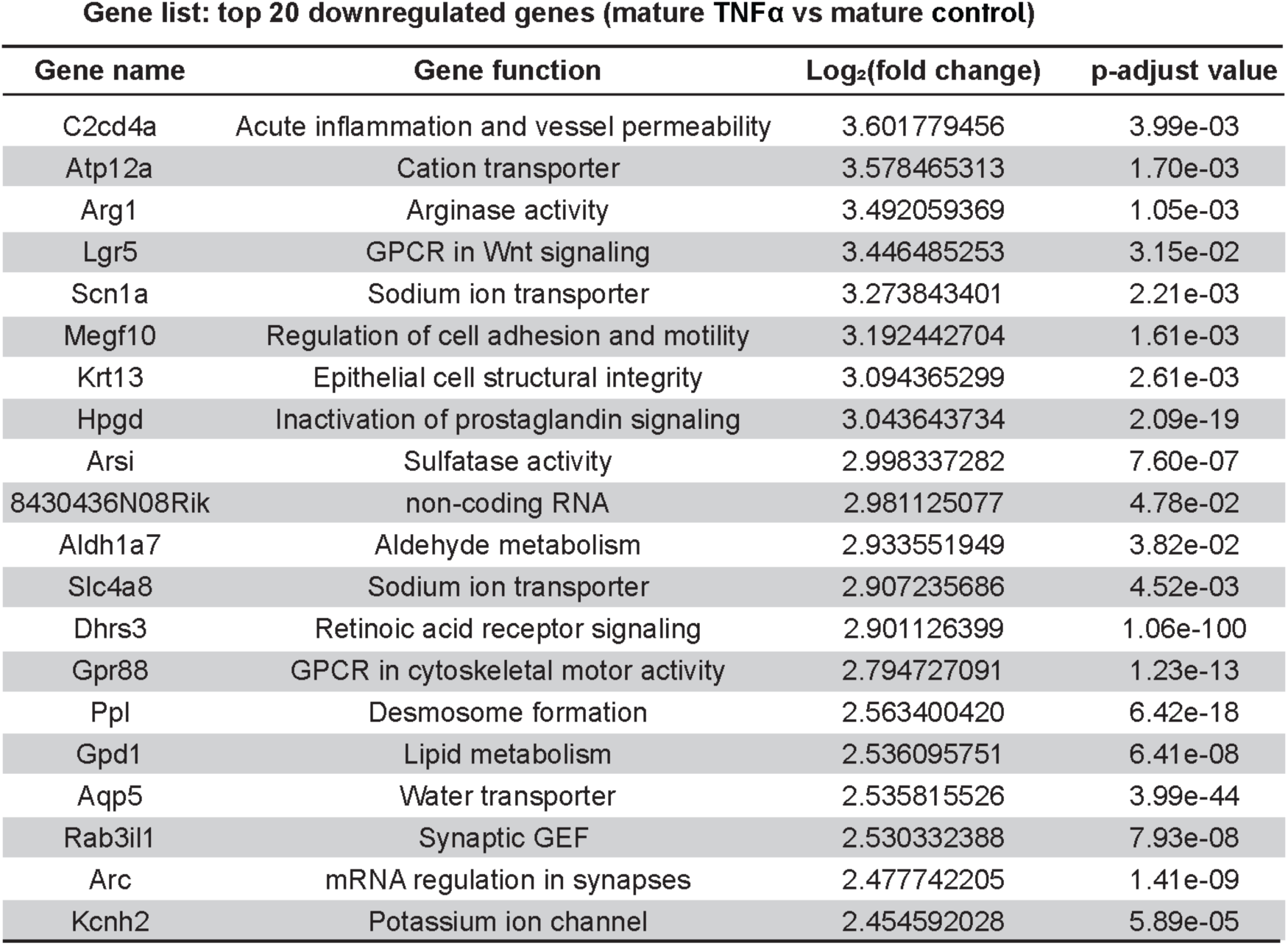
Top 20 differentially expressed genes with decreased expression in Mature TNFα compared to Mature Control.

**Table S23:**
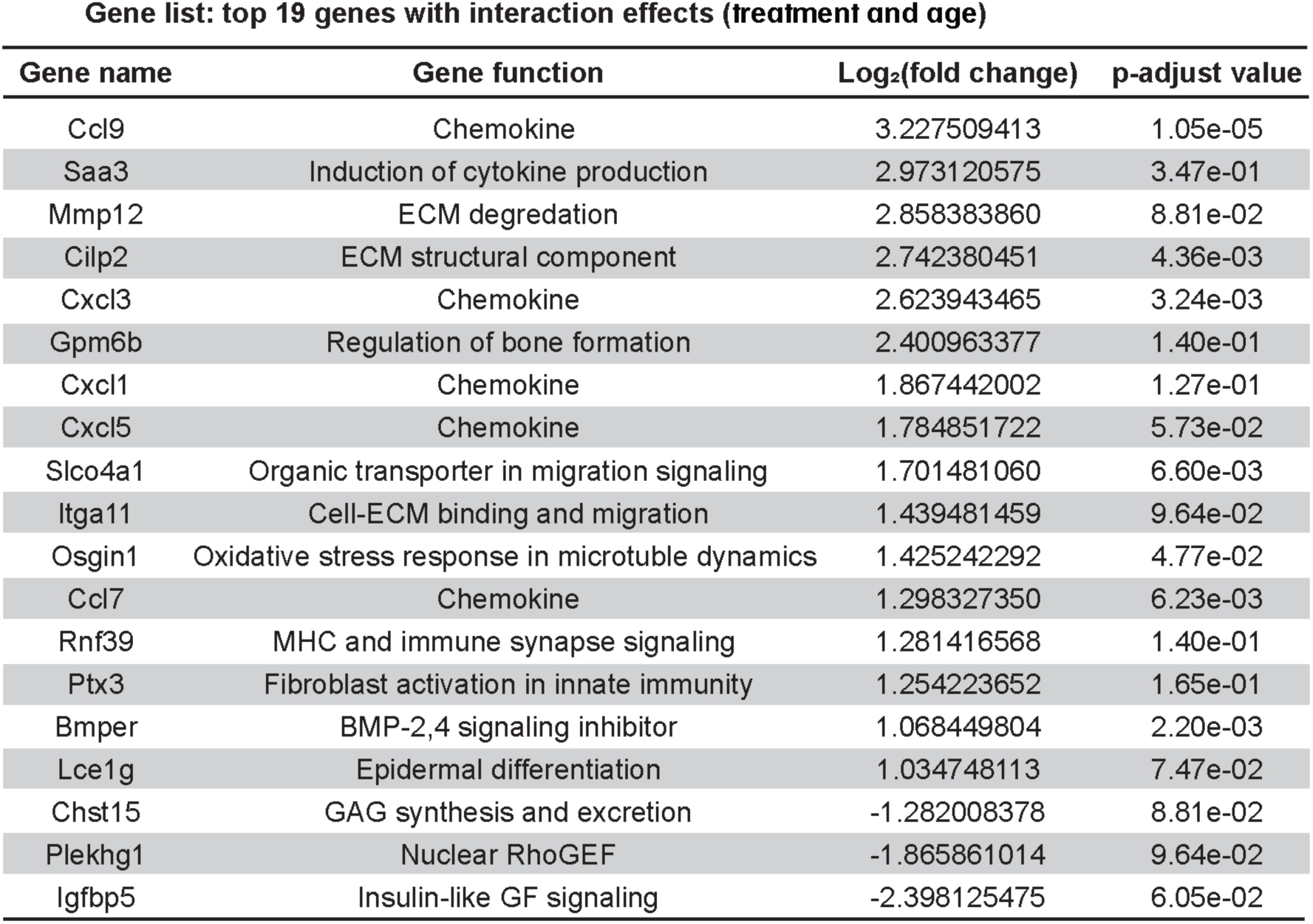
Top 19 genes identified with two-way interaction analysis (Age x Treatment) with p-adjust value < 0.4. Positive Log_2_(fold change) values indicate a more positive-valued response in mature cells, and negative Log_2_(fold change) values indicate a more positive-valued response in young cells.

**Table S24:**
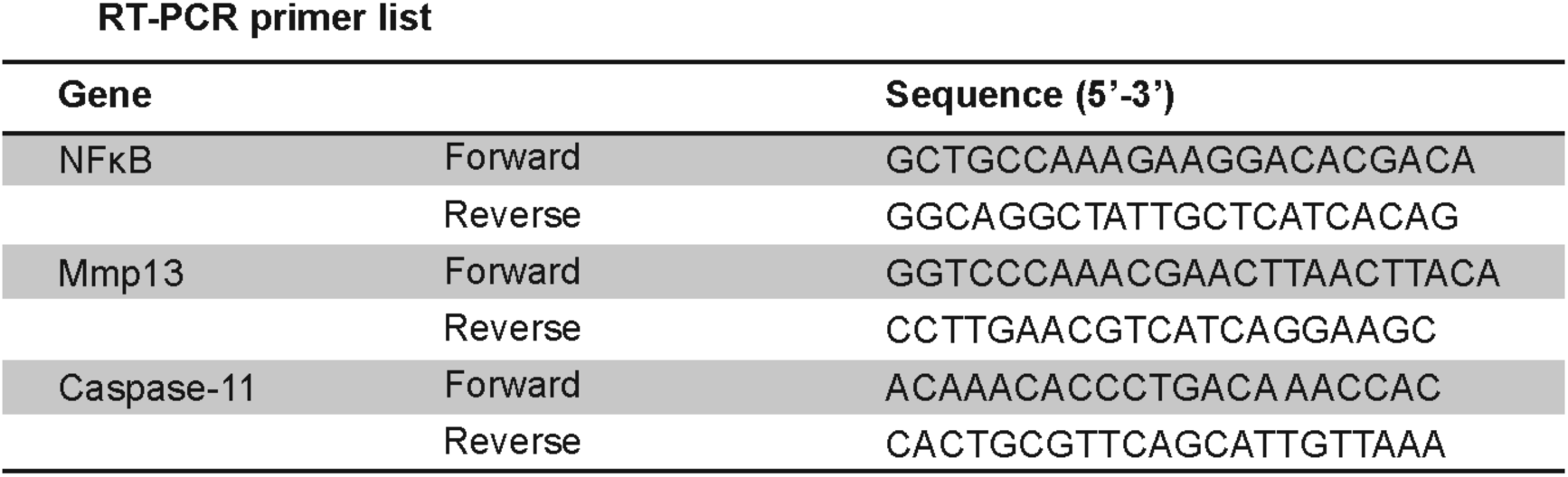
List of RT-PCR primers used in this study.

