## Supplemental information for "Age-Dependent Chromatin Remodeling Drives Inflammatory Dysregulation in Tendon Cells"

Tyler E. Blanch *et al.*

**This PDF file includes:**

Figs. S1 to S12
Tables S1 to S24

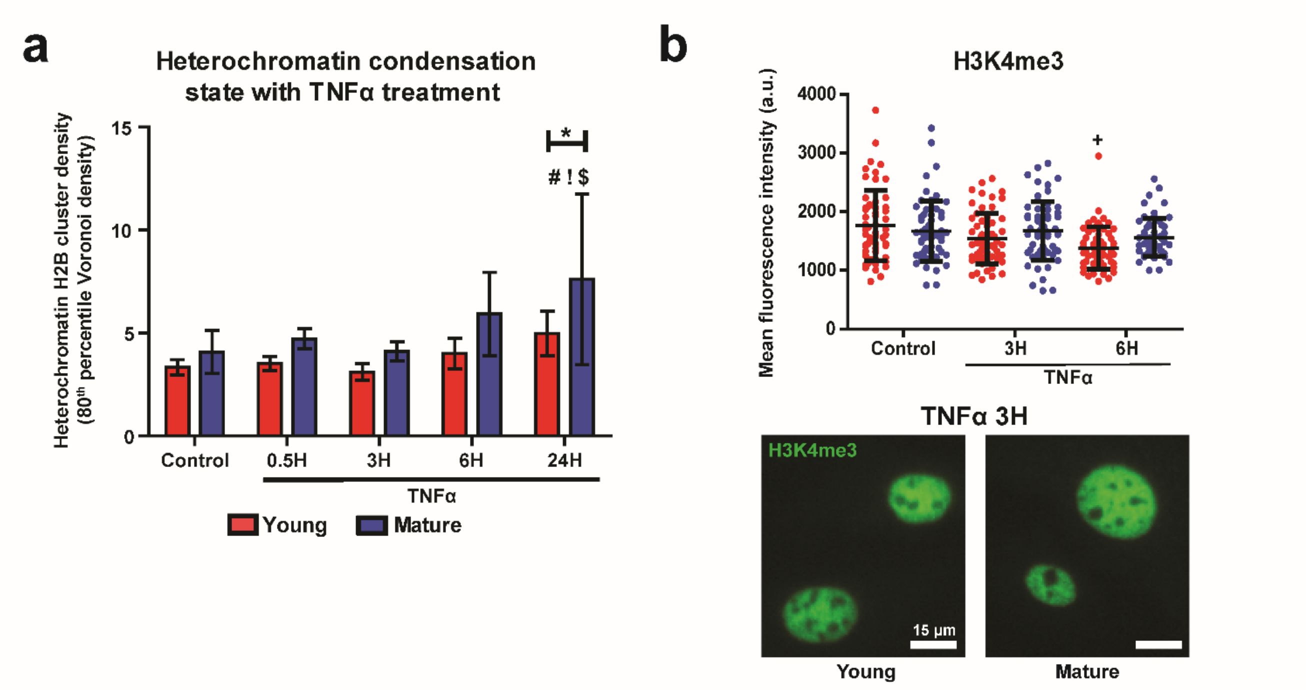


**Supplemental Figure S1: Epigenetic characterization of aged tenocytes under inflammatory treatment. (a)** Heterochromatin domain density of H2B-STORM localizations reported as the 80th percentile of the Voronoi density cumulative distribution function per cell, normalized to cell area. Shown for control and TNFα-treated conditions in young (red) and mature (blue) tenocytes. (n=16 cells/group). **(b)** Immunofluorescence (IF) mean intensity quantification (top) and representative images (bottom) for histone modification H3K4me3 in young (red) and mature (blue) tenocytes. (n=60 cells/group). (Mean ± SD, +: $p<0.05$ vs Young Control, #: $p<0.05$ vs mature Control, !: $p<0.05$ vs mature 0.5H, $: $p<0.05$ vs mature 3H, *: $p<0.05$).

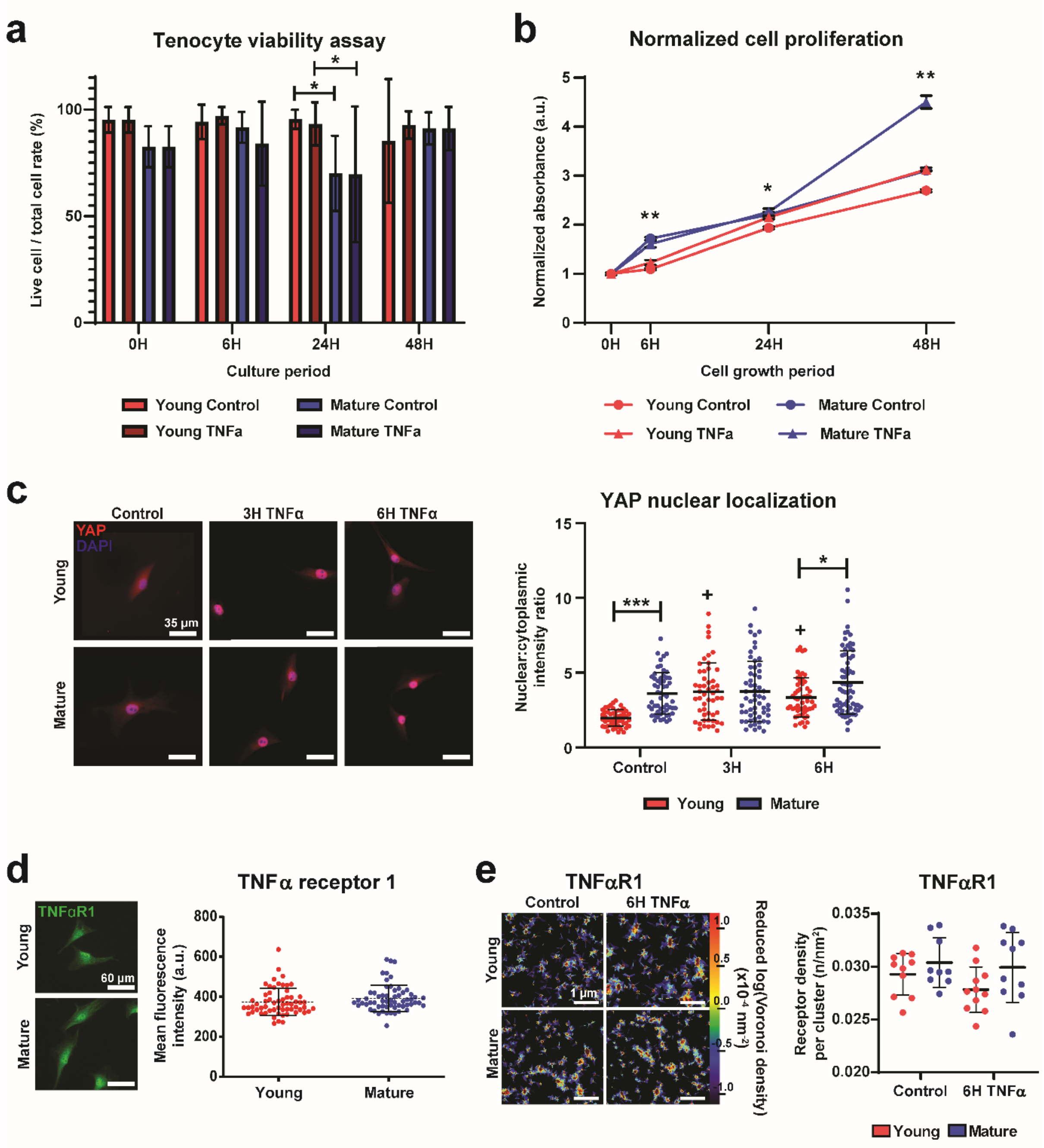


**Supplemental Figure S2: Tenocytes remain viable and mechanobiologically active under inflammatory conditions. (a)** Viability assay using Live/Dead staining of tenocytes under various control and inflammatory timepoints reported as the percentage of viable cells observed. Young groups colored in red and mature in blue, with TNFα-treated groups in darker colors. (n=100 cells/group). **(b)** Cell proliferation assay of young (red) and mature (blue) tenocytes at control (circle) and inflammatory (triangle) conditions. Values normalized to 0-hour (0H) results for each group. (n=2 technical replicates). **(c)** Representative IF images (left) and nuclear to cytoplasmic ratio (N:C) quantification (right) of YAP in young (red) and mature (blue) control and inflammatory-treated cells. (n=60 cells/group). **(d)** Representative IF images (left) and mean intensity quantification (right) for TNFα Receptor 1 in young and mature tenocytes at control conditions. (n=60 cells/group). **(e)** Representative STORM density heatmaps (left) and quantification of nanoscale receptor cluster density (right) for TNFα Receptor 1 in young (red) and mature (blue) tenocytes at control and inflammatory conditions. (n=11 cells/group, n=3 ROIs/cell). (Mean ± SD, +: $p<0.05$ vs Young Control, *: $p<0.05$, **: $p<0.01$, ***: $p<0.001$).

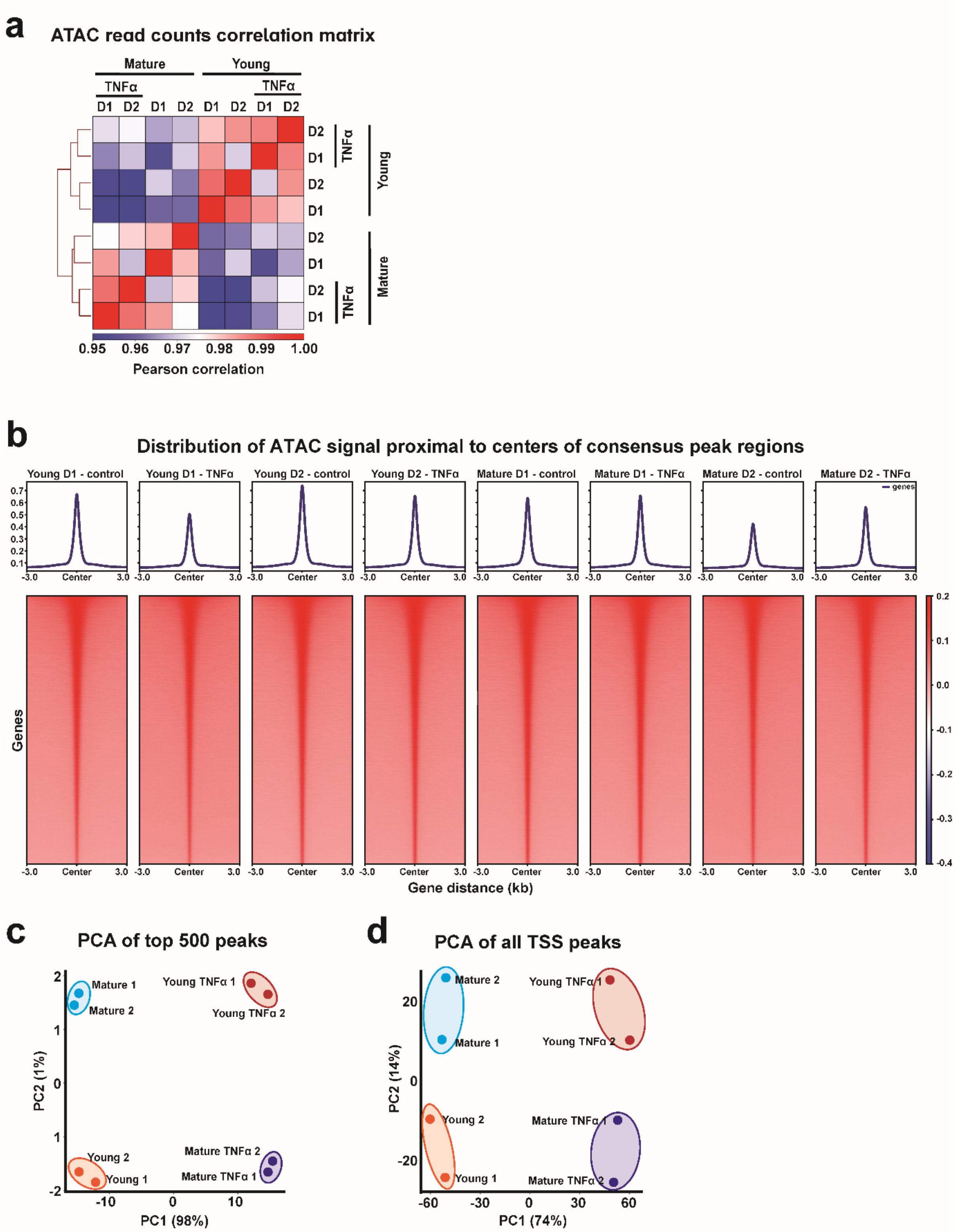


**Supplemental Figure S3: Quality control results of ATAC-seq reads. (a)** Sample correlation matrix using ATAC-seq read counts in young and mature tenocytes under control and TNFα-treated conditions. Color indicates Pearson correlation coefficient. **(b)** ATAC-seq read distribution profiles across each biological donor for control and TNFα-treated groups within 3 kb of consensus peak regions. **(c)** Principal component analysis (PCA) plot of the top 500 differential peaks comparing control and TNFα-treated young and mature tenocytes. **(d)** PCA plot of all transcription start site-associated (TSS-associated) peaks across control and TNFα-treated conditions in young and mature groups. (n=2 biological replicates per condition, TNFα treated for 6 hours, significance: $|Log_2(\text{fold change})|>1$ and p-adj<0.05).

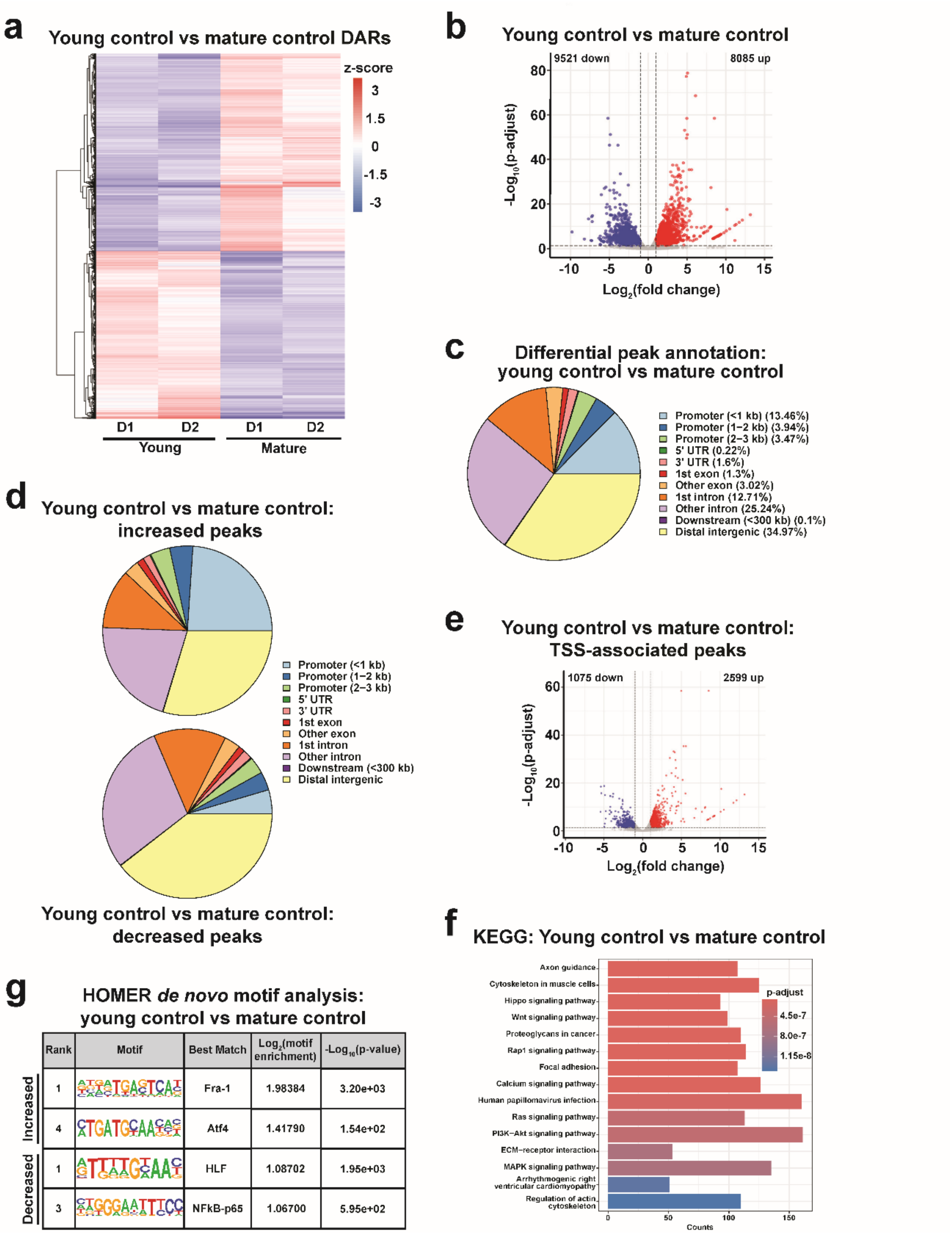


**Supplemental Figure S4: ATAC-seq analysis comparing chromatin accessibility between young and mature control tenocytes.** **(a)** Heatmap of differentially accessible regions (DARs) between Young Control and Mature Control tenocytes. Color represents z-score–scaled accessibility. **(b)** Volcano plot of all DARs between Young Control and Mature Control. Peaks with significantly increased (red) or decreased (blue) accessibility are indicated. **(c)** Gene annotation of DARs comparing Young Control and Mature Control. **(d)** Gene annotation results for increased (top) and decreased (bottom) peaks between Young Control and Mature Control group. **(e)** Volcano plot of all TSS-associated peaks between Young Control and Mature Control. Genes with significantly increased (red) or decreased (blue) accessibility are indicated. **(f)** Kyoto Encyclopedia of Genes and Genomes (KEGG) pathway enrichment analysis of TSS-associated DARs comparing Young Control and Mature Control. Color denotes statistical significance. **(g)** *De novo* motif enrichment analysis of increased (top) and decreased (bottom) TSS-associated peaks between Young Control and Mature Control tenocytes.(n=2 biological replicates per condition, significance (DARs): $|Log_2(\text{fold change})|>1$ and p-adj<0.05, significance (*de novo* motifs): $|Log_2(\text{motif enrichment})|>1$ and p-value<$1e^{-13}$).

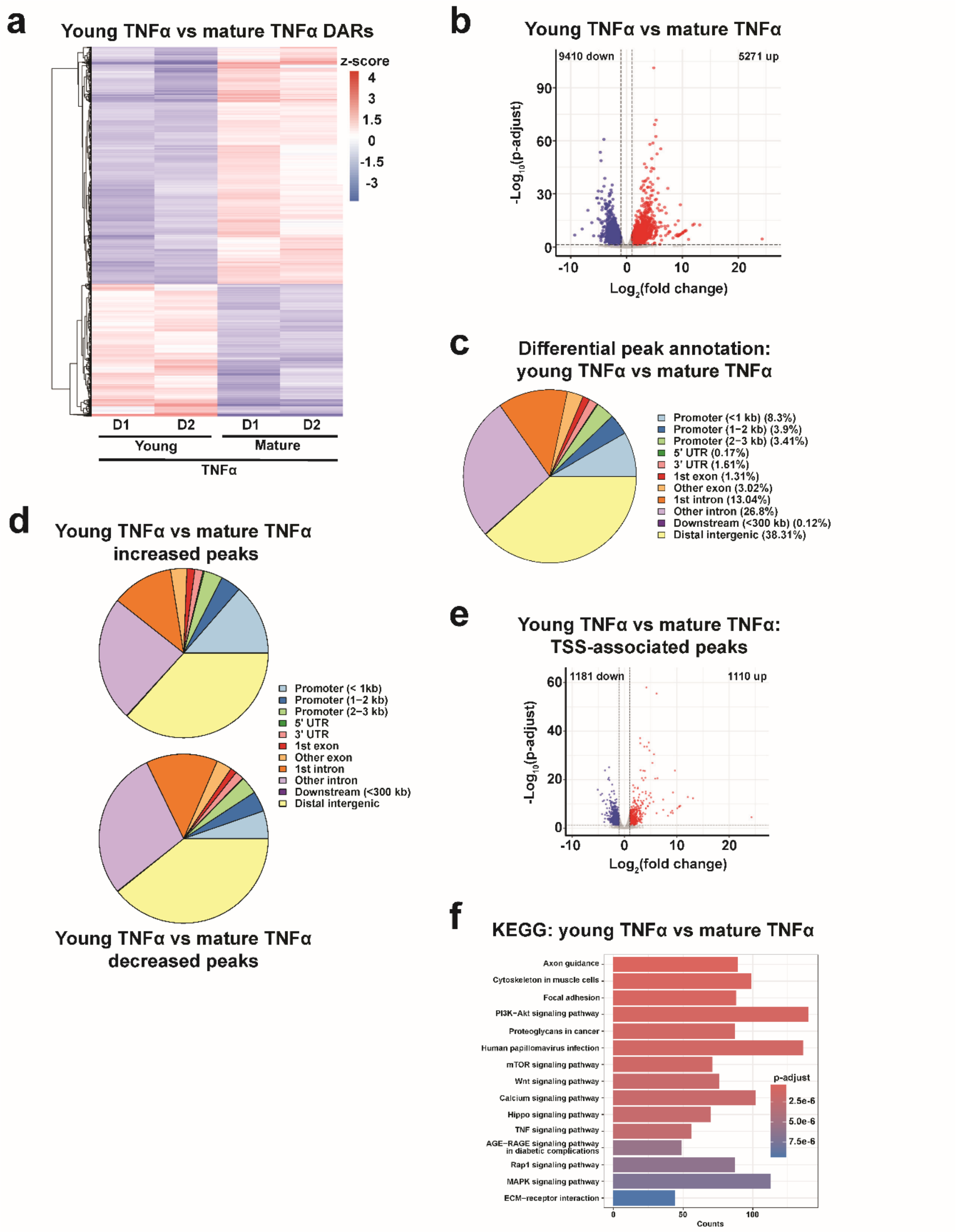


**Supplemental Figure S5: ATAC-seq analysis comparing chromatin accessibility between young and mature tenocytes under TNFα treatment. (a)** Heatmap of DARs between Young TNFα and Mature TNFα tenocytes. Color represents z-score–scaled accessibility. **(b)** Volcano plot of all DARs between Young TNFα and Mature TNFα. Peaks with significantly increased (red) or decreased (blue) accessibility are indicated. **(c)** Gene annotation of DARs comparing Young TNFα and Mature TNFα groups. **(d)** Gene annotation results for increased (top) and decreased (bottom) peaks between Young TNFα and Mature TNFα. **(e)** Volcano plot of TSS-associated peaks between Young TNFα and Mature TNFα. Genes with significantly increased (red) or decreased (blue) accessibility are indicated. **(f)** KEGG pathway enrichment analysis of TSS-associated DARs comparing Young TNFα and Mature TNFα tenocytes. Color denotes statistical significance. (n=2 biological replicates per condition, TNFα treated for 6 hours, significance: $|Log_2(\text{fold change})|>1$ and p-adj<0.05).

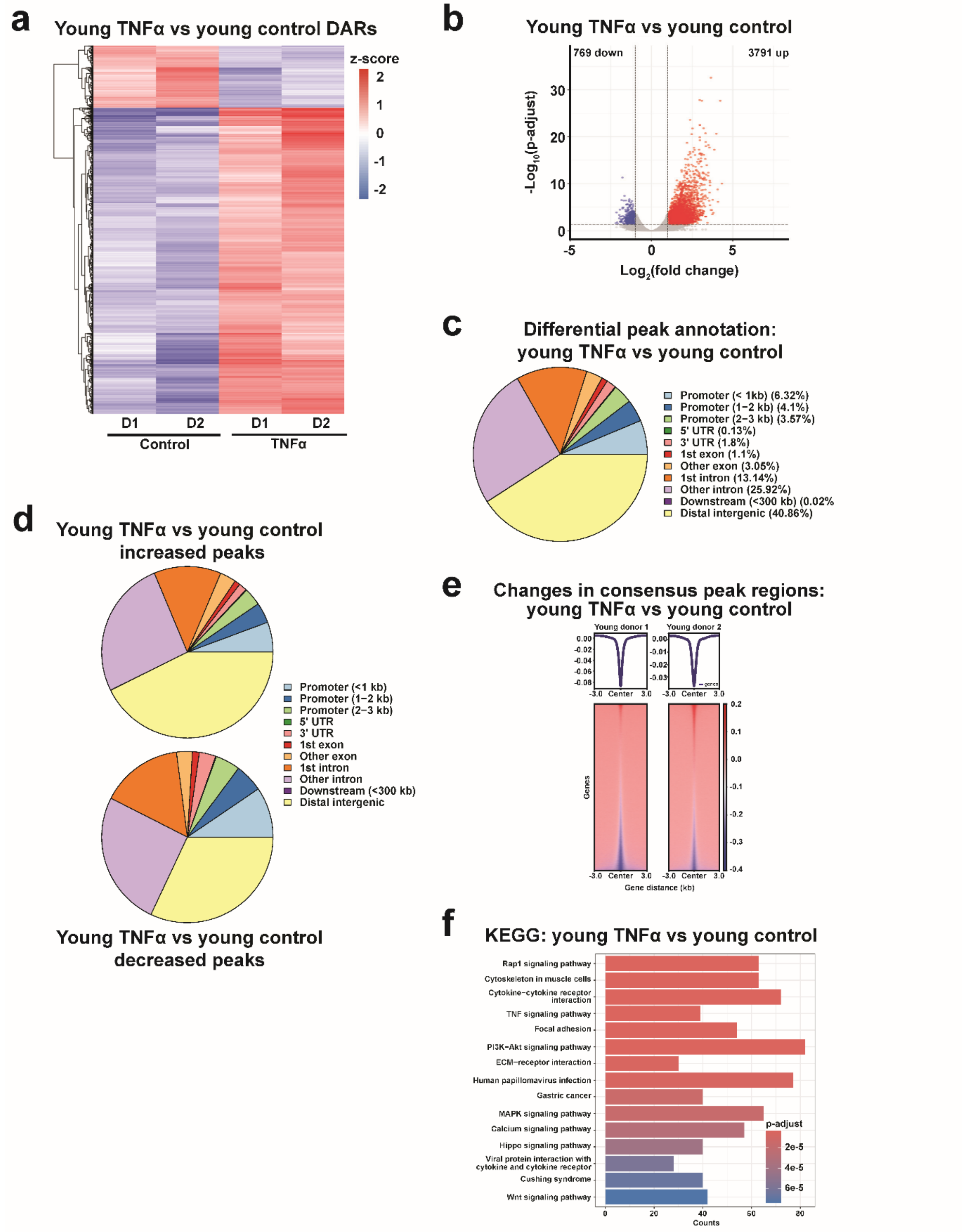


**Supplemental Figure S6: ATAC-seq analysis comparing chromatin accessibility between TNFα-treated and control young tenocytes. (a)** Heatmap of DARs between Young TNFα and Young Control tenocytes. Color represents z-score–scaled accessibility. **(b)** Volcano plot of all DARs between Young TNFα and Young Control. Peaks with significantly increased (red) or decreased (blue) accessibility are indicated. **(c)** Gene annotation of DARs comparing Young TNFα and Young Control groups. **(d)** Gene annotation results for increased (top) and decreased (bottom) peaks between Young TNFα and Young Control. **(e)** Differential peak enrichment plot of consensus peak regions between Young TNFα and Young Control. Color indicates accessibility changes. **(f)** KEGG pathway enrichment analysis of TSS-associated DARs comparing Young TNFα and Young Control tenocytes. Color denotes statistical significance. (n=2 biological replicates per condition, TNFα treated for 6 hours, significance: $|Log_2$(fold change)$|>1$ and p-adj<0.05).>1 and p-adj<0.05).

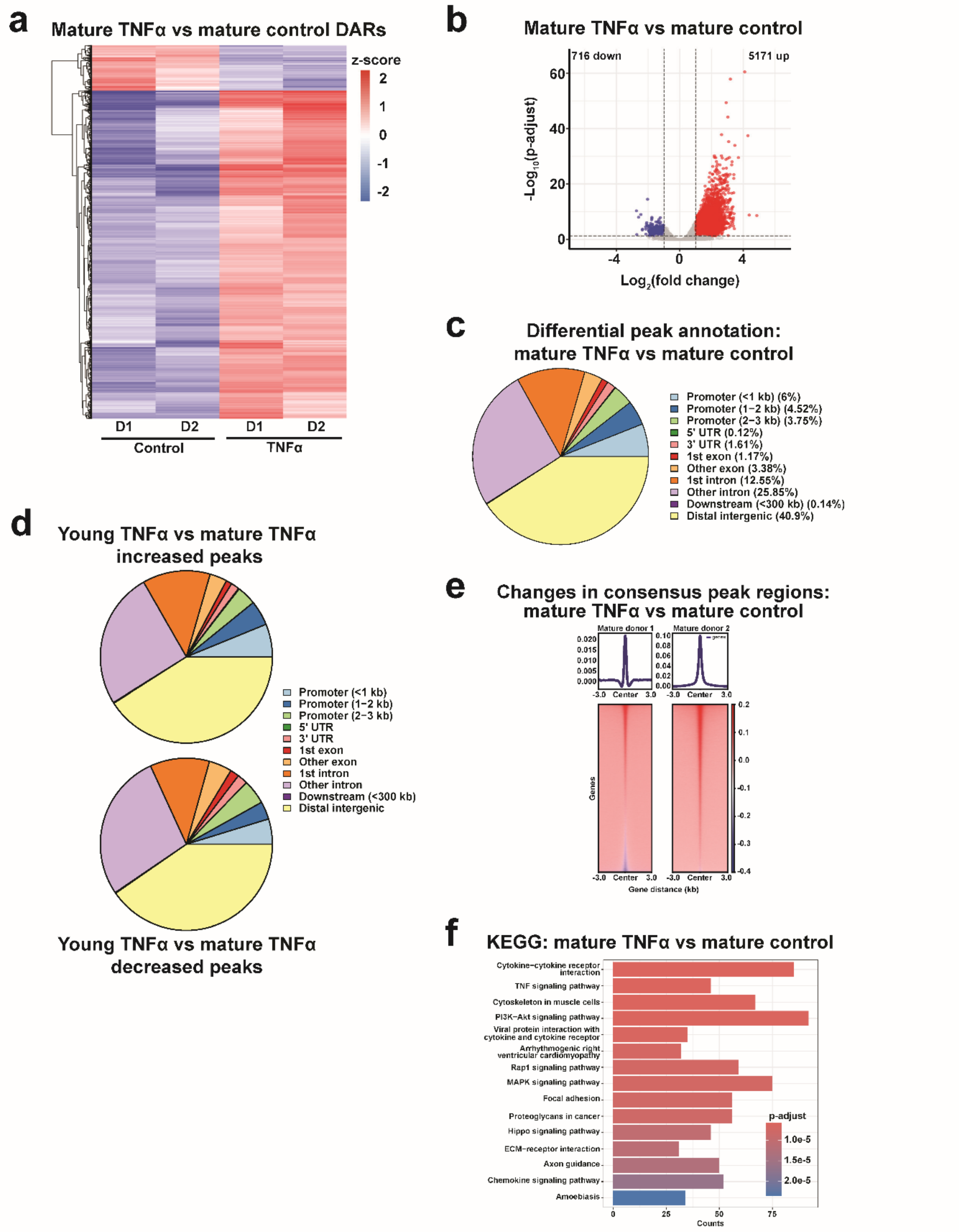


**Supplemental Figure S7: ATAC-seq analysis comparing chromatin accessibility between TNFα-treated and control mature tenocytes. (a)** Heatmap of DARs between Mature TNFα and Mature Control tenocytes. Color represents z-score–scaled accessibility. **(b)** Volcano plot of all DARs between Mature TNFα and Mature Control. Peaks with significantly increased (red) or decreased (blue) accessibility are indicated. **(c)** Gene annotation of DARs comparing Mature TNFα and Mature Control groups. **(d)** Gene annotation results for increased (top) and decreased (bottom) peaks between Mature TNFα and Mature Control. **(e)** Differential peak enrichment plot of consensus peak regions between Mature TNFα and Mature Control. Color indicates accessibility changes. **(f)** KEGG pathway enrichment analysis of TSS-associated DARs comparing Mature TNFα and Mature Control tenocytes. Color denotes statistical significance. (n=2 biological replicates per condition, TNFα treated for 6 hours, significance: $|Log_2(\text{fold change})|>1$ and p-adj<0.05).

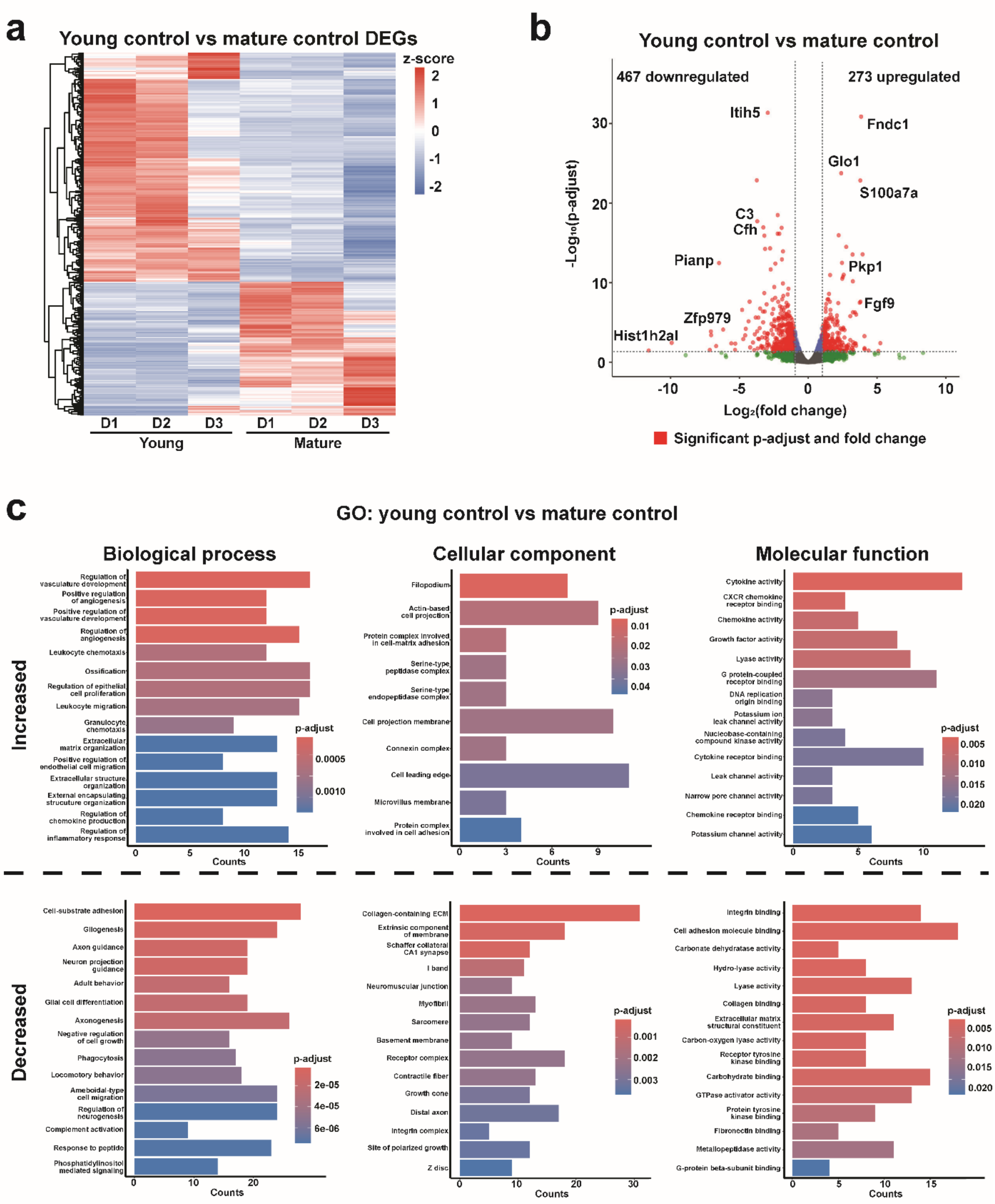


**Supplemental Figure S8: RNA-seq analysis comparing gene expression between young and mature control tenocytes. (a)** Heatmap of differentially expressed genes (DEGs) comparing Young Control and Mature Control tenocytes. Color represents z-score–scaled expression levels. **(b)** Volcano plot of DEGs between Young Control and Mature Control groups, with significantly altered genes indicated in red. **(c)** Gene Ontology (GO) enrichment analysis of DEGs comparing Young Control and Mature Control tenocytes, categorized by Biological Process (left), Cellular Component (middle), and Molecular Function (right). Terms with increased (top) or decreased (bottom) prevalence in young control cells are colored by statistical significance. (n=3 biological replicates, significance: $|Log_2(\text{fold change})| > 1$ and $p\text{-adj} < 0.05$).

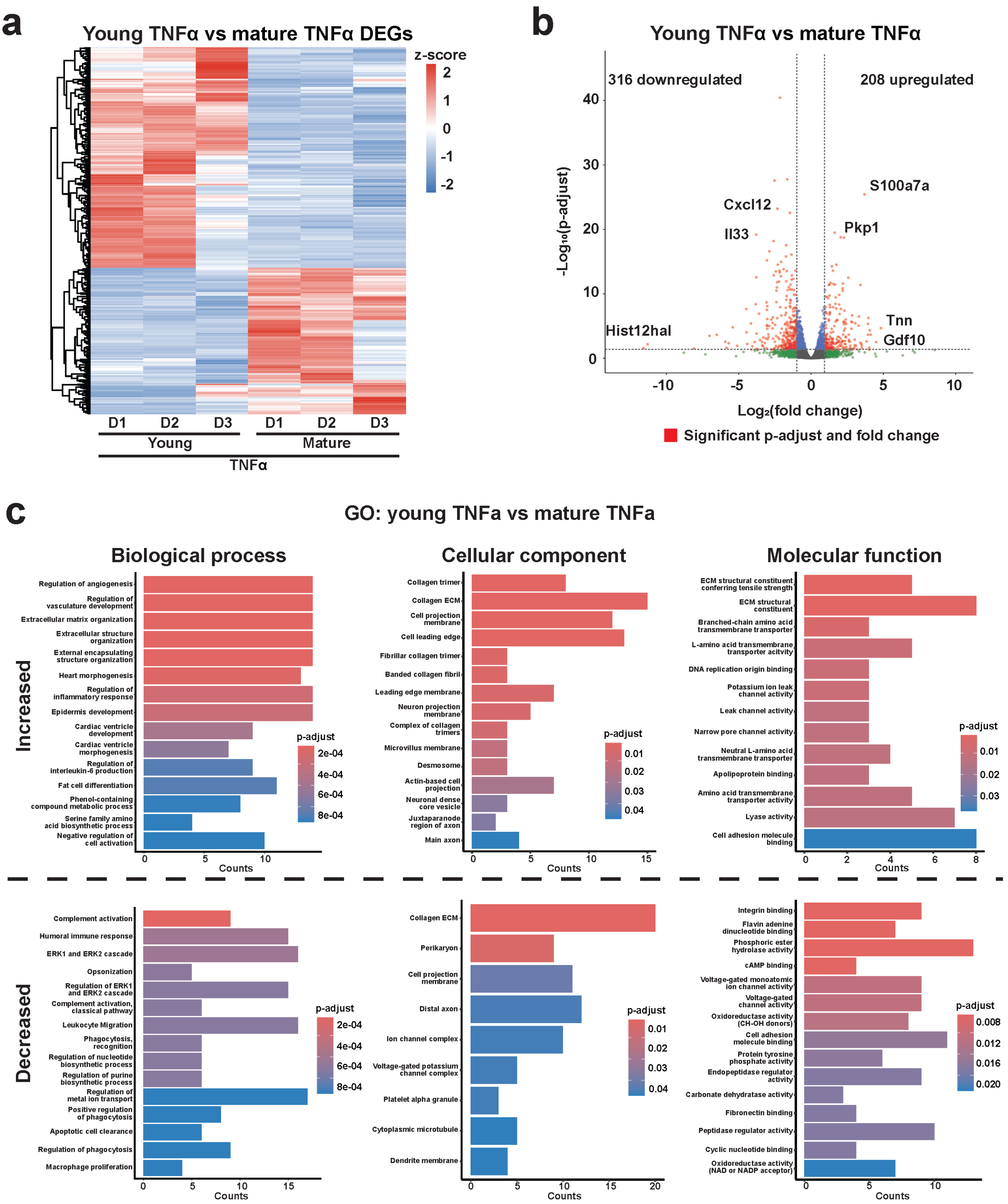


**Supplemental Figure S9: RNA-seq analysis comparing gene expression between young and mature tenocytes under TNFα stimulation. (a)** Heatmap of DEGs comparing Young TNFα and Mature TNFα tenocytes. Color represents z-score–scaled expression levels. **(b)** Volcano plot of DEGs between Young TNFα and Mature TNFα groups, with significantly altered genes highlighted in red. **(c)** GO enrichment analysis of DEGs comparing Young TNFα and Mature TNFα tenocytes, categorized by Biological Process (left), Cellular Component (middle), and Molecular Function (right). Terms with increased (top) or decreased (bottom) prevalence in young TNFα-treated cells are colored according to statistical significance. (n=3 biological replicates, TNFα treated for 6 hours, significance: $|Log_2(\text{fold change})| > 1$ and p-adj<0.05).

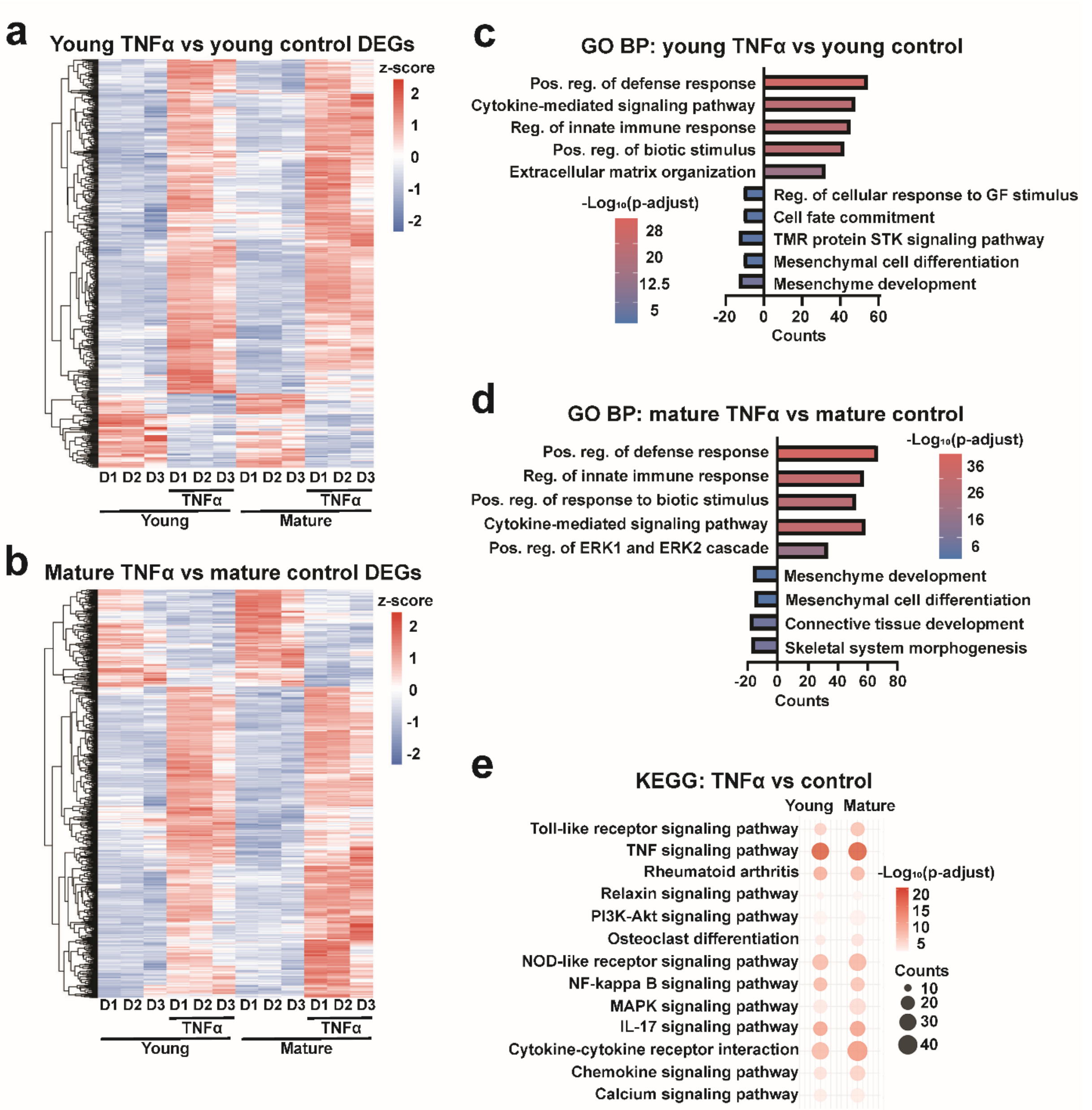


**Supplemental Figure S10: RNA-seq analysis comparing TNFα-treated and control tenocytes across age groups. (a, b)** Heatmaps of DEGs comparing Young TNFα versus Young Control **(a)** and Mature TNFα versus Mature Control **(b)** groups. Color represents z-score–scaled expression levels. **(c, d)** GO enrichment analysis of DEGs within the Biological Process (BP) category for Young TNFα versus Young Control **(c)** and Mature TNFα versus Mature Control **(d)** comparisons. Color denotes significance, and the direction of counts indicates gene upregulation (positive) or downregulation (negative). **(e)** KEGG pathway enrichment analysis of DEGs comparing TNFα-treated to control tenocytes in young (left) and mature (right) groups. Color indicates significance, and circle size represents gene count. (n=3 biological replicates, TNFα treated for 6 hours, significance: $|Log_2$(fold change)$|>1$ and p-adj<0.05).

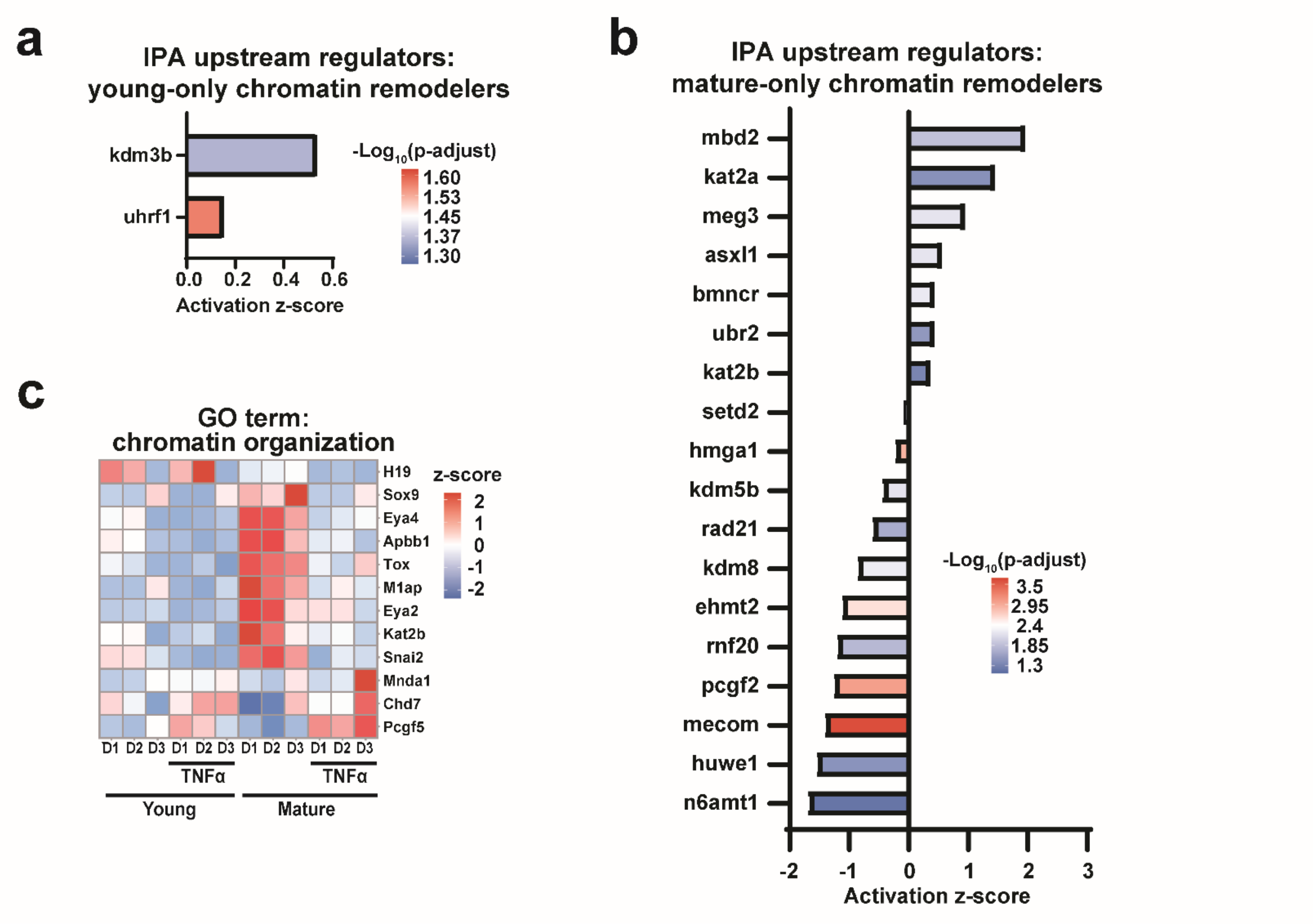


**Supplemental Figure S11: Chromatin state modifiers associated with inflammatory stimulation.** **(a, b)** Predicted upstream regulators identified by Ingenuity Pathway Analysis (IPA) from age-exclusive TNFα-induced DEGs overlapping with the GO term "Chromatin Remodelers." Results are shown for young **(a)** and mature **(b)** tenocytes, with color representing statistical significance. **(c)** Heatmap of age-exclusive DEGs within the GO term "Chromatin Organization" for young and mature tenocytes following TNFα stimulation. Color represents z-score–scaled expression levels. (n=3 biological replicates, TNFα treated for 6 hours, significance: $|Log_2(\text{fold change})|>1$ and p-adj<0.05).

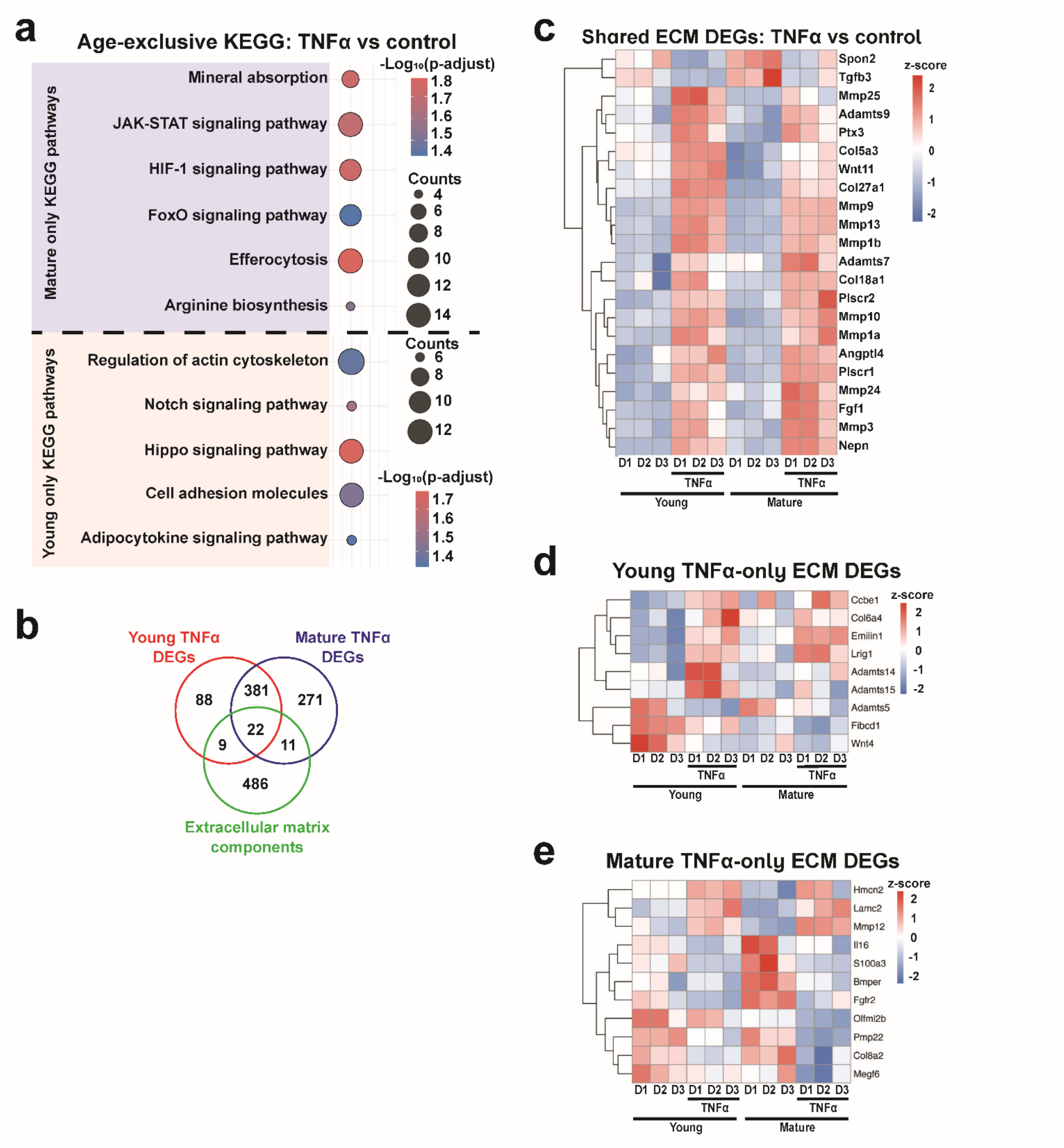


**Supplemental Figure S12: Altered extracellular matrix remodeling following TNFα stimulation in young and mature tenocytes. (a)** KEGG pathway enrichment analysis of age-exclusive DEGs comparing TNFα-treated and control tenocytes in young and mature groups. Color denotes statistical significance, and circle size represents gene count. Age-exclusive results for young and mature tenocytes are highlighted in red and blue, respectively. **(b)** Diagram showing TNFα-induced DEGs in young (red) and mature (blue) tenocytes overlapping with the GO term "Extracellular Matrix Components and Regulators" (green). **(c-e)** Heatmaps of ECM-related DEGs significantly regulated in both age groups (shared) **(c)**, or exclusive to young **(d)** or mature **(e)** tenocytes following TNFα treatment. Color represents z-score–scaled expression. (n=3 biological replicates, TNFα treated for 6 hours, significance: $|Log_2(\text{fold change})|>1$ and p-adj<0.05).

**Table S1:**

**Annotated peak list: top 20 promoter regions with increased accessibility (young control vs mature control)**

| Peak | Distance to TSS (bp) | Gene | $Log_2$(fold change) | p-adjust value |
|---|---|---|---|---|
| chr4:88,675,641-88,676,220 | 0 | Gm13286 | 13.16902442 | 6.83e-16 |
| chr14:121,042,545-121,043,237 | 459 | Gm41253 | 12.12768344 | 1.72e-13 |
| chr13:619,517,34-61,952,578 | 0 | Gm7240 | 11.72774042 | 1.80e-12 |
| chr4:88,724,023-88,724,415 | 255 | Gm13288 | 10.50005949 | 1.54e-09 |
| chr4:88,678,659-88,679,100 | 303 | Gm13283 | 10.15190476 | 2.99e-18 |
| chr11:32,249,858-32,250,373 | 0 | Hbq1a | 10.06788454 | 1.04e-08 |
| chr4:55,421,241-55,421,684 | -2,399 | Gm12505 | 9.284332468 | 4.67e-07 |
| chr4:88,700,488-88,700,970 | 120 | Ifnz | 9.225393584 | 7.61e-07 |
| chr16:36,125,036-36,125,827 | 0 | Cstdc3 | 9.219252219 | 6.21e-07 |
| chr6:3,384,136-3,384,437 | 569 | Samd9l | 9.071382725 | 1.28e-06 |
| chr16:36,078,771-36,079,217 | -792 | Gm24569 | 8.721877113 | 5.97e-06 |
| chr17:34,199,826-34,200,680 | -2,015 | H2-K2 | 8.536390503 | 3.33e-59 |
| chr4:88,712,506-88,712,857 | 0 | Gm13275 | 8.497391668 | 1.59e-05 |
| chr2:58,056,839-58,057,148 | -2,601 | Gm13546 | 8.464064695 | 1.76e-05 |
| chr14:53,276,925-53,277,242 | -2 | Gm30214 | 8.358780664 | 2.88e-05 |
| chr17:36,478,380-36,479,504 | 0 | H2-T26 | 8.095599145 | 1.23e-10 |
| chr4:88,706,626-88,707,003 | 0 | Gm13277 | 8.038489733 | 2.41e-10 |
| chr17:35,612,843-35,613,639 | 0 | H2-Q5 | 7.659514343 | 1.05e-08 |
| chr4:42,734,593-42,735,141 | -404 | 4930578G10Rik | 6.847368668 | 4.97e-10 |
| chr17:35,219,259-35,219,564 | 0 | D17H6S56E-5 | 6.661874988 | 3.79e-06 |

**Table S1: Top 20 transcription start site-associated (TSS-associated) differentially accessible regions (DARs) with increased accessibility in Young Control compared to Mature Control tenocytes.**

**Table S2:**

**Annotated peak list: top 20 promoter regions with decreased accessibility (young control vs mature control)**

| Peak | Distance to TSS (bp) | Gene | Log$_2$(fold change) | p-adjust value |
|---|---|---|---|---|
| chr8:15,046,895-15,047,128 | 759 | Arhgef10 | 5.457658689 | 1.09e-08 |
| chr2:77,769,683-77,770,655 | -2,612 | Cwc22 | 5.406802775 | 3.31e-19 |
| chr2:77,764,086-77,764,458 | 2,555 | Cwc22 | 5.345865384 | 2.05e-16 |
| chr2:77,768,380-77,769,040 | -1,309 | Cwc22 | 4.979120475 | 1.76e-19 |
| chr3:100,102,102-100,102,735 | 206 | Gdap2 | 4.960554757 | 1.86e-02 |
| chr2:77,711,914-77,713,012 | -103 | Cwc22 | 4.904789344 | 9.81e-04 |
| chr10:100,376,857-100,377,196 | 541 | Cep290 | 4.871764270 | 1.11e-15 |
| chr7:43,617,004-43,617,401 | 829 | Klk1b1 | 4.788360542 | 7.20e-06 |
| chr2:77,740,651-77,741,016 | -2,140 | Cwc22 | 4.767913811 | 5.74e-09 |
| chr19:53,445,215-53,445,544 | -2,681 | Mirt1 | 4.288250167 | 1.08e-07 |
| chr1:127,826,382-127,826,799 | -1,851 | Rab3gap1 | 4.258965574 | 2.94e-09 |
| chrX:153,877,445-153,878,004 | -783 | Magea2 | 4.199675944 | 2.93e-15 |
| chr2:77,738,208-77,738,856 | 0 | Cwc22 | 4.198893332 | 3.11e-15 |
| chr2:77,739,719-77,739,964 | -1,208 | Cwc22 | 4.194288557 | 9.65e-11 |
| chr13:59,973,219-59,973,435 | -2,258 | Tut7 | 4.089739678 | 1.30e-08 |
| chr12:25,445,770-25,446,430 | 1,016 | Gm36723 | 4.084287397 | 2.06e-08 |
| chr7:37,888,586-37,888,829 | 2,890 | 1600014C10Rik | 4.072568911 | 9.23e-05 |
| chr17:35,044,144-35,045,201 | 0 | Stk19 | 3.886137824 | 6.02e-15 |
| chr2:77,755,952-77,756,175 | -1,080 | Cwc22 | 3.851768332 | 9.50e-06 |
| chr13:99,579,005-99,579,666 | 786 | Map1b | 3.822081944 | 3.46e-16 |

**Table S2: Top 20 TSS-associated DARs with decreased accessibility in Young Control compared to Mature Control tenocytes.**

**Table S3:**

**Enriched motifs list: top 20 *de novo* motifs with increased accessibility (young control vs mature control)**

| Motif consensus | Best match | Match score | $Log_2$(motif enrichment) | $-Log_{10}$(p-value) |
|---|---|---|---|---|
| NNDVTGASTCAY | Fra-1 | 0.99 | 1.98384 | 3.20e+03 |
| GCCACACCCWNN | Eklf | 0.97 | 0.78966 | 2.14e+02 |
| BWRACCACAS | Runx1 | 0.95 | 0.54647 | 1.73e+02 |
| CTGATGCAACAB | Atf4 | 0.97 | 1.41790 | 1.54e+02 |
| GCATGACTTGGC | Nfe2l2 | 0.70 | 0.58031 | 1.53e+02 |
| TGGCAKCCTGCC | Tlx | 0.90 | 0.80995 | 1.15e+02 |
| AACATTCC | Tead2 | 0.99 | 0.33939 | 9.73e+01 |
| GGAAAATATGAC | Nfat:Ap1 | 0.84 | 0.86484 | 8.77e+01 |
| TTGCCAAG | Nfic | 0.97 | 0.35255 | 8.26e+01 |
| TGGHRASCHR | Nr2c2 | 0.66 | 0.61237 | 7.09e+01 |
| AGCGGCGGCGGC | Znf93 | 0.79 | 0.69302 | 6.87e+01 |
| CCCGGCTGCCGG | Zic1 | 0.62 | 0.56586 | 6.85e+01 |
| ATGCCGCCCG | Znf961 | 0.74 | 0.66456 | 6.85e+01 |
| RATGAGSTCA | Atf2 | 0.89 | 0.46067 | 5.76e+01 |
| CGCGCCGCGC | Znf161 | 0.77 | 0.51715 | 5.57e+01 |
| AGCCCAGC | Znf549 | 0.81 | 0.38002 | 5.51e+01 |
| AAATGCCA | Nf4a2 | 0.75 | 0.36785 | 4.92e+01 |
| GCGCCCGGAG | Znf454 | 0.65 | 0.56567 | 4.86e+01 |
| CATAAATCMT | Pbx2 | 0.94 | 0.72896 | 4.62e+01 |
| CTGTKGCCAG | Nf1 | 0.62 | 0.27681 | 4.56e+01 |

**Table S3: Top 20 differentially accessible transcription factor motifs with increased accessibility in Young Control compared to Mature Control tenocytes.**

**Table S4:**

**Enriched motifs list: top 20 *de novo* motifs with decreased accessibility (young control vs mature control)**

| Motif consensus | Best match | Match score | $Log_2$(motif enrichment) | $-Log_{10}$(p-value) |
|---|---|---|---|---|
| RTTWTGHAAY | Hlf | 0.89 | 1.08702 | 1.95e+03 |
| DATGASTCAT | Batf | 0.99 | 0.90160 | 9.08e+02 |
| HWGGGAATTTCC | Nfkb-p65 | 0.95 | 1.06700 | 5.95e+02 |
| GGAATGTKNT | Tead2 | 0.95 | 0.56353 | 4.05e+02 |
| YKGGAAAA | Nfatc1 | 0.88 | 0.38633 | 2.40e+02 |
| ATGANVTCATYY | Creb5 | 0.95 | 0.65848 | 1.94e+02 |
| TCTTGGCA | Nfic | 0.96 | 0.38286 | 1.53e+02 |
| WCTGTGGTTT | Runx1 | 0.94 | 0.56828 | 1.33e+02 |
| GSAADKCTGW | Maff | 0.63 | 0.37206 | 1.16e+02 |
| GCTCATAAAA | Hoxc13 | 0.87 | 0.98254 | 1.09e+02 |
| TGSTGAMT | Mafa | 0.80 | 0.30554 | 9.24e+01 |
| GTTGCGTAAGAC | Cebpg | 0.72 | 1.71400 | 9.17e+01 |
| ACACTGCGTG | Zbtb3 | 0.66 | 0.36603 | 8.00e+01 |
| CAGCTGGC | Tcf12 | 0.86 | 0.34555 | 7.00e+01 |
| CTCCCTGGAGAA | Bcl6 | 0.68 | 0.56116 | 6.95e+01 |
| CAGCTCTG | Myf5 | 0.72 | 0.32193 | 6.75e+01 |
| ATTACTCACACT | Mga | 0.56 | 1.04924 | 6.39e+01 |
| ATTCCTCATAAC | Stat6 | 0.68 | 0.78588 | 5.83e+01 |
| TTDCGKARCT | Gmeb2 | 0.69 | 0.74919 | 5.62e+01 |
| CTATTTTTAG | Mef2a | 0.99 | 0.90182 | 4.71e+01 |

**Table S4: Top 20 differentially accessible transcription factor motifs with decreased accessibility in Young Control compared to Mature Control tenocytes.**

**Table S5:**

**Annotated peak list: top 20 promoter regions with increased accessibility (young TNFα vs mature TNFα)**

| Peak | Distance to TSS (bp) | Gene | Log$_2$(fold change) | p-adjust value |
|---|---|---|---|---|
| chr7:103,963,777-103,964,825 | 0 | Gm15133 | 24.27654584 | 3.09e-05 |
| chr4:88,675,641-88,676,220 | 0 | Gm13286 | 13.09775853 | 4.37e-13 |
| chr14:121,042,545-121,043,237 | 459 | Gm41253 | 12.10664269 | 1.45e-13 |
| chr4:88,724,023-88,724,415 | 255 | Gm13288 | 10.65883426 | 5.99e-10 |
| chr16:36,125,036-36,125,827 | 0 | Cstdc3 | 10.48060744 | 8.12e-10 |
| chr4:88,706,626-88,707,003 | 0 | Gm13277 | 10.42526845 | 1.56e-09 |
| chr16:36,078,771-36,079,217 | -792 | Gm24569 | 10.09529233 | 6.08e-09 |
| chr4:88,678,659-88,679,100 | 303 | Gm13283 | 9.641980175 | 1.65e-24 |
| chr13:61,951,734-61,952,578 | 0 | Gm7240 | 9.260200942 | 1.85e-15 |
| chr4:88,700,488-88,700,970 | 120 | Ifnz | 9.233053269 | 5.58e-07 |
| chr17:35,612,843-35,613,639 | 0 | H2-Q5 | 9.107899460 | 6.58e-08 |
| chr17:35,219,259-35,219,564 | 0 | D17H6S56E-5 | 8.848812648 | 3.19e-07 |
| chr17:34,199,826-34,200,680 | -2,015 | H2-K2 | 8.315396018 | 7.97e-06 |
| chr4:55,421,241-55,421,684 | -2,399 | Gm12505 | 7.421861673 | 2.92e-08 |
| chr11:32,249,858-32,250,373 | 0 | Hbq1a | 7.401295580 | 1.55e-12 |
| chr17:36,478,380-36,479,504 | 0 | H2-T26 | 6.303999229 | 2.55e-21 |
| chr4:133,375,329-133,375,731 | 1,734 | Zdhhc18 | 6.127447932 | 6.17e-07 |
| chr14:51,193,303-51,194,598 | 0 | Pnp2 | 6.126136435 | 3.70e-56 |
| chr17:24,886,048-24,886,344 | -2,317 | Zfp598 | 5.891395757 | 3.53e-21 |
| chr18:43,610,633-43,611,219 | 0 | Eif3j2 | 5.638983040 | 3.34e-31 |

**Table S5: Top 20 TSS-associated DARs with increased accessibility in Young TNFα compared to Mature TNFα tenocytes.**

**Table S6:**

**Annotated peak list: top 20 promoter regions with decreased accessibility (young TNFα vs mature TNFα)**

| Peak | Distance to TSS (bp) | Gene | Log$_2$(fold change) | p-adjust value |
|---|---|---|---|---|
| chr2:77,764,086-77,764,458 | 2,555 | Cwc22 | 5.111774655 | 1.39e-16 |
| chr10:100,376,857-100,377,196 | 541 | Cep290 | 4.558850024 | 1.96e-14 |
| chrX:153,877,445-153,878,004 | -783 | Magea2 | 4.284666292 | 6.25e-15 |
| chrX:168,106,164-168,106,364 | 2,546 | Gm15246 | 4.284289659 | 1.45e-06 |
| chr6:128,951,793-128,952,230 | 0 | Gm15987 | 4.144652989 | 7.97e-12 |
| chr2:77,738,208-77,738,856 | 0 | Cwc22 | 3.903998061 | 3.30e-12 |
| chr2:77,769,683-77,770,655 | -2,612 | Cwc22 | 3.847840485 | 2.62e-03 |
| chr8:15,046,895-15,047,128 | 759 | Arhgef10 | 3.727616071 | 1.11e-04 |
| chr13:99,579,005-99,579,666 | 786 | Map1b | 3.726106396 | 1.50e-24 |
| chr11:68,900,041-68,900,539 | -755 | Pfas | 3.721695241 | 2.70e-06 |
| chr1:38,272,319-38,272,618 | -1,883 | Gm16150 | 3.545813270 | 1.25e-10 |
| chr2:77,739,719-77,739,964 | -1,208 | Cwc22 | 3.464591219 | 4.11e-07 |
| chr2:77,740,651-77,741,016 | -2,140 | Cwc22 | 3.378575713 | 7.86e-07 |
| chr2:77,768,380-77,769,040 | -1,309 | Cwc22 | 3.343382613 | 5.89e-09 |
| chr6:3,762,589-3,763,242 | 388 | Calcr | 3.336800207 | 6.45e-06 |
| chr12:25,445,770-25,446,430 | 1,016 | Gm36723 | 3.322554784 | 1.94e-12 |
| chr19:22,741,057-22,741,450 | -2,548 | Trpm3 | 3.230934092 | 1.32e-04 |
| chr13:69,685,444-69,685,838 | -2,708 | Tent4a | 3.133977061 | 6.67e-21 |
| chr18:30,402,227-30,402,593 | 1,925 | Pik3c3 | 3.130467971 | 2.59e-04 |
| chr10:111,171,776-111,172,554 | -431 | Gm40761 | 3.023841279 | 7.15e-06 |

**Table S6: Top 20 TSS-associated DARs with decreased accessibility in Young TNFα compared to Mature TNFα tenocytes.**

**Table S7:**

**Annotated peak list: top 20 promoter regions with increased accessibility (young TNFα vs young control)**

| Peak | Distance to TSS (bp) | Gene | $Log_2$(fold change) | p-adjust value |
|---|---|---|---|---|
| chr16:36,239,391-36,240,280 | 0 | Gm22263 | 3.440004907 | 1.09e-07 |
| chr19:50,241,479-50,241,987 | -363 | Sorcs1 | 3.306867516 | 9.75e-04 |
| chr12:8,780,701-8,781,100 | 1,554 | Pum2 | 3.236887090 | 6.54e-13 |
| chr14:26,076,362-26,076,730 | -999 | 4933413J09Rik | 3.229871194 | 9.00e-05 |
| chr14:103,072,867-103,073,305 | -673 | Gm34589 | 3.203496000 | 6.40e-05 |
| chr5:105,289,162-105,289,735 | -1,710 | Gbp4 | 3.182405251 | 1.41e-06 |
| chr17:37,794,365-37,794,750 | 0 | H2-M2 | 3.157941062 | 5.53e-05 |
| chr18:39,049,610-39,050,051 | 1,773 | Fgf1 | 3.092135492 | 5.06e-04 |
| chr2:31,335,907-31,336,474 | -273 | Hmcn2 | 3.041895003 | 1.54e-15 |
| chr13:74,081,671-74,081,974 | 2,342 | Trip13 | 3.035925079 | 5.68e-03 |
| chr17:15,724,407-15,724,706 | -139 | Tbp | 3.013012234 | 6.25e-07 |
| chr4:118,005,811-118,007,082 | -1,855 | Kdm4a | 2.982735157 | 3.16e-23 |
| chr11:65,627,959-65,628,451 | -2,301 | Mir744 | 2.973328142 | 6.07e-04 |
| chr18:12,773,486-12,774,198 | -2,203 | Ttc39c | 2.961050656 | 1.50e-28 |
| chr2:34,852,938-34,853,562 | -1,154 | Traf1 | 2.924820891 | 4.73e-13 |
| chr3:132,609,702-132,610,188 | 1,932 | Npnt | 2.903983965 | 7.51e-07 |
| chr2:31,867,120-31,867,621 | -2,163 | Nup214 | 2.886530541 | 6.07e-16 |
| chr12:8,757,684-8,758,330 | -1,896 | Pum2 | 2.868819147 | 1.66e-06 |
| chr14:118,495,780-118,496,330 | 0 | Mir6241 | 2.860291274 | 3.01e-07 |
| chr6:65,566,984-65,567,706 | 0 | Tnip3 | 2.808571637 | 2.56e-07 |

**Table S7: Top 20 TSS-associated DARs with increased accessibility in Young TNFα compared to Young Control tenocytes.**

**Table S8:**

**Annotated Peak list: top 20 promoter regions with decreased accessibility (young TNFα vs young control)**

| Peak | Distance to TSS (bp) | Gene | Log$_2$(fold change) | p-adjust value |
|---|---|---|---|---|
| chr6:97,333,653-97,333,855 | -2,515 | Frmd4b | 2.195486228 | 2.60e-02 |
| chr11:100,179,782-100,180,134 | 1,978 | Gm14206 | 1.971537842 | 4.66e-04 |
| chr13:44,888,768-44,889,085 | 1,252 | Jarid2 | 1.926935798 | 1.11e-02 |
| chr8:70,823,038-70,823,544 | -2,664 | Comp | 1.838897384 | 3.07e-07 |
| chr4:155,331,044-155,331,441 | -2,818 | Faap20 | 1.775874459 | 1.22e-05 |
| chr7:142,569,539-142,570,363 | -783 | Tspan32 | 1.657202327 | 3.99e-08 |
| chr11:94,837,712-94,838,103 | 1,518 | Col1a1 | 1.591206362 | 9.68e-03 |
| chr6:72,334,201-72,334,550 | 992 | Tmem150a | 1.586906865 | 1.76e-03 |
| chr13:112,614,282-112,614,651 | 2,702 | Il6st | 1.569733678 | 4.54e-02 |
| chr15:78,741,234-78,741,577 | -1,505 | Lgals2 | 1.567730824 | 4.48e-04 |
| chr11:121,442,991-121,443,436 | 2,054 | Tbcd | 1.559693906 | 6.50e-05 |
| chr14:47,604,373-47,604,639 | -569 | Lgals3 | 1.528550007 | 9.41e-05 |
| chr8:71,301,967-71,302,631 | 752 | Kcnn1 | 1.51661510 | 2.35e-02 |
| chr8:27,498,265-27,499,352 | 0 | Gm32389 | 1.516133959 | 3.01e-04 |
| chr16:10,791,659-10,792,078 | 1,365 | Litaf | 1.496714837 | 3.15e-02 |
| chr11:98,238,170-98,238,538 | -692 | Ppp1r1b | 1.479716652 | 6.03e-03 |
| chr9:78,222,625-78,223,106 | -547 | Omt2a | 1.479551778 | 2.18e-03 |
| chr16:91,902,392-91,903,015 | -134 | Mrps6 | 1.478481376 | 9.01e-03 |
| chr3:101,484,760-101,485,082 | 0 | Atp1a1 | 1.474139303 | 1.27e-02 |
| chr13:23,897,452-23,898,081 | -2,615 | Hfe | 1.467762922 | 3.44e-05 |

**Table S8: Top 20 TSS-associated DARs with decreased accessibility in Young TNFα compared to Young Control tenocytes.**

**Table S9:**

**Annotated peak list: top 20 promoter regions with increased accessibility (mature TNFα vs mature control)**

| Peak | Distance to TSS (bp) | Gene | Log$_2$(fold change) | p-adjust value |
|---|---|---|---|---|
| chr11:83,421,142-83,421,696 | 0 | Ccl5 | 3.471247244 | 1.25e-34 |
| chr8:41,376,065-41,376,302 | -2,947 | Pdgfrl | 3.377859002 | 1.51e-05 |
| chr16:36,239,391-36,240,280 | 0 | Gm22263 | 3.138830087 | 7.13e-15 |
| chr7:133,559,510-133,560,070 | 2,924 | Adam12 | 3.087903367 | 4.56e-19 |
| chr2:164,784,133-164,785,059 | 1,433 | Mmp9 | 3.086927175 | 1.82e-17 |
| chr18:12,773,486-12,774,198 | -2,203 | Ttc39c | 3.083496396 | 5.48e-36 |
| chrX:95,137,647-95,137,956 | -1,692 | Msn | 3.059099955 | 9.93e-15 |
| chr11:121,494,847-121,495,421 | 2,316 | Tbcd | 2.982042885 | 5.95e-20 |
| chr2:24,524,975-24,525,832 | 1,659 | Cacna1b | 2.926880912 | 3.73e-23 |
| chr11:55,073,569-55,074,252 | 1,651 | Slc36a2 | 2.912488815 | 4.55e-15 |
| chr2:129,153,365-129,153,952 | -1,473 | Il1a | 2.909334132 | 1.45e-29 |
| chr6:124,884,795-124,885,056 | 1,958 | Lag3 | 2.906827179 | 6.92e-05 |
| chrX:149,118,023-149,118,458 | -1,962 | Gm15104 | 2.872093655 | 1.55e-06 |
| chr2:92,210,240-92,210,489 | -2,774 | Frey1 | 2.840179588 | 8.14e-08 |
| chr4:118,005,811-118,007,082 | -1,855 | Kdm4a | 2.804751368 | 3.72e-22 |
| chrX:95,138,253-95,138,525 | -1,123 | Msn | 2.793415649 | 8.62e-07 |
| chr13:74,081,671-74,081,974 | 2,342 | Trip13 | 2.788508030 | 4.63e-08 |
| chr14:103,072,867-103,073,305 | -673 | Gm34589 | 2.771231668 | 1.50e-08 |
| chr1:87,983,132-87,983,637 | 0 | Ugt1a10 | 2.754495222 | 1.99e-21 |
| chr16:84,509,742-84,510,423 | 0 | Mir155hg | 2.721358656 | 7.67e-24 |

**Table S9: Top 20 TSS-associated DARs with increased accessibility in Mature TNFα compared to Mature Control tenocytes.**

**Table S10:**

**Annotated peak list: top 20 promoter regions with decreased accessibility (mature TNFα vs mature control)**

| Peak | Distance to TSS (bp) | Gene | $Log_2$(fold change) | p-adjust value |
|---|---|---|---|---|
| chr1:93,231,696-93,232,285 | 1,300 | Mterf4 | 2.730194575 | 5.20e-11 |
| chr1:93,230,885-93,231,348 | 2,237 | Mterf4 | 2.595531039 | 1.09e-08 |
| chr11:94,838,555-94,838,801 | 2,361 | Col1a1 | 2.350883262 | 1.74e-04 |
| chr11:94,834,638-94,835,084 | -1,110 | Col1a1 | 2.098568105 | 3.01e-06 |
| chr11:94,838,255-94,838,480 | 2,061 | Col1a1 | 1.963268395 | 6.98e-04 |
| chr13:44,888,768-44,889,085 | 1,252 | Jarid2 | 1.848522097 | 7.52e-03 |
| chr6:28,850,561-28,850,847 | 1,655 | Snd1 | 1.791063638 | 6.93e-03 |
| chr7:83,975,190-83,975,489 | -2,567 | Gm25790 | 1.782977309 | 1.09e-02 |
| chr3:95,814,976-95,815,294 | -2,973 | Car14 | 1.741379718 | 3.16e-02 |
| chr19:53,002,294-53,002,530 | -259 | Xpnpep1 | 1.689542310 | 1.57e-02 |
| chr13:44,000,352-44,001,112 | 2,916 | Gm33115 | 1.618670018 | 3.18e-04 |
| chr8:70,823,038-70,823,544 | -2,664 | Comp | 1.578019103 | 8.49e-04 |
| chr5:32,201,376-32,201,953 | -1,369 | Babam2 | 1.534068489 | 2.60e-04 |
| chr6:83,544,497-83,544,863 | -158 | Gm21284 | 1.523465421 | 5.76e-04 |
| chr19:12,764,251-12,764,994 | 2,679 | Gm44505 | 1.512292478 | 1.23e-03 |
| chr1:44,434,673-44,435,111 | 2,704 | Gm28893 | 1.478668562 | 3.79e-02 |
| chr4:120,941,751-120,942,064 | -405 | Gm23540 | 1.469201273 | 9.68e-03 |
| chr5:107,979,230-107,979,708 | 1,265 | Ube2d2b | 1.451955403 | 3.56e-02 |
| chr12:71,589,911-71,590,552 | 2,530 | 4930404H11Rik | 1.448079001 | 2.26e-02 |
| chr1:152,722,613-152,723,018 | 1,653 | Smg7 | 1.435695265 | 2.66e-02 |

**Table S10: Top 20 TSS-associated DARs with decreased accessibility in Mature TNFα compared to Mature Control tenocytes.**

**Table S11:**

**Enriched motifs list: top 14 *de novo* motifs with increased accessibility (young TNFα vs young control)**

| Motif consensus | Best match | Match score | $Log_2$(motif enrichment) | $-Log_{10}$(p-value) |
|---|---|---|---|---|
| GGAAATTCCC | Nfkb-p65 | 0.95 | 3.62246 | 2.74e+03 |
| TGASTCAT | JunB | 0.98 | 1.10340 | 7.99e+02 |
| ATGANVTCATHN | c-Jun | 0.96 | 1.50815 | 2.79e+02 |
| GGAAAGTCACTB | Nr2e1 | 0.69 | 0.78309 | 2.08e+02 |
| ATGATGCAAY | Cebpg | 0.94 | 0.71495 | 2.04e+02 |
| AYTTCCTG | Etv2 | 0.94 | 0.56924 | 1.60e+02 |
| TCCGTCAT | Atf1 | 0.81 | 0.38823 | 1.11e+02 |
| ACGATTGCTCAT | Cebpa | 0.64 | 1.45743 | 8.25e+01 |
| AGTCATGATGAC | Ere | 0.70 | 1.21195 | 7.73e+01 |
| TGAGCTGT | Znf341 | 0.69 | 0.47164 | 5.67e+01 |
| ACTCAGCGTGAG | Zic | 0.68 | 1.16840 | 4.99e+01 |
| TGTRGGWGGA | Wt1 | 0.68 | 0.86429 | 4.18e+01 |
| GGAGTCAGCC | Mafb | 0.67 | 0.57197 | 4.00e+01 |
| GGGAAGCC | Znf528 | 0.75 | 0.37974 | 3.17e+01 |

**Table S11: Top 14 differentially accessible transcription factor motifs with increased accessibility in Young TNFα compared to Young Control tenocytes.**

**Table S12:**

**Enriched motifs list: top 10 *de novo* motifs with decreased accessibility (young TNFα vs young control)**

| Motif consensus | Best match | Match score | $Log_2$(motif enrichment) | $-Log_{10}$(p-value) |
|---|---|---|---|---|
| TATGAGTCAT | Fra1 | 0.98 | 1.85951 | 1.39e+02 |
| SGTTGCCAAG | Nfic | 0.89 | 0.77955 | 8.17e+01 |
| GGAATGTT | Tead2 | 0.98 | 0.63262 | 6.30e+01 |
| HKTGTGGTTT | Runx1 | 0.96 | 1.36839 | 5.14e+01 |
| CCYATAAA | Hoxc13 | 0.93 | 0.97705 | 5.05e+01 |
| TGCCTTGGAAAG | Nfatc1 | 0.62 | 0.98620 | 3.56e+01 |
| GGCTATTTTTAG | Mef2a | 0.91 | 0.91286 | 3.39e+01 |
| TGGGAGGCTGCC | Tlx | 0.76 | 0.20730 | 3.27e+01 |
| RCYCCACCCH | Klf4 | 0.83 | 0.63021 | 3.16e+01 |
| GGGCCWGGCC | Znf213 | 0.73 | 0.51452 | 3.01e+01 |

**Table S12: Top 10 differentially accessible transcription factor motifs with decreased accessibility in Young TNFα compared to Young Control tenocytes.**

**Table S13:**

**Enriched motifs list: top 12 *de novo* motifs with increased accessibility (mature TNFα vs mature control)**

| Motif consensus | Best match | Match score | $Log_2$(motif enrichment) | $-Log_{10}$(p-value) |
|---|---|---|---|---|
| GGAATTYCCC | Nfkb-p65 | 0.93 | 2.39850 | 3.39e+03 |
| TGAGTCAB | Fra1 | 0.98 | 1.07887 | 8.87e+02 |
| RTTKCMWMAB | Cebp | 0.87 | 0.52912 | 3.58e+02 |
| DKATGACRTCAT | Creb5 | 0.95 | 0.94553 | 3.27e+02 |
| CTGGAAAGTCAC | Nr2e1 | 0.67 | 0.97589 | 2.33e+02 |
| ACAGGAAATGAC | Erg | 0.91 | 1.06095 | 1.75e+02 |
| AAAACCCC | Rel | 0.67 | 0.62846 | 1.31e+02 |
| GATGAGCAAT | Nfe2 | 0.67 | 0.70754 | 1.31e+02 |
| GGAAWVTYTG | Nfatc4 | 0.70 | 0.78390 | 1.12e+02 |
| AATGACTAGTCA | Smad5 | 0.64 | 0.55184 | 1.05e+02 |
| TGATACAGTCAG | Cphx1 | 0.67 | 0.57087 | 8.69e+01 |
| TCCGKCAT | Hoxa13 | 0.69 | 0.58802 | 3.10e+01 |

**Table S13: Top 12 differentially accessible transcription factor motifs with increased accessibility in Mature TNFα compared to Mature Control tenocytes.**

**Table S14:**

**Enriched motifs list: top 10 *de novo* motifs with decreased accessibility (mature TNFα vs mature control)**

| Motif consensus | Best match | Match score | $Log_2$(motif enrichment) | $-Log_{10}$(p-value) |
|---|---|---|---|---|
| GGAATGYGGH | Tead1 | 0.95 | 1.12608 | 1.09e+02 |
| CYTATAAA | Hoxc13 | 0.96 | 1.14987 | 1.09e+02 |
| ATGAGTCA | Jun | 1.00 | 1.20619 | 8.53e+01 |
| AAACCACA | Runx1 | 0.97 | 1.07303 | 6.24e+01 |
| YCCCTGGRGA | Ebf1 | 0.88 | 0.65063 | 5.08e+01 |
| KSRATTTGGC | Nfix | 0.89 | 1.06529 | 4.86e+01 |
| TTCCAAGCCT | Zfp809 | 0.75 | 0.77754 | 3.52e+01 |
| GAATTCCTATAA | Bcl6b | 0.74 | 0.82409 | 3.24e+01 |
| GTTTAACTAG | Hmbox1 | 0.83 | 2.14002 | 3.23e+01 |
| TTCCTGGCAGAC | Stat3 | 0.74 | 0.66146 | 3.09e+01 |

**Table S14: Top 10 differentially accessible transcription factor motifs with decreased accessibility in Mature TNFα compared to Mature Control tenocytes.**

**Table S15:**

**Gene list: top 20 upregulated genes (young control vs mature control)**

| Gene name | Gene function | Log$_2$(fold change) | p-adjust value |
|---|---|---|---|
| Gm43521 | non-coding RNA | 5.233455719 | 4.06e-03 |
| Gm38340 | Predicted pseudogene | 5.089197741 | 4.04e-02 |
| Lefty1 | TGF-B superfamily ligand | 4.374196008 | 3.39e-02 |
| Tpm3-rs7 | Direct actin filament organization | 4.083044563 | 1.78e-02 |
| Gm11839 | Predicted pseudogene | 4.082091439 | 2.48e-02 |
| Rpsa-ps10 | Structural ribosome component | 4.079079502 | 5.41e-04 |
| C1qtnf3 | Cellular metabolism and negative regulation of NFκB | 3.945209544 | 2.69e-14 |
| Fndc1 | Cellular signaling in response to stress | 3.839540960 | 1.34e-31 |
| Fgf9 | Fibroblast activation and proliferation | 3.795558348 | 2.52e-08 |
| S100a7a | Calcium signaling related to inflammation | 3.776730259 | 1.43e-23 |
| Spink5 | Anti-inflammatory peptidase inhibitor | 3.731409288 | 3.24e-08 |
| Kcnk1 | Potassium ion channel | 3.475495519 | 9.98e-07 |
| Rxfp3 | GCPR for cytokine regulation | 3.468998364 | 8.56e-03 |
| Gm37201 | Predicted pseudogene | 3.443192112 | 4.37e-03 |
| Col2a1 | ECM structural component | 3.391264108 | 5.50e-07 |
| Afap1l1 | Actin integrity modulator and Src binding partner | 3.384185412 | 4.67e-03 |
| Gsta1 | Antioxidation and metabolism | 3.261330272 | 2.95e-02 |
| Gm10704 | RNA regulation and processing | 3.255735732 | 3.10e-03 |
| Gm13577 | long non-coding RNA | 3.250360608 | 5.45e-07 |
| Tnn | Tendon ECM protein | 3.238296408 | 8.28e-03 |

**Table S15: Top 20 differentially expressed genes with increased expression in Young Control compared to Mature Control.**

**Table S16:**

**Gene list: top 20 downregulated genes (young control vs mature control)**

| Gene name | Gene function | $Log_2$(fold change) | p-adjust value |
|---|---|---|---|
| Hist1h2al | Chromatin structural protein | 11.590312990 | 3.45e-02 |
| Gm42047 | MHC class 1-associated lncRNA | 9.925056896 | 3.66e-03 |
| Gm6969 | Predicted pseudogene | 7.150207010 | 2.91e-02 |
| Dmp1 | ECM and adhesive protein | 7.073668553 | 1.38e-04 |
| Zfp979 | Transcriptional regulation | 7.055103329 | 3.94e-04 |
| Ptprv | Tyrosine phosphatase | 6.701480925 | 9.43e-03 |
| Pianp | Neuronal protein signaling | 6.485878972 | 3.35e-13 |
| Gm24598 | snoRNA | 6.181795884 | 7.94e-05 |
| Sprr1a | Peptide crosslinking and skin structural component | 5.587672797 | 4.32e-03 |
| Gm13736 | Predicted pseudogene | 5.319306249 | 2.90e-02 |
| Best1 | Neuronal gated ion channel activation | 5.302356837 | 4.63e-03 |
| Gm10499 | MHC class1 antigen presentation | 4.806033359 | 2.61e-07 |
| Rpl15-ps2 | Ribosome structural protein | 4.718928857 | 1.26e-03 |
| Sncg | Neurological activation | 4.580660108 | 4.25e-04 |
| Srd5a2 | Steroid activation | 4.537643562 | 6.36e-05 |
| Galnt15 | Protein glycosylation | 4.341021623 | 7.20e-06 |
| Rbpj-ps3 | Notch signaling modulator | 4.265197091 | 2.46e-08 |
| Coch | Structural ECM component | 4.219125920 | 1.36e-02 |
| Zbp1 | Innate immune detection of DNA | 4.022051265 | 3.33e-03 |
| Car3 | Oxidative homeostasis | 3.929801591 | 3.21e-02 |

**Table S16: Top 20 differentially expressed genes with decreased expression in Young Control compared to Mature Control.**

1 **Table S17:**

**Gene list: top 20 upregulated genes (young TNFα vs mature TNFα)**

| Gene name | Gene function | $Log_2$(fold change) | p-adjust value |
|---|---|---|---|
| Tnn | Tendon ECM protein | 4.829778722 | 2.12e-05 |
| Gdf10 | TGF-B superfamily ligand | 4.495013591 | 3.09e-03 |
| Cntn2 | Cell adhesion and migration | 4.124983262 | 2.18e-04 |
| C1qtnf3 | Regulation of NFkB signaling and cytokine production | 4.059853162 | 4.48e-05 |
| Tpm3-rs7 | Direct actin filament organization | 4.018777186 | 2.32e-02 |
| Lefty1 | TGF-B superfamily ligand | 4.000425916 | 4.84e-02 |
| Gm13212 | Fatty acid metabolism | 3.961371739 | 5.55e-07 |
| Fndc1 | Cellular signaling in response to stress | 3.935280171 | 4.64e-06 |
| Fermt1 | Integrin and focal adhesion signaling | 3.932906244 | 2.25e-04 |
| Afap1l1 | Actin integrity modulator and Src binding partner | 3.844217264 | 2.37e-03 |
| S100a7a | Calcium signaling related to inflammation | 3.698617377 | 3.79e-26 |
| Rpsa-ps10 | Structural ribosome component | 3.513436767 | 2.54e-02 |
| Spink5 | Anti-inflammatory peptidase inhibitor | 3.501372016 | 1.12e-02 |
| Lilrb4a | Anti-inflammatory receptor | 3.484486697 | 5.92e-03 |
| Gsta4 | Antioxidation and metabolism | 3.416539716 | 4.05e-03 |
| Meox1 | Developmental transcription factor | 3.416311003 | 4.22e-12 |
| Kcnk1 | Potassium ion channel | 3.303470642 | 3.47e-07 |
| Col2a1 | Cartilage ECM protein | 3.285349588 | 8.58e-06 |
| Car6 | Carbonate dehydrogenase | 3.187136665 | 4.20e-02 |
| Ednra | Cell adhesion receptor | 3.112821209 | 1.11e-02 |

**Table S17: Top 20 differentially expressed genes with increased expression in Young TNFα compared to Mature TNFα.**

**Table S18:**

**Gene list: top 20 downregulated genes (young TNFα vs mature TNFα)**

| Gene name | Gene function | $Log_2$(fold change) | p-adjust value |
|---|---|---|---|
| Hist1h2al | Chromatin structural protein | 11.57615849 | 2.61e-02 |
| Gm42047 | MHC class 1-associated lncRNA | 11.29805008 | 6.63e-03 |
| Rpl15-ps2 | Ribosomal structural protein | 8.097734760 | 3.10e-02 |
| Zfp979 | Transcriptional regulation | 7.020823322 | 3.36e-04 |
| Gm24598 | snoRNA | 6.519295747 | 2.29e-04 |
| Gm6969 | Predicted pseudogene | 5.928960952 | 2.83e-03 |
| Ptprv | Tyrosine phosphatase | 5.842419451 | 2.40e-02 |
| Htra4 | Chaperone protease | 5.688662278 | 4.73e-03 |
| Srd5a2 | Steroid metabolism | 5.412678906 | 1.53e-05 |
| Gm13736 | Predicted pseudogene | 5.109780477 | 1.17e-02 |
| Mcpt8 | Basophil endopeptidase | 4.481033215 | 2.31e-08 |
| Sncg | Neurological regulation | 4.467024175 | 9.26e-05 |
| Coch | Structural ECM protein | 4.409361334 | 6.18e-03 |
| Zfp984 | Transcriptional regulation | 4.154170116 | 4.01e-06 |
| Gm13337 | Predicted pseudogene | 4.118710173 | 4.52e-02 |
| Hmgb1-ps7 | Predicted pseudogene | 4.053980885 | 1.40e-02 |
| Car3 | Oxidative homeostasis | 3.891693685 | 3.01e-03 |
| Esm1 | Regulation of angiogenesis, inflammation, and vascular permeability | 3.824885484 | 4.10e-09 |
| Il33 | Pro-inflammatory cytokine | 3.792829181 | 6.66e-20 |
| Piwil2 | Germline development | 3.789915459 | 2.78e-13 |

**Table S18: Top 20 differentially expressed genes with decreased expression in Young TNFα compared to Mature TNFα.**

**Table S19:**

**Gene list: top 20 upregulated genes (young TNFα vs young Control)**

| Gene name | Gene function | $Log_2$(fold change) | p-adjust value |
|---|---|---|---|
| Gpr84 | GPCR in inflammatory signaling | 10.213456000 | 2.03e-14 |
| Pglyrp2 | Innate immune surveillance | 10.055453130 | 1.13e-06 |
| Nos2 | Nitric oxide production | 9.823295234 | 2.34e-05 |
| Cxcl2 | Chemokine | 9.638945349 | 3.02e-24 |
| Gm16685 | non-coding RNA | 9.005654173 | 4.94e-16 |
| Acod1 | Innate immune response | 8.419792857 | 4.57e-07 |
| Gbp5 | Inflammasome activator | 8.371451033 | 2.17e-04 |
| Olr1 | Endocytosis receptor | 7.866714066 | 5.43e-17 |
| Ccl5 | Chemokine | 7.837928914 | 3.94e-84 |
| Ccl20 | Chemokine | 7.571017598 | 9.63e-16 |
| Cyp4f39 | Ceramide biosynthesis | 7.521793893 | 4.31e-11 |
| Neurl3 | Anti-viral response | 7.422309303 | 1.62e-09 |
| Mmp9 | ECM degredation | 7.264247161 | 1.67e-110 |
| Nepn | TGF-B signaling | 7.184381214 | 3.55e-05 |
| Sprr1a | Nerve regeneration | 7.132651321 | 4.75e-06 |
| Gpr132 | GPCR in neuronal signaling | 7.080346187 | 6.23e-05 |
| Cxcl10 | Chemokine | 7.062319196 | 2.64e-29 |
| Gm29371 | long non-coding RNA | 6.977715631 | 9.19e-05 |
| Cd83 | T cell immune regulation | 6.894328824 | 1.25e-04 |
| Traf1 | TNFα-signaling pathway | 6.826004214 | 1.72e-27 |

**Table S19: Top 20 differentially expressed genes with increased expression in Young TNFα compared to Young Control.**

**Table S20:**

**Gene list: top 20 downregulated genes (young TNFα vs young control)**

| Gene name | Gene function | Log$_2$(fold change) | p-adjust value |
|---|---|---|---|
| Gm37333 | long non-coding RNA | 4.092636786 | 2.61e-02 |
| Arg1 | Arginase activity | 3.573196312 | 5.92e-06 |
| Hpgd | Inactivation of prostaglandin signaling | 3.246583945 | 4.73e-13 |
| Wnt4 | Promote Wnt signaling | 2.999173390 | 7.25e-07 |
| Dhrs3 | Retinoic acid receptor signaling | 2.691241318 | 8.25e-27 |
| Rasl11a | Regulation of RNA polymerase 1 and G protein activity | 2.679054980 | 6.57e-22 |
| Cdh3 | Intercellular adhesion | 2.582034069 | 3.09e-02 |
| Gpr88 | GPCR in cytoskeletal motor activity | 2.580099925 | 7.57e-05 |
| Hey2 | Cardiovascular transcription factor | 2.552368141 | 2.63e-02 |
| Arsi | Sulfatase activity | 2.411818770 | 1.81e-07 |
| Spon2 | Immunomodulatory ECM protein | 2.347631111 | 4.39e-18 |
| Aqp5 | Water transporter | 2.244084996 | 5.40e-19 |
| Npy | Neurotransmitter | 2.242566307 | 5.96e-14 |
| Fos | Mesenchymal transcription factor | 2.236613432 | 4.86e-07 |
| Foxq1 | Cell survival ranscriptional factor | 2.192482944 | 1.59e-02 |
| Dio3 | Thyroid hormone regulator | 2.178858111 | 1.55e-20 |
| Crabp2 | Retinoic acid signaling modulation | 2.078209424 | 8.91e-12 |
| Adcy5 | Production of cAMP | 2.048849806 | 4.52e-02 |
| Phyhipl | Nervous system homeostasis | 2.041744373 | 2.96e-05 |
| Wnt16 | Regulation of bone homeostasis | 1.927363946 | 6.95e-07 |

**Table S20: Top 20 differentially expressed genes with decreased expression in Young TNFα compared to Young Control.**

**Table S21:**

**Gene list: top 20 upregulated genes (mature TNFα vs mature control)**

| Gene name | Gene function | $Log_2$(fold change) | p-adjust value |
|---|---|---|---|
| Nos2 | Nitric oxide production | 11.407642270 | 5.49e-08 |
| Acod1 | Innate immune response | 10.996960430 | 6.82e-06 |
| Pglyrp2 | Innate immune surveillance | 10.909468310 | 1.20e-06 |
| Ccl5 | Chemokine | 10.676376050 | 2.16e-15 |
| Cxcl2 | Chemokine | 10.537634370 | 3.61e-14 |
| Gpr84 | GPCR in inflammatory signaling | 9.861863617 | 4.91e-09 |
| Ccl20 | Chemokine | 9.226885177 | 1.42e-09 |
| Cxcl10 | Chemokine | 8.704209654 | 4.33e-29 |
| Traf1 | TNFα-signaling pathway | 7.997024867 | 1.27e-07 |
| Heatr9 | Induction of cytokine production | 7.948656062 | 4.28e-08 |
| Nlrp3 | Inflammasome formation | 7.934333277 | 3.98e-08 |
| Mmp9 | ECM degredation | 7.858990498 | 1.66e-22 |
| Gm11419 | non-coding RNA | 7.782013752 | 1.12e-07 |
| Cxcl9 | Chemokine | 7.595370840 | 4.55e-07 |
| Csf2 | Macrophage cytokine | 7.503024646 | 2.57e-03 |
| Mmp1b | ECM degredation | 7.477470877 | 4.46e-05 |
| Cyp4f39 | Ceramide biosynthesis | 7.241472919 | 2.19e-04 |
| Clec4e | Innate immunity pattern recognition | 7.026876861 | 1.72e-07 |
| Mx1 | Inhibition of viral replication | 6.935322193 | 2.44e-05 |
| Gm16685 | non-coding RNA | 6.902468105 | 1.07e-42 |

**Table S21: Top 20 differentially expressed genes with increased expression in Mature TNFα compared to Mature Control.**

**Table S22:**

**Gene list: top 20 downregulated genes (mature TNFα vs mature control)**

| Gene name | Gene function | Log$_2$(fold change) | p-adjust value |
|---|---|---|---|
| C2cd4a | Acute inflammation and vessel permeability | 3.601779456 | 3.99e-03 |
| Atp12a | Cation transporter | 3.578465313 | 1.70e-03 |
| Arg1 | Arginase activity | 3.492059369 | 1.05e-03 |
| Lgr5 | GPCR in Wnt signaling | 3.446485253 | 3.15e-02 |
| Scn1a | Sodium ion transporter | 3.273843401 | 2.21e-03 |
| Megf10 | Regulation of cell adhesion and motility | 3.192442704 | 1.61e-03 |
| Krt13 | Epithelial cell structural integrity | 3.094365299 | 2.61e-03 |
| Hpgd | Inactivation of prostaglandin signaling | 3.043643734 | 2.09e-19 |
| Arsi | Sulfatase activity | 2.998337282 | 7.60e-07 |
| 8430436N08Rik | non-coding RNA | 2.981125077 | 4.78e-02 |
| Aldh1a7 | Aldehyde metabolism | 2.933551949 | 3.82e-02 |
| Slc4a8 | Sodium ion transporter | 2.907235686 | 4.52e-03 |
| Dhrs3 | Retinoic acid receptor signaling | 2.901126399 | 1.06e-100 |
| Gpr88 | GPCR in cytoskeletal motor activity | 2.794727091 | 1.23e-13 |
| Ppl | Desmosome formation | 2.563400420 | 6.42e-18 |
| Gpd1 | Lipid metabolism | 2.536095751 | 6.41e-08 |
| Aqp5 | Water transporter | 2.535815526 | 3.99e-44 |
| Rab3il1 | Synaptic GEF | 2.530332388 | 7.93e-08 |
| Arc | mRNA regulation in synapses | 2.477742205 | 1.41e-09 |
| Kcnh2 | Potassium ion channel | 2.454592028 | 5.89e-05 |

**Table S22: Top 20 differentially expressed genes with decreased expression in Mature TNFα compared to Mature Control.**

**Table S23:**

**Gene list: top 19 genes with interaction effects (treatment and age)**

| Gene name | Gene function | $Log_2$(fold change) | p-adjust value |
|---|---|---|---|
| Ccl9 | Chemokine | 3.227509413 | 1.05e-05 |
| Saa3 | Induction of cytokine production | 2.973120575 | 3.47e-01 |
| Mmp12 | ECM degredation | 2.858383860 | 8.81e-02 |
| Cilp2 | ECM structural component | 2.742380451 | 4.36e-03 |
| Cxcl3 | Chemokine | 2.623943465 | 3.24e-03 |
| Gpm6b | Regulation of bone formation | 2.400963377 | 1.40e-01 |
| Cxcl1 | Chemokine | 1.867442002 | 1.27e-01 |
| Cxcl5 | Chemokine | 1.784851722 | 5.73e-02 |
| Slco4a1 | Organic transporter in migration signaling | 1.701481060 | 6.60e-03 |
| Itga11 | Cell-ECM binding and migration | 1.439481459 | 9.64e-02 |
| Osgin1 | Oxidative stress response in microtuble dynamics | 1.425242292 | 4.77e-02 |
| Ccl7 | Chemokine | 1.298327350 | 6.23e-03 |
| Rnf39 | MHC and immune synapse signaling | 1.281416568 | 1.40e-01 |
| Ptx3 | Fibroblast activation in innate immunity | 1.254223652 | 1.65e-01 |
| Bmper | BMP-2,4 signaling inhibitor | 1.068449804 | 2.20e-03 |
| Lce1g | Epidermal differentiation | 1.034748113 | 7.47e-02 |
| Chst15 | GAG synthesis and excretion | -1.282008378 | 8.81e-02 |
| Plekhg1 | Nuclear RhoGEF | -1.865861014 | 9.64e-02 |
| Igfbp5 | Insulin-like GF signaling | -2.398125475 | 6.05e-02 |

**Table S23: Top 19 genes identified with two-way interaction analysis (Age x Treatment) with p-adjust value < 0.4.** Positive $Log_2$(fold change) values indicate a more positive-valued response in mature cells, and negative $Log_2$(fold change) values indicate a more positive-valued response in young cells.

**Table S24:**

**RT-PCR primer list**

| Gene | | Sequence (5'-3') |
|---|---|---|
| NFκB | Forward | GCTGCCAAAGAAGGACACGACA |
| | Reverse | GGCAGGCTATTGCTCATCACAG |
| Mmp13 | Forward | GGTCCCAAACGAACTTAACTTACA |
| | Reverse | CCTTGAACGTCATCAGGAAGC |
| Caspase-11 | Forward | ACAAACACCCTGACA AACCAC |
| | Reverse | CACTGCGTTCAGCATTGTTAAA |

**Table S24: List of RT-PCR primers used in this study.**
